# Evolutionary analysis in *Enterobacterales* of the Rcs-repressor protein IgaA unveils two cytoplasmic small β-barrel domains central for function

**DOI:** 10.1101/2022.08.18.504389

**Authors:** Leticia Rodríguez, Marcos Peñalver, Patricia Casino, Francisco García-del Portillo

## Abstract

The Rcs sensor system, comprised by the proteins RcsB/RcsC/RcsD and RcsF, is used by bacteria of the order *Enterobacterales* to withstand envelope damage. Under non-stress conditions, the system is repressed by the membrane protein IgaA. How IgaA has evolved within *Enterobacterales* in concert with the Rcs system has not been explored. Here, we report phylogenetic data supporting co-evolution of IgaA with the inner membrane proteins RcsC and RcsD. Functional assays showed that IgaA from representative genera as *Shigella* and *Dickeya*, but not those from *Yersinia* or the endosymbionts *Photorhabdus* and *Sodalis*, repress the Rcs system when expressed in a heterogenous host like *Salmonella enterica* serovar Typhimurium. IgaA structural features have therefore diverged among *Enterobacterales*. Modelling of IgaA structure unveiled one periplasmic and two cytoplasmic β-rich architectures forming partially-closed small β-barrel (SBB) domains related to OB (oligonucleotide/oligosaccharide binding motif) fold domains. Interactions among conserved residues were mapped in a connector linking SBB-1 domain of cytoplasmic region cyt1 to SBB-2 domain of region cyt2 (residues E180-R265); the C-terminus of cyt1 facing cyt2 (R188-E194-D309 and T191-H326); and, between cyt2-cyt3 regions (H293-E328-R686). These interactions identify a previously unnoticed "hybrid" SBB-2 domain. We also identified interactions absent in the IgaA variants not functional in *S.* Typhimurium, including H192-P249, which links cyt1 to cyt2, R255-D313 and D287-R314. A short α-helix (α6) located in the SSB-1 domain is also missing in the non-complementing IgaA tested. Taken together, our data support a central role of the two cytoplasmic SBB domains in IgaA function and evolution.

**SIGNIFICANCE:** The "intracellular growth attenuator A" protein (IgaA) was first reported as repressor of the Rcs system in *S. enterica* serovar Typhimurium. IgaA orthologs were later studied in other genera and families of the *Enterobacterales* order, mainly in *Escherichia coli*. Despite intense investigation about the mechanism by which IgaA controls the Rcs system, the extent at which IgaA evolved within families of the *Enterobacterales* order has not been investigated. Using a combination of functional assays and *in silico* structural analyses, our work provides a detail map of conserved and divergent residues in IgaA representing interactions occurring in all *Enterobacterales* and others that may have diverged concomitantly to interacting proteins, probably for responding to specific environments. Future studies involving mutagenesis of these residues in IgaA of *Enterobacterales* families and genera of interest will certainly provide valuable insights into the regulation acting in the IgaA-Rcs axis.

## INTRODUCTION

The Rcs system plays a key sensory role in bacteria of the order *Enterobacterales* for combating insults to their envelope.^1,2^ Upon signal perception, the Rcs system is activated by a phosphorelay cascade involving the inner membrane proteins RcsC/RcsD, which transduce the signal to the cytoplasmic transcriptional regulator RcsB.^1^ Studies performed mostly in *Salmonella enterica* serovar Typhimurium (*S*. Typhimurium) and *Escherichia coli* have provided compelling evidence for a tight control of the Rcs system under non-stress conditions. This control impedes unnecessary activation, which has been shown to be detrimental for viability.^3–7^

An important element in the control of the Rcs system is the negative regulator IgaA, an essential inner membrane protein identified in *S*. Typhimurium as a factor that reduces the growth rate of this pathogen following the entry into eukaryotic cells.^3,8^ Previously, an IgaA ortholog of *Proteus mirabilis* named UmoB, was shown to control flagellar synthesis and swarming^9^, functions controlled by the Rcs system. IgaA has five transmembrane domains with three defined cytoplasmic regions and a large periplasmic loop. The mechanism by which IgaA represses Rcs signaling is under intense investigation. An interaction between the outer membrane protein RcsF and the periplasmic region of IgaA was proposed to relieve its repression over RcsC/RcsD.^10^ More recent studies have reported interactions between the outer membrane protein OmpA and RcsF, modulating the RcsF-IgaA interaction,^11^ and, between RcsD and IgaA,^6,7^ this latter involving periplasmic and cytoplasmic domains.^7^

Besides *E. coli* and *S*. Typhimurium, IgaA orthologs have been studied in other *Enterobacterales* genera, mainly in relation to virulence given the control that this protein exerts over the Rcs system, required for expression of flagella and capsules among other virulence-related factors.^1,2^ Ortholog members of the IgaA family characterized to date include UmoB of *Proteus miriabilis*^9^ and GumB of *Serratia marcenses*.^12^ In contrast to *S*. Typhimurium and *E. coli*,^3,5–7^ the lack of the IgaA orthologs UmoB and GumB does not compromise viability in *P. mirabilis* or *S. marcenses*.^9,12^ These findings indicate that, although conserving its master role as repressor of the Rcs system, IgaA may have evolved differently within the *Enterobacterales* order. This variability probably relies on factors related to the lifestyle or the environmental niche that is occupied by the bacterium, resulting in different signals perceived by the IgaA-Rcs axis. Consistent with this hypothesis, studies performed in *S. marcenses* demonstrated that a null *gumB* mutation could be complemented by expressing IgaA of *S*. Typhimurium or *E.coli* whereas expression of a predicted IgaA ortholog of *Klebsiella pneumoniae*, KumO, restored wild type colony morphology but not pigmentation.^12^ Differences in the capacity of RcsC from pathogenic *E. coli*, *S. enterica* or *Yersinia pestis* to complement an *rcsC* mutation in *E. coli* K-12 were also reported^13^.

In this study, we performed a phylogenetic analysis of IgaA and Rcs proteins in *Enterobacterales* genera with genome information available in databases. This was followed by expression of IgaA from several *Enterobacterales* genera in *S*. Typhimurium to interrogate via a lethality assay, for their capacity to replace endogenous IgaA in its role as Rcs repressor. The complementation, obtained for only a few of the heterologous IgaA expressed, support divergence in the evolution of IgaA within the *Enterobacterales* order. Modelling and structural analyses of the distinct IgaA tested allowed us to map conserved and varied positions relevant for Rcs repression and to assign a functional role to a previously unnoticed partially-closed small β-barrel (SBB) domain^14^ related to OB (oligonucleotide/oligosaccharide binding motif) fold domain.^15^ One of these SBB domains is formed by residues of the two major cytoplasmic regions of IgaA, cyt1 and cyt2.

## RESULTS

### Phylogenetic analysis of IgaA and Rcs proteins within the *Enterobacterales* order

Although IgaA orthologs have been characterized in some *Enterobacterales* genera such as *Salmonella, Escherichia, Proteus* and *Serratia*, to our knowledge no systematic analysis of the distribution and evolution of the membrane components of the Rcs system and its repressor IgaA has been performed. Using as source the "Bacterial and Viral Bioinformatics Resource Center" (BV-BRC) (https://www.bv-brc.org/), we identified IgaA, RcsC, RcsD and RcsF orthologs by BLASTp in most of the *Enterobacterales* genera, with the exception of the obligate endosymbionts *Buchnera aphidicola* and *Wigglesworthia glossinidia,* which lack the entire set of proteins (Tables S1-S2). Interestingly, other bacteria show loss or truncation of specific components. Thus, all isolates of *Y. pestis* with genome sequenced encode a truncated RcsD as well as some *S. enterica* subp. *arizonae* isolates (Table S1). Truncation of RcsD in *Y. pestis* was already inferred in an early study^16^. Other interesting cases are *Budvicia aquatica*, which lacks RcsB, RcsC and RcsD but encodes a full-length IgaA and RcsF; *Pleisomonas shigelloides*, which lacks IgaA, RcsB, and RcsD, but encodes a full-length RcsF and a truncated RcsC; and the insect symbiont *Arsnophonus nasoniae*, which a genome encoding a truncated RcsF (Table S1). We next performed a phylogenetic analysis of the IgaA/Rcs proteins encoded by 118 genomes deposited in the BV-BRC database covering all known *Enterobacterales* genera (Table S1). This analysis revealed more similarity in the phylogenetic trees obtained for RcsC and RcsD in comparison to those of IgaA, RcsF and RcsB (Figure S1). This finding suggests that the interaction in the phosphorelay known for RcsC and RcsD could have been the origin of the IgaA/Rcs regulatory axis. With this information, we reanalysed the phylogenetic data of IgaA and Rcs proteins with 40 species from a reduced number of *Enterobacterales* genera representing different lifestyles and occupying distinct ecological niches: *Shigella, Dickeya*, *Yersinia*, *Photorhabdus* and *Sodalis* (Table S3). These genera are classified in different families of the *Enterobacteriales* order: *Shigella* (*Enterobacteriaceae*); *Dickeya* (*Pectobacteriaceae*); *Photorhadbus* (*Morganellaceae*); and *Sodalis* (*Bruguierivoracaceae*).^17^ Importantly, the phylogenetic tree for IgaA of these representative genera, including species of the genus *Escherichia* (to which *Shigella* and *Salmonella* are close phylogenetically), showed a marked overlap with those of RcsC and RcsD (Figure 1). This feature was however not observed for RcsF, RcsB or an unrelated protein as the chaperone DnaK (Figure 1). Taken together, these phylogenetic analyses indicated that in bacteria of the *Enterobacterales* order IgaA could have integrated into the Rcs regulatory network primarily by interacting with RcsC/RcsD.

**Figure 1.**
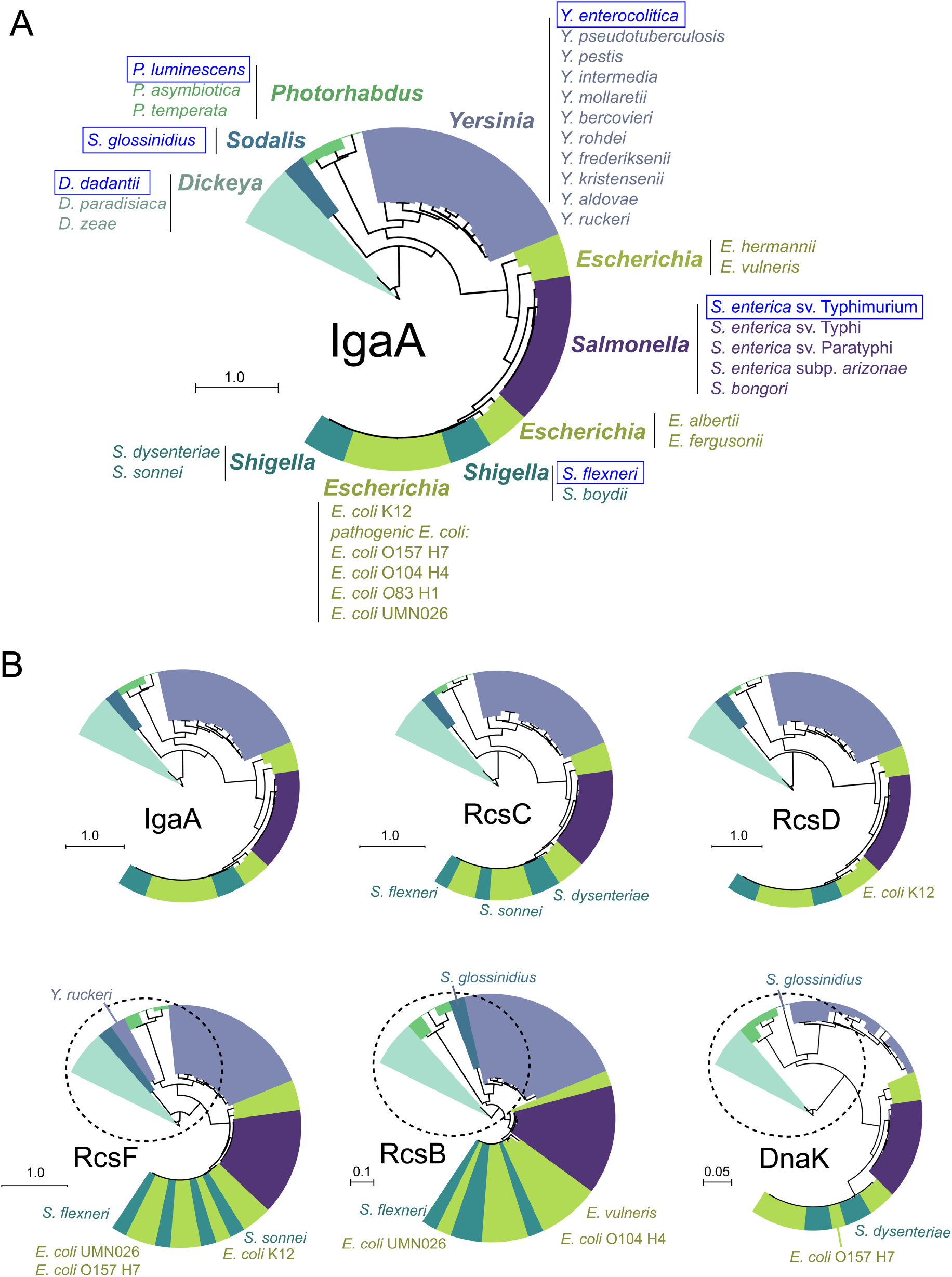
Phylogeny of IgaA and proteins of the Rcs system in selected genera of the *Enterobacterales* order. (A) Phylogenetic tree obtained for IgaA after analyzing a total of 38 representative species within families with different lifestyles belonging to the *Enterobacterales* order: *Salmonella, Escherichia* and *Shigella* (family *Enterobacteriaceae*)*; Dickeya* (*Pectobacteriaceae*); *Yersinia* (*Yersiniaceae*), *Photorhadbus* (*Morganellaceae*); and *Sodalis* (*Bruguierivoracaceae*); (B) Phylogenetic trees obtained from representative species for IgaA, RcsC, RcsD, RcsB, RcsF and DnaK, this latter unrelated protein chosen as control. Protein trees for each set were built using the BV-BRC Gen Tree tool with no end trimming or gappy sequences removal and using RAxML algorithm with LG evolutionary model. Trees were visualized using iTOL version 6.5.7. Bars indicate phylogenetic distance.

### Expression of heterologous IgaA proteins in *S*. Typhimurium to repress the Rcs system

Based on the phylogenetic data (Figure 1), we cloned orthologs of representative species to test their capacity to replace endogenous IgaA and, therefore, to repress the *S*. Typhimurium Rcs system. To this aim, the *igaA* gene was cloned from the following bacterial genera and species: the plant pathogen *D. dadantii* (formerly *Erwinia chrysanthemi*), the animal pathogens *S. flexneri* and *Y. enterocolitica,* and the endosymbionts *S. glossinidius* and *P. luminescens*. The rationale for testing function of IgaA from *S. glossinidius* (tsetse fly *-Glossina* spp.-endosymbiont) and *P. luminescens* (nematode endosymbiont) was the known loss of this protein together with the entire set of Rcs proteins in obligate endosymbionts like *B. aphidicola* or *W. glossinidia* (see Table S1). These observations suggested that IgaA function might become dispensable once bacteria adapt to the invariant and stable intracellular niche of host cells. The IgaA variants were engineered with a Myc-tag in the C-terminus to monitor their production and were expressed from pBAD24, a plasmid bearing a promoter that is repressed with glucose and inducible with arabinose. Immunoassays with anti-Myc antibody confirmed the expression of these heterologous IgaA variants expressed in *S*. Typhimurium strain SL1344 (Figure 2A). The lack of mutations in the respective sequences of these heterologous IgaA variants was confirmed by sequencing.

**Figure 2.**
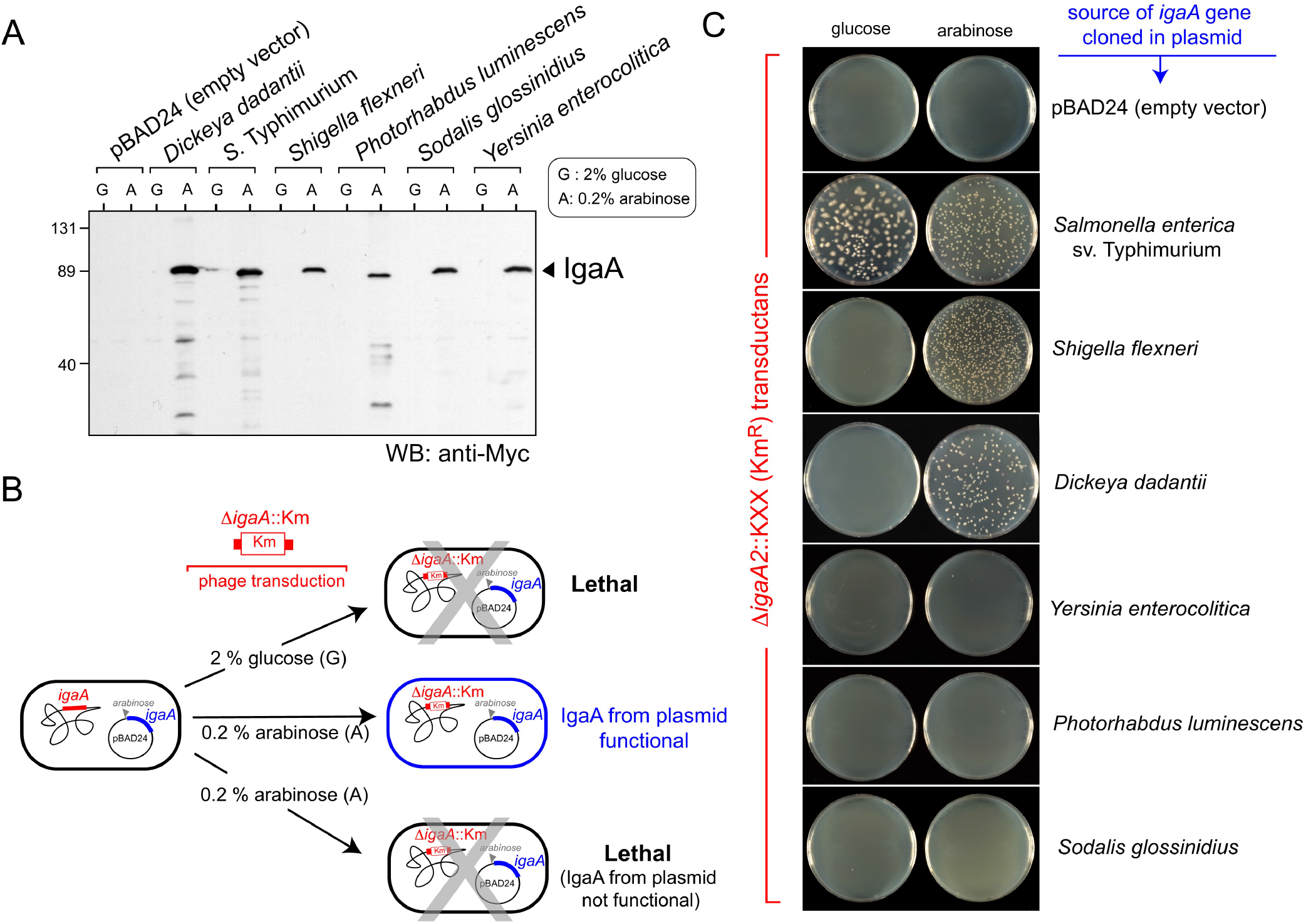
Functional assay of IgaA from several *Enterobacterales* genera produced ectopically in *S*. Typhimurium. (A) *igaA* wild type alleles from the indicated *Enterobacterales* genera and species were cloned in pBAD24 plasmid, a vector bearing an arabinose-induced promoter to drive expression of the insert. These *igaA* alleles were cloned as tagged version with a Myc epitope located at the 3’ end. Shown are total extracts of the indicated *S*. Typhimurium strains grown either in the presence of glucose, a repressor of the pBAD promoter, or arabinose as inducer. Numbers in the left side refer to the molecular weight markers (in kDa); (B) Scheme of the experimental design used to test function of IgaA from distinct *Enterobacterales* genera in *S*. Typhimurium based on suppression of lethality due to the absence of endogenous IgaA; (C) Result of the phage transduction of an *igaA2*::KXX allele in the presence of glucose or arabinose. Note that transductants were obtained only when expressing IgaA from *S*. Typhimurium, and the genera *Shigella* and *Dickeya* (*Enterobacteriaceae* and *Pectobacteriaceae* families).

Next, we took advantage of the essentiality of IgaA in *S*. Typhimurium to test whether the ectopically-expressed heterologous IgaA could replace endogenous IgaA. To that purpose, we used P22 phage to transduce a null Δ*igaA2*::KXX allele.^3^ This allele was transduced to *S*. Typhimurium strains bearing the respective IgaA-expressing plasmids in the absence/presence of the inducer (0.2% arabinose) (Figure 2B). An additional positive control consisting of pBAD24 expressing the *S*. Typhimurium IgaA, was included.

Transductants bearing the Δ*igaA2*::KXX allele following induction with arabinose were obtained only in *S*. Typhimurium strains expressing IgaA of *S. flexneri* and *D. dadantii* (Figure 2C). This result showed that IgaA of *Y. enterocolitica*, *P. luminescens*, and *S. glossinidius*, which represent the *Yersiniaceae*, *Morganellaceae* and *Bruguierivoracaceae* families respectively, cannot repress the Rcs system when produced in *S*. Typhimurium, which belongs to the *Enterobacteriaceae* family. Interestingly, even in the presence of 2% glucose, some transductants bearing the Δ*igaA2*::KXX were obtained exclusively when producing ectopically IgaA of *S*. Typhimurium (see Figure 2C). Most of these transductants exhibited a mucoid phenotype, denoting overactivation of the Rcs system.^3^ These mucoid transductants lacking IgaA arise spontaneously in *S*. Typhimurium by compensatory mutations that decrease levels of RscC or RcsD.^5^ The reason for obtaining these mucoid clones in glucose-containing plates only in the strain bearing the plasmid with *igaA* from *S*. Typhimurium is at present unknown. Minute amounts of highly active endogenous IgaA produced from the plasmid even in the absence of inducer could be sufficient to allow the emergence of such compensatory mutations.

The data obtained following the expression in *S*. Typhimurium of heterologous IgaA, in which complementation was detected only in defined cases, supported variability in IgaA within the *Enterobacterales* order. We reasoned this variability could rely on structural differences that limit interactions with the endogenous Rcs proteins from *S*. Typhimurium and/or the recognition of yet unknown signalling molecules.

### Structural modelling of IgaA from *Enterobacterales* unravel conserved positions among families of the order and the presence of three small β-barrel (SBB) domains

Based on the phenotypic data obtained with the different IgaA expressed in *S*. Typhimurium (Figure 2), we searched for regions or residues that could vary in the non-complementing versus complementing IgaA variants by a sequence-based structural approach (Figure 3). To that purpose, we used the publicly available tool AlphaFold (https://alphafold.ebi.ac.uk/)^18,19^ to predict the structure of the different IgaA expressed in *S*. Typhimurium. As previously proposed,^4^ the overall structure predicted by AlphaFold comprises five transmembrane helices (TM), three defined cytoplasmic regions (cyt1, cyt2, cyt3) and a periplasmic region (Figure 4A-4B). A first unexpected finding resulting from modelling of IgaA was the presence of three β-rich architectures forming partially-closed small β-barrel (SBB) domains,^14^ which are related to OB (oligonucleotide/oligosaccharide binding motif) fold domains.^15^ The two SSB domains in cytoplasmic regions were termed SBB-1 and SBB-2 while the periplasmic domain was named SBB-3 (Figure 4A-4B). SBB-1, SBB-2 and SBB-3 share similarity to the fold present in a domain of the IgaA ortholog from *E. coli,* YrfF (residues 36-154, PDB: 4UZM),^20^ showing root-mean-square deviation (RMSD) values of 1.5 Å for 101 residues in SBB-1; 2.1 Å for 73 residues for SBB-2; and, 1.6 Å for 105 residues for SBB-3 (Figure S2). Each of the three SBB domains present in IgaA are formed by two β-sheets (β-sheet I and β-sheet II) where the first β-strand is connected to an α-helix and the second β-strand is curved being shared by both β-sheets (Figure 4C). In SSB-3, another β-strand (β19) is however curved and shared by the two β-sheets contributing to the β-barrel shape (Figure 4C). Remarkably, SSB-1 is located entirely in the first cytoplasmic region cyt1 whereas SSB-2 is "hybrid". Thus, SSB-2 is built by residues of region cyt1 that comprise the β7 strand (sequence 183-ELLNIRQ-189 in IgaA of *S*. Typhimurium) that is connected to α7 to form the "SBB-connector" as well as most of region cyt2 (Figure 4C).

**Figure 3.**
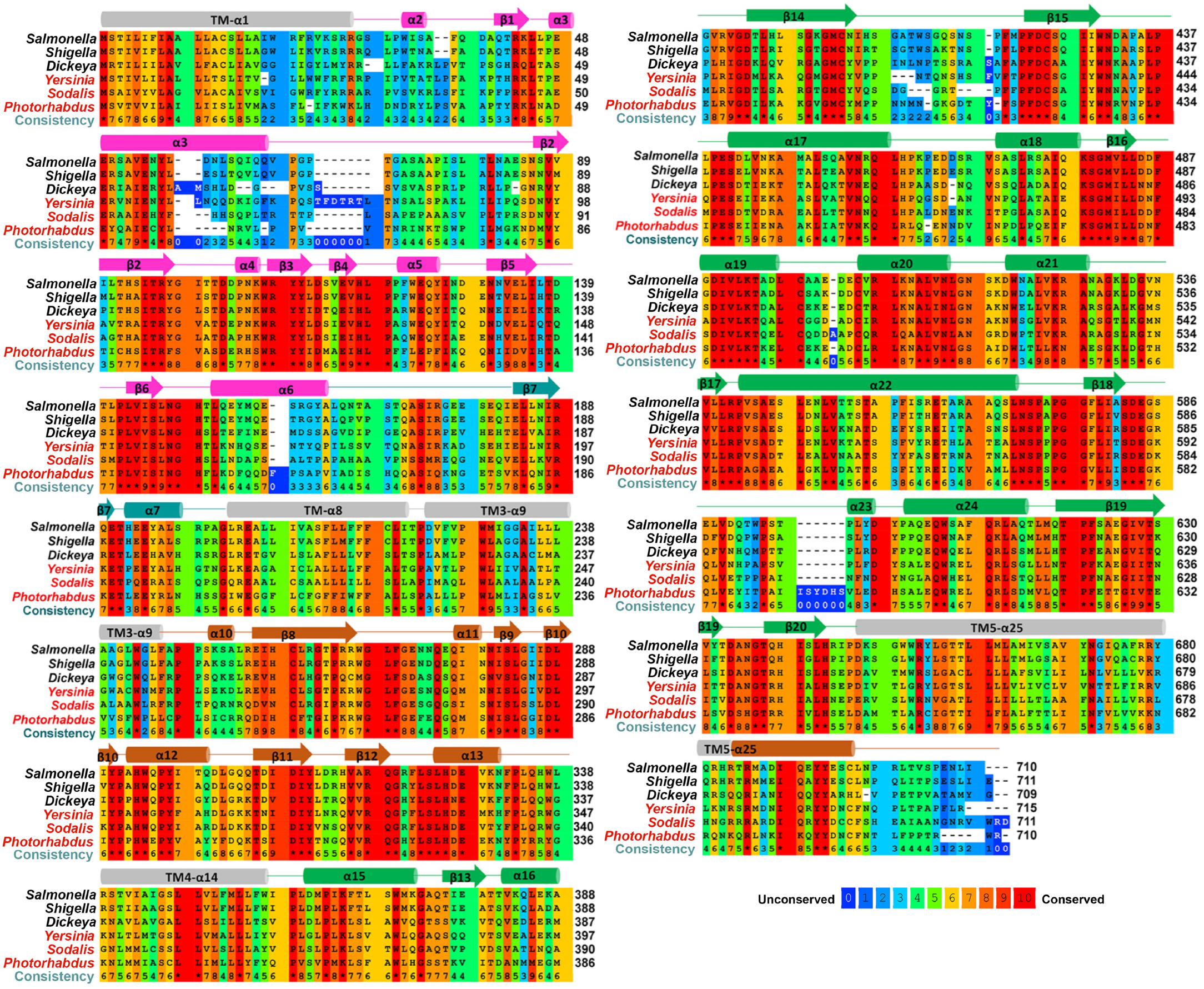
Alignment of IgaA sequences from the selected *Enterobacterales* genera that were produced in *S*. Typhimurium. IgaA sequences from the *Salmonella, Shigella, Dickeya, Yersinia*, *Photorhadbus* and *Sodalis* genera were aligned with the PRALINE server. Conservation of the residues is shown according to the colour key. The secondary structure is shown on the sequence where the structural elements are coloured according to Figure 4B (helix α in TMs in gray, SSB-1 domain in magenta, SBB connector in cyan, SBB-2 in orange and the periplasmic region having SBB-3 in green).

**Figure 4.**
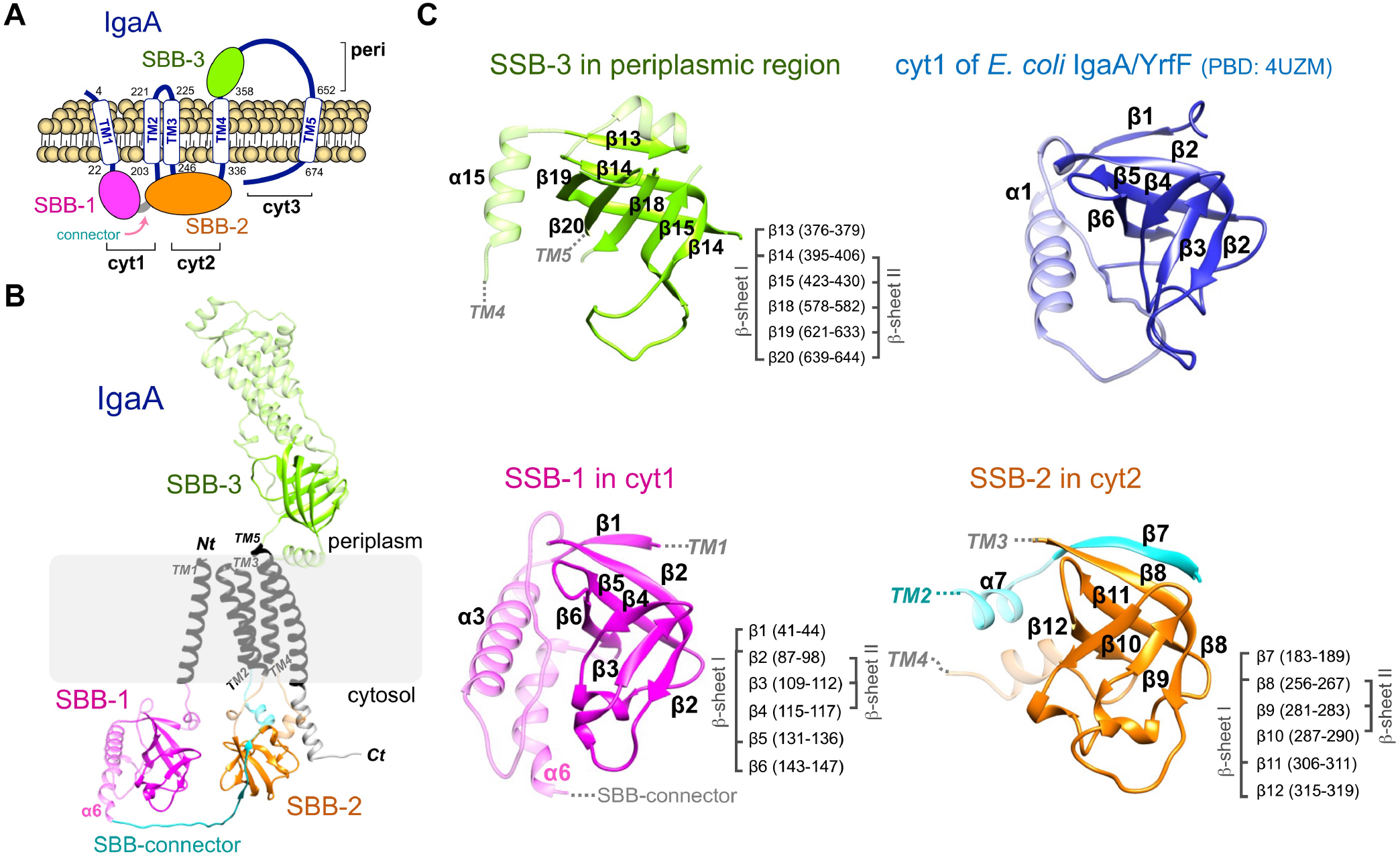
Modelling of IgaA structure reveals the presence of three SBB domains. (A) Cartoon showing the new domain configuration predicted for IgaA with two cytoplasmic SBB fold domains, with the named SBB-2 conformed by residues of regions cyt1 and cyt2. The C-terminus of region cyt3 is shown close to SBB-2 since at least residue R686 is predicted to interact with this domain (see Figure 6). The drawing also shows the presence of a connector between SBB-1 and SBB-2. Numbers indicate the positions of residues at the start and end of each of the five transmembrane domains.; (B) IgaA structure predicted by AlphaFold in which the five transmembrane regions are shown in dark grey, SSB-1 domain in magenta, SBB connector in cyan, SBB-2 in orange and the periplasmic SBB-3 in green; (C) The three SBB domains of *S*. Typhimurium IgaA predicted by AlphaFold and compared to the domain previously reported from a fragment of *E. coli* IgaA (YrfF).^20^ For each SBB domain the residues stretches involved in formation of β-sheet I and β-sheet II, are indicated.

We then modelled IgaA from the diverse *Enterobacterales* genera used in our functional study. Sequence identity with *Salmonella* is high for *Shigella* (84.1%) but moderate with the rest of the genera tested (49.3 % with *Dickeya*, 52.8% with *Yersinia*, 39.3% with *Photorhabdus*, and 47.7 % with *Sodalis*). All these IgaA variants conserve the three SBB domains (Figure S3). However, we observed sequence and length variability in several regions between the complementing and non-complementing IgaA variants (Figure 3). One was the loop between α3 and β2 in the SBB-1 domain, which is rather long in *Yersinia* and shorter in *Photorhabdus* (Figures 3, 5), existing also sequence divergence between the region spanning from the C-terminus of α3 and the N-terminus of the loop (Figure 3). Another variable region was the α6 helix in cyt1, which is almost absent in *Yersinia* and *Sodalis* whereas it is five residues shorter in *Photorhabdus*, showing sequence variability together with the N-terminal of the SBB connector (Figures 3, 5). Other regions showing sequence and length variability in the non-complementing IgaA variants include: i) the loop between β14-β15 in SBB-3; ii) the C-terminal end of TM-α1 connected to the N-terminal of β1 in SBB-1; and iii) the C-terminal end in cyt3 displaying a high degree of sequence variability (Figure 3).

**Figure 5.**
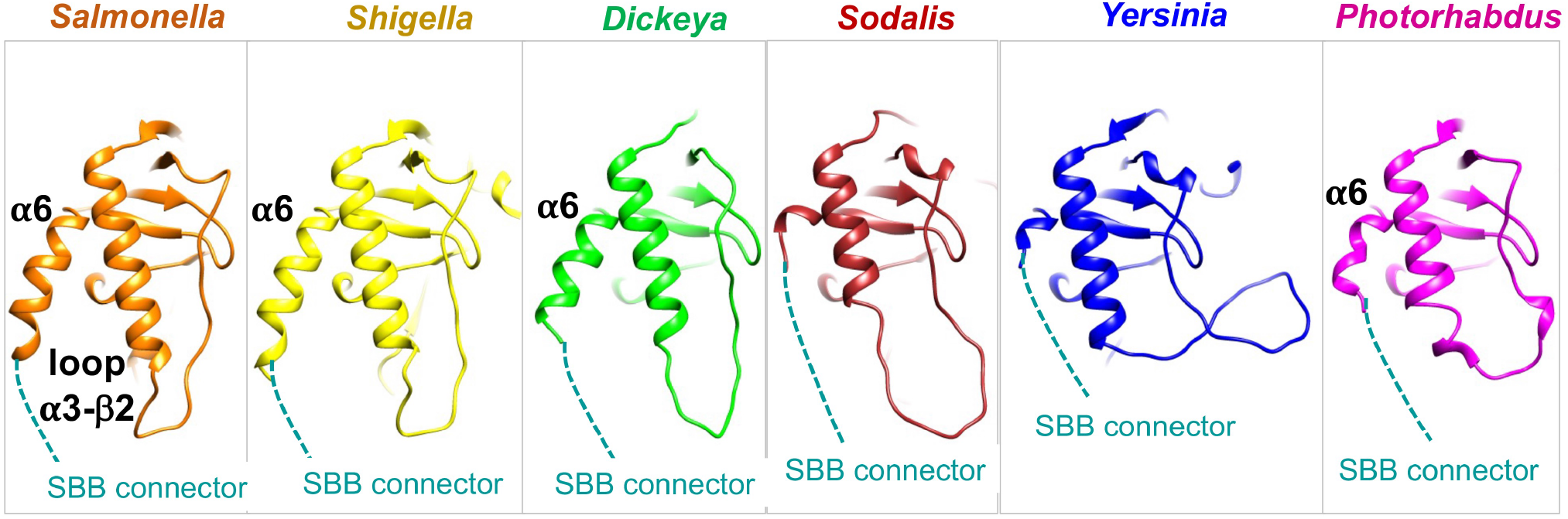
Structural features of the SBB-1 domain show divergence in the non-complementing IgaA variants. Modelled structure of the SBB-1 domain in cartoon representation showing differences in the length of loop α3-β2 and α6 in the *Enterobacterales* tested in this study: *Salmonella* (orange)*, Shigella* (yellow)*, Dickeya* (green)*, Sodalis* (maroon)*, Yersinia* (blue), *Photorhadbus* (magenta). The connection of α6 to the SBB connector is shown as a dashed line in cyan.

### The IgaA variants tested functionally in *S*. Typhimurium differ in some interacting sites involving the cyt1-cyt2 interface in the SBB-2 domain

We further exploited AlphaFold to search in IgaA for distances ≤ 4Å between residues, inferring in this manner interactions putatively relevant for domain stability. Among these interactions, we mapped E180-R265, which involves moderately conserved residues, one present in a stretch acting as SBB-1/SBB-2 connector (E180) to face another SBB-2 residue (R265) (Figure 6A). Additional interactions involving conserved residues seem to ensure folding of the SBB-2 domain. These include: i) the triad E194-R188-D309, which facilitates proximity between strand β7 of region cyt1 and SBB-2 residues that are part of the cytoplasmic region cyt2 (Figure 6B); and, ii) the triad H293-E328-R686, which implicates two residues of cytoplasmic region cyt2 forming part of SBB-2 and a residue of cytoplasmic region cyt3 (R686) (Figure 6B). Interestingly, an R188H mutation, probably disrupting the E194-R188-D309 interaction, was the first spontaneous mutation reported to affect IgaA function.^8^ The putative stabilizing role of R686 would implicate for the first time a functional role for the cyt3 region. Importantly, these predicted interactions sites are conserved in all IgaA that were expressed in *S*. Typhimurium (Figures 3, 6C). This evidence supports the idea of a "hybrid" SBB-2 domain conformed by residues of three different cytoplasmic regions of IgaA (cyt1, cyt2, cyt3) and absolutely required to repress the Rcs system.

**Figure 6.**
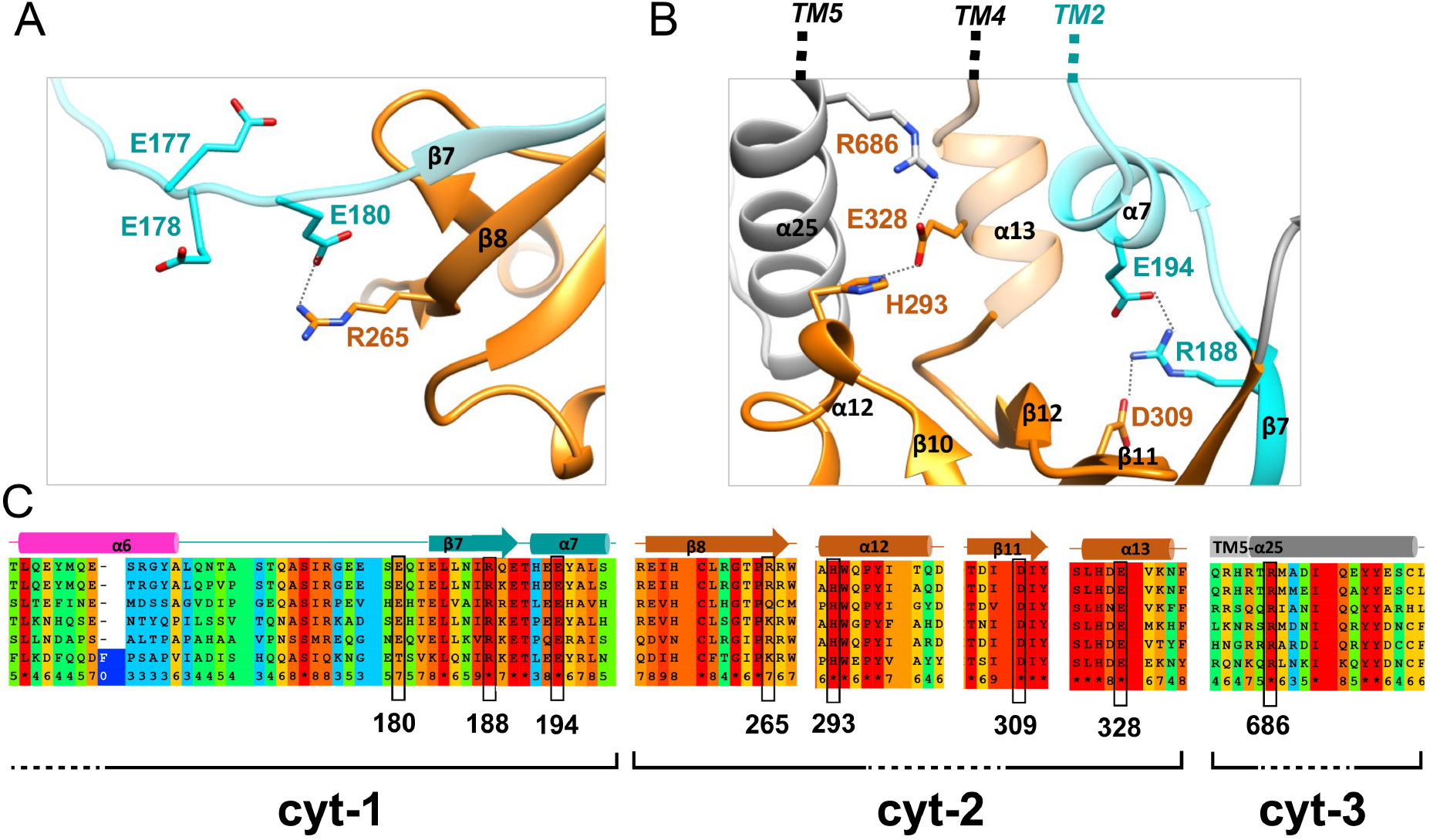
Conserved interactions in IgaA of diverse *Enterobacterales* genera connecting residues of the cyt1, cyt2 and cyt3 regions. Modelled structure in cartoon representation of *S*. Typhimurium IgaA showing salt bridge interactions conserved in the *Enterobacterales* tested. (A) Residue E180 in the SBB-connector (cyan) belonging to cyt1 interact with R265 located in the β8 of the SBB-2 domain in cyt2 (orange); (B) Residues in the β7 (R188) and α7 (E194) within the SBB-connector of cyt1 (cyan) interact with D309 of the SBB-2 (orange). Residues E328 and H293 in cyt2 (orange) interact with R686 in cyt3 (gray); (C) Sequence alignment, as shown in Figure 3, of the structural features shown in panels A and B. Dashed lines between residues indicate interacting distances in the modelled structures.

Next, we sought to get insights into the structural variations that could explain the lack of complementation exhibited by some of the IgaA expressed in *S*. Typhimurium (Figure 2). We searched in the AlphaFold structural predictions for residues that were not conserved when comparing those IgaA functional in *S*. Typhimurium (versions of *S. flexneri* and *D. dadantii*) versus those non-complementing (*Y. enterocolitica*, *P. luminescences* and *S. glossidinius*). We reasoned these divergent residues could be involved in interactions required for proper folding of IgaA in *S*. Typhimurium but probably not existing in IgaA of other *Enterobacterales* genera. H192-P249 was found as a putative interacting site diverging among the two groups of IgaA proteins, complementing versus non-complementing (Figure 7). Interestingly, H192 is located in α7, within the SBB-connector, contributing to the “hybrid” SBB-2 domain formed by residues of both cyt1 and cyt2 regions. H192 is also next to T191, a conserved residue previously described to be important for IgaA function,^4^ whose role can now be explained by its interaction with the conserved H326 (Figure 7). Thus, sequence diversity in the β7-α7 region within the SBB-connector contributing to the “hybrid” SBB2 domain seems to have an important impact in IgaA function not only providing anchoring to the SSB-2 domain but also a structural support. Other two sites differing between these two sets of IgaA proteins were R255-D313 and D287-R314, interaction sites that involve again the SBB-2 domain creating an electrostatic surface that varies in the non-complementing IgaA variants (Figure S4). Collectively, these findings reinforce the relevance that the "hybrid" SBB-2 domain has for IgaA function.

**Figure 7.**
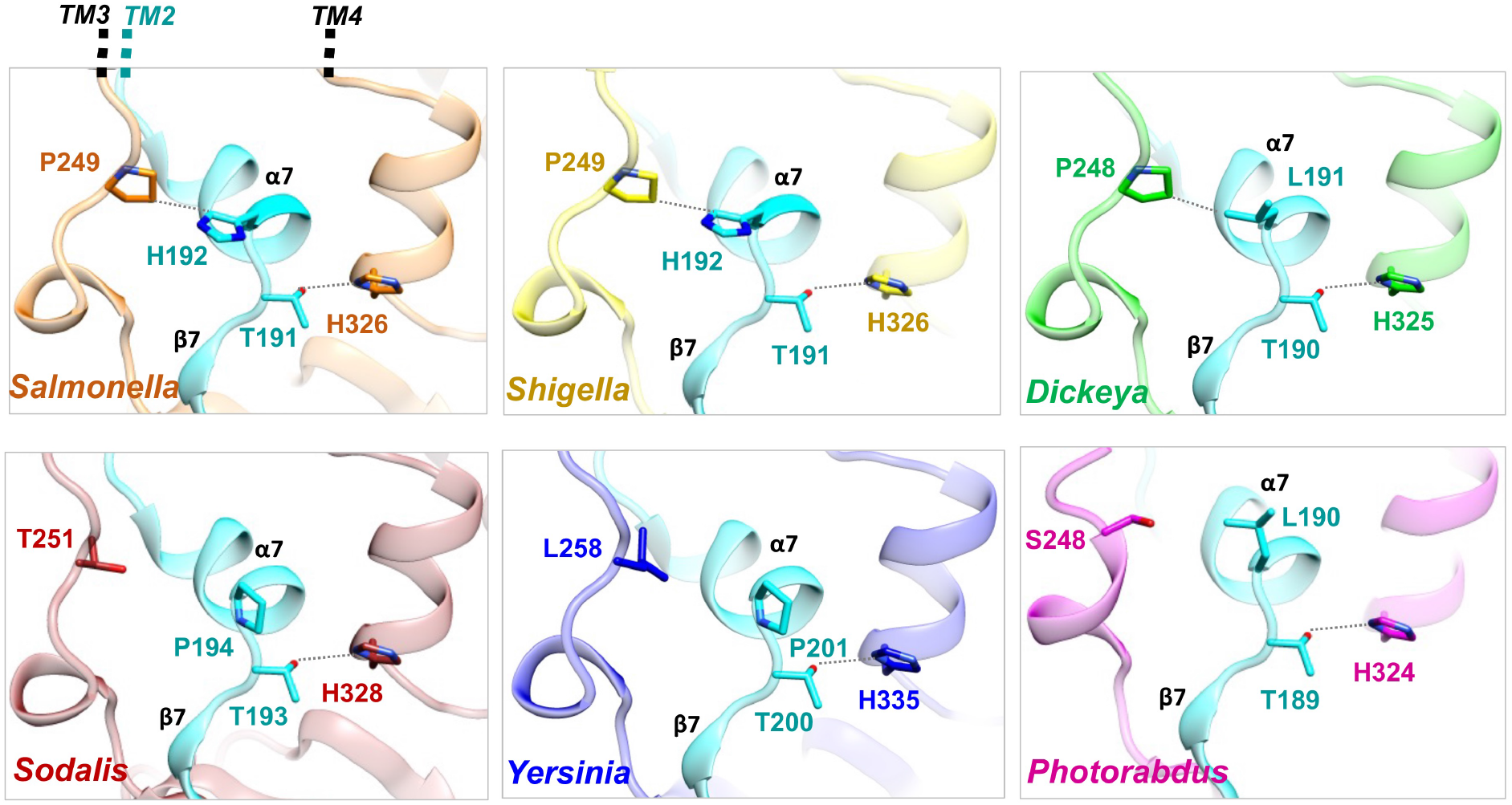
The SBB-connector provides specific anchoring to the SSB-2 domain as well as a structural support. Modelled structures by cartoon representation of the SBB-2 “hybrid” domain in the *Enterobacterales* tested in this study: *Salmonella* (orange), *Shigella* (yellow), *Dickeya* (green), *Sodalis* (maroon), *Yersinia* (blue), and *Photorhadbus* (magenta). Residues in the β7-α7 structural feature of the SBB-connector from cyt1 (in cyan) show divergence and conservation to contact the SBB-2. Dashed lines between residues indicate interacting distances in the modelled structures.

## DISCUSSION

In this study, we examined from an evolutionary perspective the repression that the inner membrane protein IgaA exerts over the Rcs system. Our strategy involving expression of different IgaA from distinct families of the *Enterobacterales* order was insightful to uncover regions critical for function and to establish the degree of conservation or divergence of this protein within this order. The functional data obtained together with the *in silico* analyses of the structures predicted by AlphaFold all point to a previously unnoticed presence of an SBB-2 domain, related to the OB fold domain.^15^ SBB-2 is formed by most of cytoplasmic region cyt2, a β-strand (β7) supplied by the C-terminal residues of cyt1 creating a clear cyt1-cyt2 interface and defined residues of the cyt3 region like R686. A previous study reported the presence of an OB fold domain in a fragment of *E. coli* IgaA (formerly YrfF) encompassing residues 37-154, all positioned in region cyt1,^20^ and which corresponds to what we termed SSB-1 in IgaA of *S*. Typhimurium. This consistency provides reliability to the AlphaFold prediction extended to full-length IgaA showing the presence of two additional SBB domains. SSB domains differ from β-barrels of integral membrane proteins in having much less β-sheets, in the order of five or six, and of shorter length.^14^ SSB domain are also of small size, generally encompassing ≤ 100 residues. Other striking features of the SSB domains are the orthogonal package of the β-sheets and the presence of short invariant α-helixes connecting such β-sheets.^14^ Remarkably, a distant OB fold domain was previously assigned to region cyt1 of *E. coli* IgaA^20^ whereas our structural analysis reveal that this domain should be more strictly considered as SSB (Figure S2). Other studies involving periplasmic proteins have predicted OB-fold domains, as it is the case of YdeI and YgiW of *S*. Typhimurium, two proteins that interact with outer membrane porins and play a role in antimicrobial resistance, stress and virulence.^21,22^ Intriguingly, YdeI is regulated positively by the Rcs system and important for *Salmonella* persistent infections.^23^ Considering our findings in IgaA, the presence of OB fold domains in these proteins should be revisited to dissect whether SSB domains, besides its wide presence in proteins related to DNA, RNA and protein metabolism, could also be common in proteins involved in stabilizing envelope architecture. To our knowledge, IgaA represents the first example of protein possessing SBB domains facing at both the outer and inner leaflets of the plasma membrane, opening the possibility of SBB domains being also involved in signal transduction.

In a recent study, Wall et al. provided compelling evidence in *E. coli* for IgaA interacting with RcsD at both their periplasmic and cytoplasmic regions.^7^ These authors generated constructs deficient in either cyt1 or cyt2 or the periplasmic region of IgaA, with all of them exhibiting loss-of-function. Of major interest, the cyt1 deletion encompassed residues 36-181,^7^ therefore making possible the presence an intact hybrid SBB-2 domain folded with the contribution of the β7 strand (see Figure 4). In the context of that study and our analyses, it can be hypothesized that the three SBB domains act in concert to transduce signal from the periplasm to the cytoplasm, being the basis of the mechanism by which IgaA could repress the Rcs system. The essential role played by the cytoplasmic regions of IgaA, as pointed by Wall et al.^7^ and further inferred from our study, also agrees with the data of Hussein et al.^6^ Despite restoring viability in *E. coli* transiently depleted of IgaA but expressing IgaA cyt1 and cyt2 fragments, these authors were unable to complement a Δ*igaA*::Km null allele with the individual fragments.^6^

The most accepted model of communication between IgaA and the Rcs system is a control at the periplasmic side exerted by an IgaA-RcsF interaction,^10^ which is relieved upon stress, an alteration that is supposed to be translated to cytoplasmic regions of IgaA. This model, however, contrasts at some extent with our functional studies with IgaA from different *Enterobacterales* families and the structural *in silico* analyses presented here. Thus, in the primary sequences of the IgaA tested functionally, and basically with the only exception of the region linking β14-β15 sheets, we did not find other motifs in the periplasmic region diverging between the complementing IgaA variants and those non-complementing of *Y. enterocolitica*, *S. glossidinius* and *P. luminescences* (Figure 3). By contrast, more variable sequences and structures were mapped in the cyt1, cyt2 and cyt3, regions, including the α3-β2 junction, α6 helix, P249, R255, D287, D313, R314, and the C-terminus (Figures 3, 5-7, and S4). Although we are uncertain for the reason of detecting more divergence in the cytoplasmic versus the periplasmic regions, the structural requirements in the periplasm might be less strict compared to cyt1 and cyt2. Supporting this idea is the capacity to repress the Rcs system shown by a recombinant IgaA in which its periplasmic domain was replaced for the equivalent region of MalF.^6^ Moreover, among the spontaneous mutations selected in *S*. Typhimurium for partial loss of function in IgaA, those mapping in cyt1 and cyt2 like R188H, T191P and G262R have a more profound phenotypic effect regarding the loss of repression over the Rcs system and attenuation of virulence.^4^ It is remarkable that R188H, the original mutation selected in *S*. Typhimurium for a phenotype of increased growth inside eukaryotic cells, which was used to coin this protein,^8^ involves a residue mapping in the β7 strand that links cyt1 to cyt2 to conform the hybrid SSB-2 domain (Figures 4,6).

Overall, our work represents a novel approach that has exploited evolutionary data to unveil the functional relevance of the cytoplasmic regions of IgaA and that is further sustained by *in silico* analyses. Moreover, our approach complements recent work by other groups ^6,7^ and provides a detail map of variable and conserved sites as well as structures directly related to function. These differences may have established structural constrains in the interaction of IgaA with RcsD specific for defined *Enterobacteriales* genera, which might not occur when the protein is expressed in *S*. Typhimurium. Additionally, it cannot be ruled out that IgaA from *Y. enterocolitica*, *P. luminescences* and *S. glossidinius* may not recognize in *S*. Typhimurium a hypothetical signalling molecule of different nature.

## MATERIALS AND METHODS

### Bacterial strains and growth conditions

All *S. enterica* serovar Typhimurium strains used in this study derive from SV5015 ^24^, a His^+^ prototroph of virulent strain SL1344 ^25^. *E. coli* strains used for cloning are derivate of strain DH10B ^26^. The bacterial strains and plasmids used in this study are listed in Table S2 in the Supporting Information. Bacteria were cultured in Luria-Bertani (LB) broth at 37ºC in shaking (150 rpm) conditions. To repress/induce expression of *igaA* gene cloned in plasmid, bacteria were grown in LB contaning 2% glucose or 0.2% arabinose, respectively, during 2 h. The sugars were added to the medium in mid exponential phase (OD600~0.5). When required, the medium was supplemented with ampicillin (50 µg/mL), or kanamycin (30 µg/mL).

### Cloning of *igaA* from different *Enterobacterales* genera

The *igaA* gene from selected *Enterobacterales* genera were cloned using genomic DNA from these sources: *D. dadantii* strain 3937 ^27^; *P. luminescens* strain TTO1 ^28^; *S. glossinidius* strain *morsitants* ^29^; *S. flexneri* strain M90T ^30^; *Y. enterocolitica* strain 8081 ^31^; and, *S*. Typhimurium strain SL1344 ^25^. *igaA* was PCR-amplified with the oligonucleotides listed in Table S3 of Supporting Information using Takara DNA polymerase. Oligonucleotides were designed with NheI/HindIII sites to allow cloning of PCR-products in pGEM-T cloning vector and the sequence of the Myc epitope (5’-CAG ATC CTC TTC TGA GAT GAG TTT TTG TTG −3’) in one of the primers (Table S3). From the pGEM-T plasmid, inserts were transferred to the expression vector pBAD24 for inducing expression with 0.2% arabinose. *igaA* from *D. dadantii*, *S*. Typhimurium, *Y. enterocolitica* and *S. glossinidius* were cloned in pBAD24 using NheI/HindIII enzymes. *igaA* of *P. luminescens* and *S. flexneri* were cloned in pBAD24 using XbaI/EcoRI sites to insert a SpeI/EcoRI product obtained from pGEM-T. In all cloning procedures, *E. coli* DH10B ^26^ was used a host. The series of pBAD24 plasmids with the different *igaA* variants were finally used to transform *S*. Typhimurium strain SL1344.

### Phage transduction and viability assays

IgaA is an essential protein in *S*. Typhimurium in a Rcs^+^ genetic background ^3^. To monitor the capacity of IgaA from other *Enterobacterales* genera to replace the endogenous protein, a P22 H105/1 *int201* transducing phage lysate obtained from *S*. Typhimurium strain SV4441 (*igaA2*::KXX *rcsC*::Mu*d*A), a gift from Prof. J. Casadesús, was used. This lysate was employed to transduce the null *igaA2*::KXX allele and select for transductants resistant to kanamycin, only obtained when the exogenous IgaA produced from plasmid complemented endogenous IgaA. Therefore, the appearance of transductants on plates containing 30 µg/mL kanamycin provided a direct proof for the activity of exogenous IgaA in *S*. Typhimurium.

### Immunoassays for detection of IgaA-Myc recombinant proteins

IgaA-Myc of different *Enterobacterales* genera produced in *S*. Typhimurium were detected with mouse monoclonal anti-Myc-tag antibody (clone #9B11, Cell Signaling Technology, Danvers, MA) at 1:1,000 dilution and secondary goat anti-mouse peroxidase at 1:2,000 dilution. The amount of sample loaded per well corresponded to ~5×10^7^ bacteria. Preparation of protein extracts, electrophoresis and immunoassay conditions were as described ^4^.

### Phylogenetic analysis of IgaA and Rcs proteins

A total of 114 genomes belonging to *Enterobacterales* order bacteria were retrieved from “The Bacterial and Viral Bioinformatics Resource Center (BV-BRC)” (https://www.bv-brc.org/) based on completeness, total contig number and the consideration of being “reference” or “representative” genomes. These genomes were interrogated using the BLASTP tool using as queries the following protein sequences from *S. enterica* serovar Typhimurium strain SL1344 (in brackets Uniprot accesion number): IgaA (E1WIS2), RscB (A0A0H3NNS0), RcsC (A0A0H3NF64), RscD (A0A0H3NDQ0), RcsF (A0A0H3N9F1) and DnaK (A0A0H3NCG3). The assignment for presence/absence considered BLASTP scores and the BV-BRC annotations. For those entries lacking functional annotation, a search of orthologs was performed using eggNOG 5.0 sequence search web service ^32^ in order to confirm that they belonged to the predicted orthologous group.

To compose the phylogenetic trees, two sets of amino acids sequences of IgaA, RcsB, RcsC, RcsD, RcsF and DnaK were built for each protein; one containing all the sequences predicted to be homologous found in the 114 proteomes and, the second with only with those belonging to some specific genera: *Salmonella, Shigella, Escherichia, Yersinia, Dickeya, Sodalis*, and *Photorhabdus*. Protein trees for each set were built using the BV-BRC GenTree tool with no end trimming or gappy sequences removal and using the “Randomized Axelerated Maximum Likelihood” (RAxML) algorithm with the LG general amino acid replacement matrix ^33^. Trees were visualized using iTOL version 6.5.7 ^34^.

### Structural analyses on models obtained by AlphaFold

The structure of *S*. Typhimurium IgaA was obtained from the AlphaFold database,^19^ code AF-E1WIS2-F1. The modelled structures of the diverse IgaA tested in this study, from *Shigella*, *Dickeya*, *Yersinia*, *Photorhadbus* and *Sodalis*, were obtained loading its sequence in the ColabFold,^35^ an open-source software to predict protein structure using AlphaFold2. Structural superposition was produced with Superpose and Gesamt, programs supported in the CCP4 Suite.^36^ Multiple sequence alignment was performed using PRALINE.^37^ The structure of the SBB domains found in modelled IgaA were compared with other proteins in the PDB using the DALI server.^38^ The cartoon representation of the structures was produced using UCSF Chimera.^39^

## SUPPLEMENTARY MATERIAL

**Figure S1. Phylogenetic tree of IgaA and Rcs proteins from 40 representative genera of the *Enterobacterales* order.** A total of 114 genomes deposited in the “The Bacterial and Viral Bioinformatics Resource Center” (BV-BRC) were analysed *in silico* for genes encoding IgaA and the indicated Rcs proteins. The DnaK chaperone was also analysed as unrelated protein. Sequences from the identified proteins were aligned to generate a Newick file. Protein trees were built using the BV-BRC Gen Tree tool with no end trimming or gappy sequences removal and using RAxML algorithm with LG evolutionary model. Trees were visualized using iTOL version 6.5.7. Bars indicate phylogenetic distance.

**Figure S2. Superposition of the SBB domains in the modelled structure of *Salmonella* IgaA with the NMR structure of the distant OB-fold related domain from *E. coli* IgaA (PDB: 4UZM).** Each of the SBB domains in cartoon representation (SBB-1 in magenta, SBB-2 in orange and SBB-3 in green) identified from the modelled IgaA structure are shown superposed to PDB: 4UZM (in blue) demonstrating structural similarities. The connectivity of the SBB domains to the TM regions are also indicated.

**Figure S3. Modelled structures of heterologous IgaA variants from distinct *Enterobacterales* genera that were expressed in *S*. Typhimurium show the same three SBB domains.** Cartoon representation of the cytoplasmic and periplasmic regions of the modelled structures of IgaA in *Shigella* (yellow), *Dickeya* (green), *Sodalis* (maroon), *Yersinia* (blue), *Photorhadbus* (magenta) showing the superposition of the SBB domains identified in IgaA from *Salmonella* (SBB-1 in magenta, SBB-2 in orange and SBB-3 in green).

**Figure S4. Residues in the SBB-2 domain contribute as an electrostatic surface that varies along the *Enterobacterales* genera studied**. Cartoon representation of a surface within the SBB-2 domain in the modelled structures of IgaA from *Salmonella* (orange), *Shigella* (yellow), *Dickeya* (green), *Sodalis* (maroon), *Yersinia* (blue) and *Photorhadbus* (magenta) where residues involved in salt-bridges in IgaA from *Salmonella* show divergence. Dashed lines between residues indicate interacting distances in the modelled structure of IgaA.

**Table S1.** Presence/absence of IgaA/RcsB/RcsC/RcsD/RcsF orthologs in bacteria of the order *Enterobacterales*.

**Table S2.** Metadata obtained from BV-BRC database for IgaA/RcsB/RcsC/RcsD/RcsF orthologs in bacteria of the order *Enterobacterales*.

**Table S3.** Genera and species of the order *Enterobacterales* selected for the functional analysis of IgaA orthologs.

**Table S4.** *S*. Typhimurium/*E. coli* strains and plasmids used in this study.

**Table S5.** Primer oligonucleotides used in this study.

## ACKNOWLEDGEMENTS

This work was supported by grant PID2020-112971GB-I00/10.13039/501100011033 (to F.G.dP.) and PID2019-110630GB-I00 (to P.C.) from the Spanish Ministry of Science and Innovation. We thank L.A. Fernández (Centro Nacional de Biotecnología-CSIC, Madrid, Spain) for *E. coli* strain DH10B; H. Niki (National Institute of Genetics, Shizuoka, Japan) for pBAD24 plasmid; J. Casadesús (University of Seville, Spain) for *S*. Typhimurium strain SV4441; D. Clarke (University College Cork, Ireland) for *P. luminescens* strain TTO1; and Henar González (CNB-CSIC) for technical support.

