## Supplementary material for "Evolutionary analysis in *Enterobacterales* of the Rcs-repressor protein IgaA unveils two cytoplasmic small β-barrel domains central for function": Figures-S1-to-S4-Tables-S1-to-S5

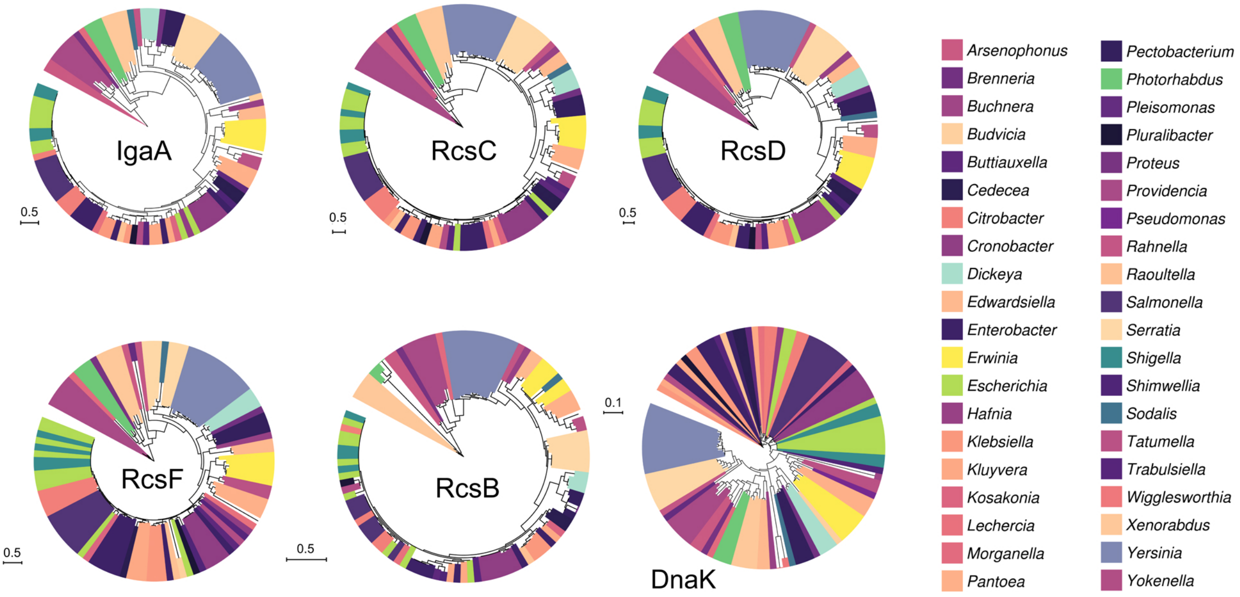
IgaA
RcsC
RcsD
RcsF
RcsB
DnaK
0.5
0.5
0.5
0.5
0.5
0.1
Arsenophonus
Brenneria
Buchnera
Budvicia
Buttiauxella
Cedecea
Citrobacter
Cronobacter
Dickeya
Edwardsiella
Enterobacter
Erwinia
Escherichia
Hafnia
Klebsiella
Kluyvera
Kosakonia
Lechercia
Morganella
Pantoea
Pectobacterium
Photorhabdus
Pleisomonas
Pluralibacter
Proteus
Providencia
Pseudomonas
Rahnella
Raoultella
Salmonella
Serratia
Shigella
Shimwellia
Sodalis
Tatumella
Trabulsiella
Wigglesworthia
Xenorabdus
Yersinia
Yokenella

**Figure S1**

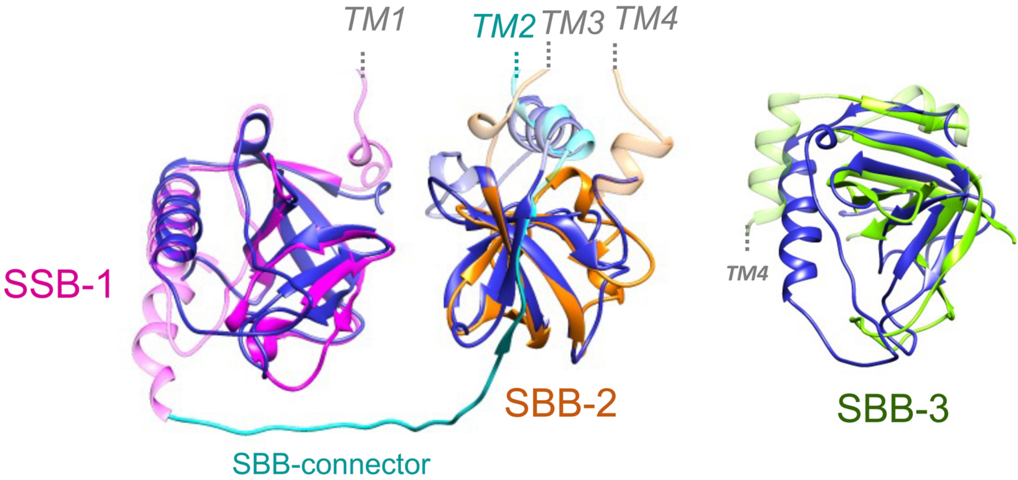


**Figure S2**

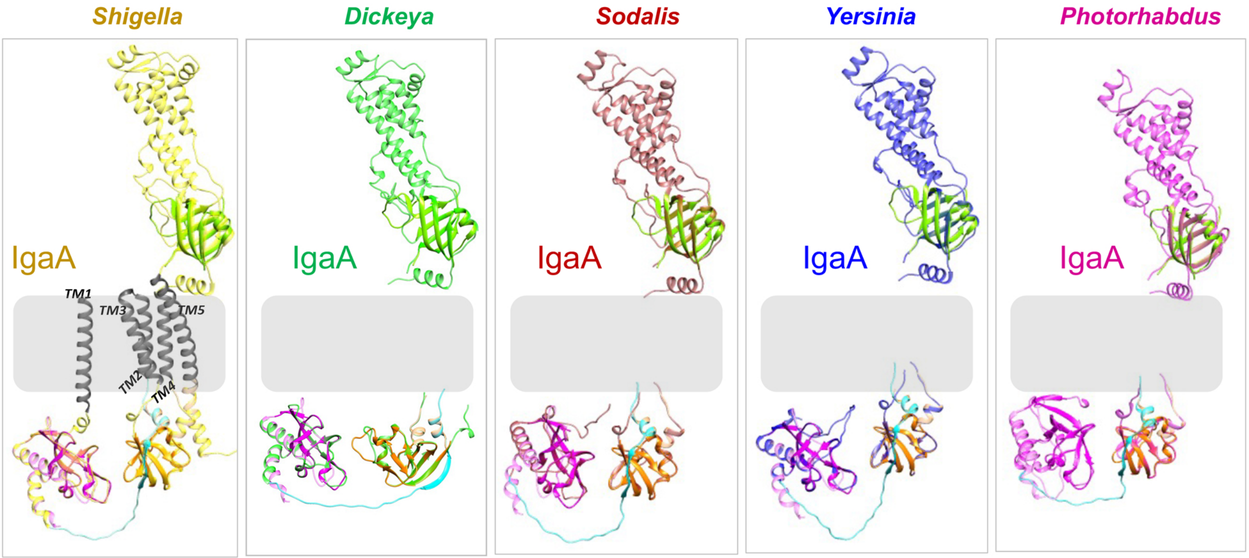

Shigella
Dickeya
Sodalis
Yersinia
Photorhabdus
IgaA
TM1
TM3
TM5
TM2
TM4
IgaA
IgaA
IgaA
IgaA

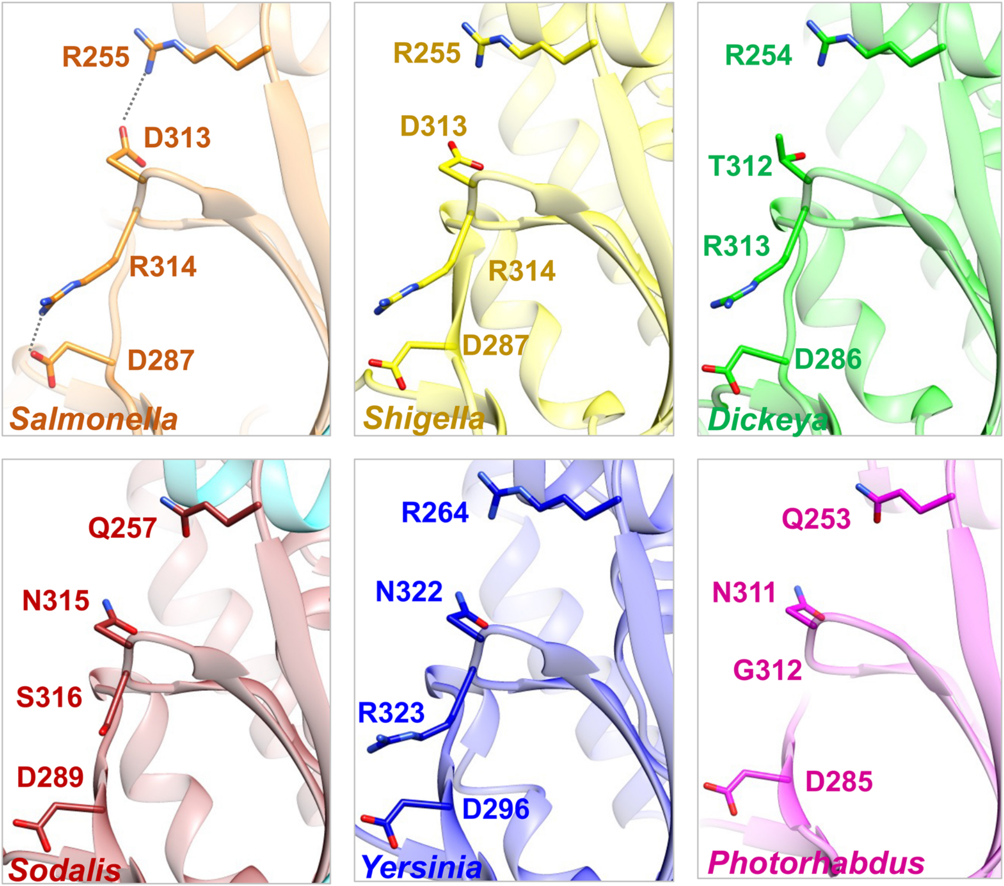
R255
D313
R314
D287
Salmonella
R255
D313
R314
D287
Shigella
R254
T312
R313
D286
Dickeya
Q257
N315
S316
D289
Sodalis
R264
N322
R323
D296
Yersinia
Q253
N311
G312
D285
Photorhabdus

**Figure S4**

**Table S1.** Presence/absence of IgaA/RcsB/RcsC/RcsD/RcsF orthologs in bacteria of the order *Enterobacterales*

Absence
Presence of truncated protein
Presence in only some strains of the species

| | FROM PATRIC ANNOTATION | | | | |
|---|---|---|---|---|---|
| | **IgaA** | **RcsB** | **RcsC** | **RcsD** | **RcsF** |
| *Yokenella regensburgei ATCC 43003* | IgaA | RcsB | RcsC | RcsD | RcsF |
| *Tatumella ptyseos ATCC 33301* | IgaA | RcsB | RcsC | RcsD | RcsF |
| *Kluyvera ascorbata ATCC 33433* | IgaA | RcsB | RcsC | RcsD | RcsF |
| *Klebsiella oxytoca KCTC 1686* | IgaA | RcsB | RcsC | RcsD | RcsF |
| *Enterobacter aerogenes KCTC 2190* | IgaA | RcsB | RcsC | RcsD | RcsF |
| *Cronobacter condimenti 1330* | IgaA | RcsB | RcsC | RcsD | RcsF |
| *Cronobacter universalis NCTC 9529* | IgaA | RcsB | RcsC | RcsD | RcsF |
| *Pantoea rwandensis ND04* | IgaA | RcsB | RcsC | RcsD | RcsF |
| *Budvicia aquatica DSM 5075* | IgaA | RcsB | RcsC | RcsD | RcsF |
| *Escherichia hermannii NBRC 105704* | IgaA | RcsB | RcsC | RcsD | RcsF |
| *Escherichia vulneris NBRC 102420* | IgaA | RcsB | RcsC | RcsD | RcsF |
| *Arsenophonus nasoniae DSM 15247* | IgaA | RcsB | RcsC | RcsD | RcsF |
| *Enterobacter pulveris DSM 19144* | IgaA | RcsB | RcsC | RcsD | RcsF |
| *Morganella morganii subsp. morganii KT* | IgaA | RcsB | RcsC | RcsD | RcsF |
| *Escherichia coli O104:H4 str. 2011C-3493* | IgaA | RcsB | RcsC | RcsD | RcsF |
| *Providencia sneebia DSM 19967* | IgaA | RcsB | RcsC | RcsD | RcsF |
| *Providencia burhodogranariea DSM 19968* | IgaA | RcsB | RcsC | RcsD | RcsF |
| *Providencia rettgeri Dmel1* | IgaA | RcsB | RcsC | RcsD | RcsF |
| *Providencia stuartii MRSN 2154* | IgaA | RcsB | RcsC | RcsD | RcsF |
| *Cronobacter malonaticus 681* | IgaA | RcsB | RcsC | RcsD | RcsF |
| *Kluyvera cryocrescens NBRC 102467* | IgaA | RcsB | RcsC | RcsD | RcsF |
| *Dickeya paradisiaca NCPPB 2511* | IgaA | RcsB | RcsC | RcsD | RcsF |
| *Kosakonia sacchari SP1* | IgaA | RcsB | RcsC | RcsD | RcsF |
| *Arsenophonus endosymbiont str. Hangzhou of Nilaparvata lugens* | IgaA | RcsB | RcsC | RcsD | RcsF |
| *Raoultella ornithinolytica B6* | IgaA | RcsB | RcsC | RcsD | RcsF |
| *Citrobacter freundii CFNIH1* | IgaA | RcsB | RcsC | RcsD | RcsF |
| *Pseudomonas flectens ATCC 12775* | IgaA | RcsB | RcsC | RcsD | RcsF |
| *Serratia liquefaciens ATCC 27592* | IgaA | RcsB | RcsC | RcsD | RcsF |
| *Cronobacter zurichensis LMG 23730* | IgaA | RcsB | RcsC | RcsD | RcsF |
| *Xenorhabdus szentirmaii DSM 16338* | IgaA | RcsB | RcsC | RcsD | RcsF |
| *Serratia fonticola RB-25* | IgaA | RcsB | RcsC | RcsD | RcsF |
| *Hafnia alvei FB1* | IgaA | RcsB | RcsC | RcsD | RcsF |
| *Cedecea neteri M006* | IgaA | RcsB | RcsC | RcsD | RcsF |
| *Shigella flexneri 5a str. M90T* | IgaA | RcsB | RcsC | RcsD | RcsF |
| *Dickeya dadantii 3937* | IgaA | RcsB | RcsC | RcsD | RcsF |
| *Salmonella enterica subsp. enterica serovar Typhi str. Ty2* | IgaA | RcsB | RcsC | RcsD | RcsF |
| *Yersinia pestis CO92* | IgaA | RcsB | RcsC | RcsD | RcsF |
| *Salmonella enterica subsp. enterica serovar Typhimurium str. SL1344* | IgaA | RcsB | RcsC | RcsD | RcsF |
| *Pectobacterium atrosepticum SCRI1043* | IgaA | RcsB | RcsC | RcsD | RcsF |
| *Salmonella bongori NCTC 12419* | IgaA | RcsB | RcsC | RcsD | RcsF |
| *Salmonella enterica subsp. enterica serovar Typhi str. CT18* | IgaA | RcsB | RcsC | RcsD | RcsF |
| *Buchnera aphidicola str. Bp (Baizongia pistaciae)* | IgaA | RcsB | RcsC | RcsD | RcsF |
| *Photorhabdus temperata subsp. thracensis strain DSM 15199* | IgaA | RcsB | RcsC | RcsD | RcsF |
| *Photorhabdus luminescens subsp. laumondii TTO1* | IgaA | RcsB | RcsC | RcsD | RcsF |
| *Klebsiella pneumoniae subsp. pneumoniae MGH 78578* | IgaA | RcsB | RcsC | RcsD | RcsF |
| *Yersinia pseudotuberculosis IP 32953* | IgaA | RcsB | RcsC | RcsD | RcsF |
| *Serratia marcescens subsp. marcescens Db11* | IgaA | RcsB | RcsC | RcsD | RcsF |
| *Citrobacter koseri ATCC BAA-895* | IgaA | RcsB | RcsC | RcsD | RcsF |
| *Cronobacter sakazakii ATCC BAA-894* | IgaA | RcsB | RcsC | RcsD | RcsF |
| *Photorhabdus asymbiotica strain ATCC 43949* | IgaA | RcsB | RcsC | RcsD | RcsF |
| *Yersinia ruckeri strain Big Creek 74* | IgaA | RcsB | RcsC | RcsD | RcsF |
| *Salmonella enterica subsp. enterica serovar Paratyphi A str. ATCC 9150* | IgaA | RcsB | RcsC | RcsD | RcsF |
| *Shigella dysenteriae Sd197* | IgaA | RcsB | RcsC | RcsD | RcsF |
| *Shigella boydii Sb227* | IgaA | RcsB | RcsC | RcsD | RcsF |
| *Shigella sonnei Ss046* | IgaA | RcsB | RcsC | RcsD | RcsF |
| *Sodalis glossinidius str. 'morsitans'* | IgaA | RcsB | RcsC | RcsD | RcsF |
| *Yersinia intermedia ATCC 29909* | IgaA | RcsB | RcsC | RcsD | RcsF |
| *Yersinia frederiksenii ATCC 33641* | IgaA | RcsB | RcsC | RcsD | RcsF |
| *Yersinia mollaretii ATCC 43969* | IgaA | RcsB | RcsC | RcsD | RcsF |
| *Yersinia bercovieri ATCC 43970* | IgaA | RcsB | RcsC | RcsD | RcsF |
| *Xenorhabdus doucetiae strain FRM16 = DSM 17909* | IgaA | RcsB | RcsC | RcsD | RcsF |
| *Wigglesworthia glossinidia endosymbiont of Glossina morsitans* | IgaA | RcsB | RcsC | RcsD | RcsF |
| *Wigglesworthia glossinidia endosymbiont of Glossina brevipalpis* | IgaA | RcsB | RcsC | RcsD | RcsF |
| *Trabulsiella odontotermitis strain TbO2.3* | IgaA | RcsB | RcsC | RcsD | RcsF |
| *Escherichia coli O157:H7 str. Sakai* | IgaA | RcsB | RcsC | RcsD | RcsF |
| *Yersinia enterocolitica subsp. enterocolitica 8081* | IgaA | RcsB | RcsC | RcsD | RcsF |
| *Serratia proteamaculans 568* | IgaA | RcsB | RcsC | RcsD | RcsF |
| *Xenorhabdus nematophila ATCC 19061* | IgaA | RcsB | RcsC | RcsD | RcsF |
| *Xenorhabdus bovienii SS-2004* | IgaA | RcsB | RcsC | RcsD | RcsF |
| *Salmonella enterica subsp. arizonae serovar 62:z4,z23:- strain RSK2980* | IgaA | RcsB | RcsC | RcsD | RcsF |
| *Erwinia tasmaniensis Et1/99* | IgaA | RcsB | RcsC | RcsD | RcsF |
| *Edwardsiella tarda EIB202* | IgaA | RcsB | RcsC | RcsD | RcsF |
| *Enterobacter cancerogenus ATCC 35316* | IgaA | RcsB | RcsC | RcsD | RcsF |
| *Citrobacter youngae ATCC 29220* | IgaA | RcsB | RcsC | RcsD | RcsF |
| *Escherichia albertii TW07627* | IgaA | RcsB | RcsC | RcsD | RcsF |
| *Escherichia coli str. K-12 substr. MG1655* | IgaA | RcsB | RcsC | RcsD | RcsF |
| *Providencia alcalifaciens DSM 30120* | IgaA | RcsB | RcsC | RcsD | RcsF |
| *Yersinia aldovae ATCC 35236* | IgaA | RcsB | RcsC | RcsD | RcsF |
| *Yersinia rohdei ATCC 43380* | IgaA | RcsB | RcsC | RcsD | RcsF |
| *Yersinia kristensenii ATCC 33638* | IgaA | RcsB | RcsC | RcsD | RcsF |

| | | | | | |
|---|---|---|---|---|---|
| *Proteus mirabilis HI4320* | IgaA | RcsB | RcsC | RcsD | RcsF |
| *Pantoea agglomerans strain FDAARGOS_160* | IgaA | RcsB | RcsC | RcsD | RcsF |
| *Dickeya zeae Ech1591* | IgaA | RcsB | RcsC | RcsD | RcsF |
| *Pectobacterium carotovorum subsp. carotovorum PC1* | IgaA | RcsB | RcsC | RcsD | RcsF |
| *Pectobacterium wasabiae WPP163* | IgaA | RcsB | RcsC | RcsD | RcsF |
| *Cedecea davisae DSM 4568* | IgaA | RcsB | RcsC | RcsD | RcsF |
| *Escherichia fergusonii ATCC 35469* | IgaA | RcsB | RcsC | RcsD | RcsF |
| *Escherichia coli UMN026* | IgaA | RcsB | RcsC | RcsD | RcsF |
| *Salmonella enterica subsp. enterica serovar Typhimurium str. 14028S* | IgaA | RcsB | RcsC | RcsD | RcsF |
| *Brenneria sp. EniD312* | IgaA | RcsB | RcsC | RcsD | RcsF |
| *Pluralibacter gergoviae FB2* | IgaA | RcsB | RcsC | RcsD | RcsF |
| *Kluyvera intermedia strain CAV1151* | IgaA | RcsB | RcsC | RcsD | RcsF |
| *Shimwellia blattae DSM 4481 = NBRC 105725* | IgaA | RcsB | RcsC | RcsD | RcsF |
| *Erwinia pyrifoliae Ep1/96* | IgaA | RcsB | RcsC | RcsD | RcsF |
| *Erwinia billingiae Eb661* | IgaA | RcsB | RcsC | RcsD | RcsF |
| *Edwardsiella ictaluri 93-146* | IgaA | RcsB | RcsC | RcsD | RcsF |
| *Citrobacter rodentium ICC168* | IgaA | RcsB | RcsC | RcsD | RcsF |
| *Klebsiella variicola At-22* | IgaA | RcsB | RcsC | RcsD | RcsF |
| *Enterobacter asburiae LF7a* | IgaA | RcsB | RcsC | RcsD | RcsF |
| *Tatumella morbirosei LMG 23360* | IgaA | RcsB | RcsC | RcsD | RcsF |
| *Erwinia amylovora CFBP1430* | IgaA | RcsB | RcsC | RcsD | RcsF |
| *Serratia odorifera DSM 4582* | IgaA | RcsB | RcsC | RcsD | RcsF |
| *Escherichia coli O83:H1 str. NRG 857C* | IgaA | RcsB | RcsC | RcsD | RcsF |
| *Cronobacter turicensis z3032* | IgaA | RcsB | RcsC | RcsD | RcsF |
| *Enterobacter lignolyticus SCF1* | IgaA | RcsB | RcsC | RcsD | RcsF |
| *Plesiomonas shigelloides strain NCTC10360* | IgaA | RcsB | RcsC | RcsD | RcsF |
| *Pantoea ananatis LMG 20103* | IgaA | RcsB | RcsC | RcsD | RcsF |
| *Erwinia amylovora ATCC 49946* | IgaA | RcsB | RcsC | RcsD | RcsF |
| *Enterobacter cloacae subsp. cloacae ATCC 13047* | IgaA | RcsB | RcsC | RcsD | RcsF |
| *Rahnella aquatilis CIP 78.65 = ATCC 33071* | IgaA | RcsB | RcsC | RcsD | RcsF |
| *Serratia plymuthica AS9* | IgaA | RcsB | RcsC | RcsD | RcsF |
| *Buttiauxella agrestis strain MCE* | IgaA | RcsB | RcsC | RcsD | RcsF |
| *Leclercia adecarboxylata strain USDA-ARS-USMARC-60222* | IgaA | RcsB | RcsC | RcsD | RcsF |
| *Enterobacter hormaechei ATCC 49162* | IgaA | RcsB | RcsC | RcsD | RcsF |

**Table S2.** Metadata obtained from BV-BRC database for IgaA/RcsB/RcsC/RcsD/RcsF orthologs in bacteria of the order *Enterobacterales*

| BRC.ID | homolog | Genome | Genome.ID | Accession | RefSeq.L | Alt.Locus | Feature.I | Annotation | Feature.T | Start | End | Length | Strand | FIGfam.II | PATRIC.g | PATRIC.( | Protein.II | AA.Leng | Gene.Sy | Product | GO |
|---|---|---|---|---|---|---|---|---|---|---|---|---|---|---|---|---|---|---|---|---|---|
| fig\|124702 | DnaK | Arsenophonus endosymbiont str. Hangzhou of Nilaparva | 1247024.4 | JRLH01000004 | | | PATRIC.12 | PATRIC | CDS | 39340 | 41265 | 1926 | + | FIG000233 | PLF_637_ | PGF_10357457 | | 641 | | Chaperone protein DnaK | |
| fig\|124702 | RcsF | Arsenophonus endosymbiont str. Hangzhou of Nilaparva | 1247024,4 | JRLH01000008 | | | PATRIC.12 | PATRIC | CDS | 30517 | 30915 | 399 | - | FIG013040 | PLF_637_ | PGF_04751602 | | 132 | | hypothetical protein | |
| fig\|124702 | IgaA | Arsenophonus endosymbiont str. Hangzhou of Nilaparva | 1247024,4 | JRLH01000008 | | | PATRIC.12 | PATRIC | CDS | 55806 | 57950 | 2145 | + | FIG000040 | PLF_637_ | PGF_00013790 | | 714 | | IgaA: a membrane protein that prevents overactivation of the Rcs regulatory system | |
| fig\|124702 | RcsD | Arsenophonus endosymbiont str. Hangzhou of Nilaparva | 1247024,4 | JRLH01000002 | | | PATRIC.12 | PATRIC | CDS | 57081 | 59786 | 2706 | + | FIG000045 | PLF_637_ | PGF_00034034 | | 901 | | Phosphotransferase RcsD | |
| fig\|124702 | RcsB | Arsenophonus endosymbiont str. Hangzhou of Nilaparva | 1247024,4 | JRLH01000002 | | | PATRIC.12 | PATRIC | CDS | 59788 | 60435 | 648 | + | FIG000041 | PLF_637_ | PGF_00421954 | | 215 | | DNA-binding capsular synthesis response regulator RcsB | |
| fig\|124702 | RcsC | Arsenophonus endosymbiont str. Hangzhou of Nilaparva | 1247024,4 | JRLH01000002 | | | PATRIC.12 | PATRIC | CDS | 60552 | 63341 | 2790 | - | FIG000042 | PLF_637_ | PGF_00050577 | | 929 | | Sensor hist | GO:0004673\|protein histidine kinase activity |
| fig\|112101 | DnaK | Arsenophonus nasoniae DSM 15247 | 1121018.3 | AUCC01000109 | | VBIArsNas | PATRIC.11 | PATRIC | CDS | 3598 | 5523 | 1926 | + | FIG000233 | PLF_637_ | PGF_10357457 | | 641 | | Chaperone protein DnaK | |
| fig\|112101 | IgaA | Arsenophonus nasoniae DSM 15247 | 1121018,3 | AUCC01000051 | | VBIArsNas | PATRIC.11 | PATRIC | CDS | 16688 | 18832 | 2145 | + | FIG000040 | PLF_637_ | PGF_03915705 | | 714 | | hypothetical protein | |
| fig\|112101 | RcsD | Arsenophonus nasoniae DSM 15247 | 1121018,3 | AUCC01000010 | | VBIArsNas | PATRIC.11 | PATRIC | CDS | 7240 | 9945 | 2706 | + | FIG000045 | PLF_637_ | PGF_00034034 | | 901 | | Phosphotransferase RcsD | |
| fig\|112101 | RcsB | Arsenophonus nasoniae DSM 15247 | 1121018,3 | AUCC01000010 | | VBIArsNas | PATRIC.11 | PATRIC | CDS | 9947 | 10594 | 648 | + | FIG000041 | PLF_637_ | PGF_00421954 | | 215 | | DNA-binding capsular synthesis response regulator RcsB | |
| fig\|112101 | RcsC | Arsenophonus nasoniae DSM 15247 | 1121018,3 | AUCC01000010 | | VBIArsNas | PATRIC.11 | PATRIC | CDS | 10712 | 13345 | 2634 | - | FIG000042 | PLF_637_ | PGF_00050577 | | 877 | | Sensor hist | GO:0004673\|protein histidine kinase activity |
| fig\|598467 | DnaK | Brenneria sp. EniD312 | 598467.3 | AFWW01000003 | | VBIBreSp1 | PATRIC.59 | PATRIC | CDS | 123662 | 125578 | 1917 | + | FIG000233 | PLF_7165 | PGF_10357457 | | 638 | | Chaperone protein DnaK | |
| fig\|598467 | RcsD | Brenneria sp. EniD312 | 598467,3 | AFWW01000010 | | VBIBreSp1 | PATRIC.59 | PATRIC | CDS | 2258335 | 2261001 | 2667 | + | FIG000045 | PLF_7165 | PGF_00034034 | | 888 | | Phosphotransferase RcsD | |
| fig\|598467 | RcsB | Brenneria sp. EniD312 | 598467,3 | AFWW01000010 | | VBIBreSp1 | PATRIC.59 | PATRIC | CDS | 2260994 | 2261650 | 657 | + | FIG000041 | PLF_7165 | PGF_00421954 | | 218 | | DNA-binding capsular synthesis response regulator RcsB | |
| fig\|598467 | RcsC | Brenneria sp. EniD312 | 598467,3 | AFWW01000010 | | VBIBreSp1 | PATRIC.59 | PATRIC | CDS | 2261706 | 2264570 | 2865 | - | FIG000042 | PLF_7165 | PGF_00050577 | | 954 | | Sensor hist | GO:0004673\|protein histidine kinase activity |
| fig\|598467 | IgaA | Brenneria sp. EniD312 | 598467,3 | AFWW01000001 | | VBIBreSp1 | PATRIC.59 | PATRIC | CDS | 435056 | 437197 | 2142 | - | FIG000040 | PLF_7165 | PGF_00013790 | | 713 | | IgaA: a membrane protein that prevents overactivation of the Rcs regulatory system | |
| fig\|598467 | RcsF | Brenneria sp. EniD312 | 598467,3 | AFWW01000003 | | VBIBreSp1 | PATRIC.59 | PATRIC | CDS | 407250 | 407483 | 234 | - | FIG013040 | PLF_7165 | PGF_00037945 | | 77 | | Protein RcsF | |
| fig\|224915 | DnaK | Buchnera aphidicola str. Bp (Baizongia pistaciae) | 224915.9 | NC_004545 | bbp142 | VBIBucAp | PATRIC.22 | PATRIC | CDS | 158590 | 160506 | 1917 | - | FIG000233 | PLF_3219 | PGF_1035 | NP_77777 | 638 | dnaK | Chaperone protein DnaK | |
| fig\|111172 | DnaK | Budvicia aquatica DSM 5075 | 1111728.3 | ATYS01000002 | | VBIBudAq | PATRIC.11 | PATRIC | CDS | 108566 | 110476 | 1911 | + | FIG00023369 | | PGF_10357457 | | 636 | | Chaperone protein DnaK | |
| fig\|111172 | IgaA | Budvicia aquatica DSM 5075 | 1111728,3 | ATYS01000011 | | VBIBudAq | PATRIC.11 | PATRIC | CDS | 49538 | 51721 | 2184 | + | FIG00004077 | | PGF_00013790 | | 727 | | IgaA: a membrane protein that prevents overactivation of the Rcs regulatory system | |
| fig\|111172 | RcsF | Budvicia aquatica DSM 5075 | 1111728,3 | ATYS01000001 | | VBIBudAq | PATRIC.11 | PATRIC | CDS | 235955 | 236230 | 276 | - | | | PGF_00037945 | | 91 | | Protein RcsF | |
| fig\|82977.3 | DnaK | Buttiauxella agrestis strain MCE | 82977.3 | JPRU01000002 | | | PATRIC.82 | PATRIC | CDS | 1361656 | 1363575 | 1920 | - | FIG000233 | PLF_8297 | PGF_10357457 | | 639 | | Chaperone protein DnaK | |
| fig\|82977.3 | RcsD | Buttiauxella agrestis strain MCE | 82977,3 | JPRU01000001 | | | PATRIC.82 | PATRIC | CDS | 1518919 | 1521576 | 2658 | + | FIG000045 | PLF_8297 | PGF_00034034 | | 885 | | Phosphotransferase RcsD | |
| fig\|82977.3 | RcsB | Buttiauxella agrestis strain MCE | 82977,3 | JPRU01000001 | | | PATRIC.82 | PATRIC | CDS | 1521593 | 1522243 | 651 | + | FIG000041 | PLF_8297 | PGF_00421954 | | 216 | | DNA-binding capsular synthesis response regulator RcsB | |
| fig\|82977.3 | RcsC | Buttiauxella agrestis strain MCE | 82977,3 | JPRU01000001 | | | PATRIC.82 | PATRIC | CDS | 1522383 | 1525235 | 2853 | - | FIG000042 | PLF_8297 | PGF_00050577 | | 950 | | Sensor hist | GO:0004673\|protein histidine kinase activity |
| fig\|82977.3 | IgaA | Buttiauxella agrestis strain MCE | 82977,3 | JPRU01000003 | | | PATRIC.82 | PATRIC | CDS | 45936 | 48077 | 2142 | + | FIG000040 | PLF_8297 | PGF_00013790 | | 713 | | IgaA: a membrane protein that prevents overactivation of the Rcs regulatory system | |
| fig\|82977.3 | RcsF | Buttiauxella agrestis strain MCE | 82977,3 | JPRU01000004 | | | PATRIC.82 | PATRIC | CDS | 247673 | 248077 | 405 | - | FIG013040 | PLF_8297 | PGF_00037945 | | 134 | | Protein RcsF | |
| fig\|566551 | DnaK | Cedecea davisae DSM 4568 | 566551.4 | ATDT01000003 | HMPREF0 | VBICedDa | PATRIC.56 | PATRIC | CDS | 420539 | 422461 | 1923 | + | FIG000233 | PLF_1584 | PGF_1035 | EPF20740 | 640 | | Chaperone protein DnaK | |
| fig\|566551 | RcsD | Cedecea davisae DSM 4568 | 566551,4 | ATDT01000026 | HMPREF0 | VBICedDa | PATRIC.56 | PATRIC | CDS | 190229 | 192889 | 2661 | + | FIG000045 | PLF_1584 | PGF_0003 | EPF15942 | 886 | | Phosphotransferase RcsD | |
| fig\|566551 | RcsB | Cedecea davisae DSM 4568 | 566551,4 | ATDT01000026 | HMPREF0 | VBICedDa | PATRIC.56 | PATRIC | CDS | 192906 | 193556 | 651 | + | FIG000041 | PLF_1584 | PGF_0042 | EPF15943 | 216 | | DNA-binding capsular synthesis response regulator RcsB | |
| fig\|566551 | RcsC | Cedecea davisae DSM 4568 | 566551,4 | ATDT01000026 | HMPREF0 | VBICedDa | PATRIC.56 | PATRIC | CDS | 193631 | 196483 | 2853 | - | FIG000042 | PLF_1584 | PGF_0005 | EPF15944 | 950 | | Sensor hist | GO:0004673\|protein histidine kinase activity |
| fig\|566551 | IgaA | Cedecea davisae DSM 4568 | 566551,4 | ATDT01000036 | HMPREF0 | VBICedDa | PATRIC.56 | PATRIC | CDS | 50910 | 53045 | 2136 | + | FIG000040 | PLF_1584 | PGF_0001 | EPF13158 | 711 | | IgaA: a membrane protein that prevents overactivation of the Rcs regulatory system | |
| fig\|566551 | RcsF | Cedecea davisae DSM 4568 | 566551,4 | ATDT01000003 | HMPREF0 | VBICedDa | PATRIC.56 | PATRIC | CDS | 603044 | 603175 | 132 | - | FIG013040 | PLF_1584 | PGF_0003 | EPF20912 | 43 | | Protein RcsF | |
| fig\|158822 | DnaK | Cedecea neteri M006 | 158822.7 | CP009458 | | | PATRIC.15 | PATRIC | CDS | 1817991 | 1819913 | 1923 | + | FIG000233 | PLF_1584 | PGF_10357457 | | 640 | | Chaperone protein DnaK | |
| fig\|158822 | RcsD | Cedecea neteri M006 | 158822,7 | CP009458 | | | PATRIC.15 | PATRIC | CDS | 4526795 | 4529458 | 2664 | + | FIG000045 | PLF_1584 | PGF_00034034 | | 887 | | Phosphotransferase RcsD | |
| fig\|158822 | RcsB | Cedecea neteri M006 | 158822,7 | CP009458 | | | PATRIC.15 | PATRIC | CDS | 4529475 | 4530125 | 651 | + | FIG000041 | PLF_1584 | PGF_00421954 | | 216 | | DNA-binding capsular synthesis response regulator RcsB | |
| fig\|158822 | RcsC | Cedecea neteri M006 | 158822,7 | CP009458 | | | PATRIC.15 | PATRIC | CDS | 4530188 | 4533043 | 2856 | - | FIG000042 | PLF_1584 | PGF_00050577 | | 951 | | Sensor hist | GO:0004673\|protein histidine kinase activity |
| fig\|158822 | IgaA | Cedecea neteri M006 | 158822,7 | CP009458 | | | PATRIC.15 | PATRIC | CDS | 778591 | 780726 | 2136 | + | FIG000040 | PLF_1584 | PGF_00013790 | | 711 | | IgaA: a membrane protein that prevents overactivation of the Rcs regulatory system | |
| fig\|133384 | DnaK | Citrobacter freundii CFNIH1 | 1333848.3 | CP007557 | CFNIH1_09795 | | PATRIC.13 | PATRIC | CDS | 2025203 | 2027125 | 1923 | + | FIG000233 | PLF_544_ | PGF_1035 | AHY11785 | 640 | dnaK | Chaperone protein DnaK | |
| fig\|133384 | RcsF | Citrobacter freundii CFNIH1 | 1333848,3 | CP007557 | CFNIH1_10660 | | PATRIC.13 | PATRIC | CDS | 2227459 | 2227812 | 354 | - | FIG013040 | PLF_544_ | PGF_0003 | AHY11954 | 117 | rcsF | Protein RcsF | |
| fig\|133384 | RcsD | Citrobacter freundii CFNIH1 | 1333848,3 | CP007557 | CFNIH1_22705 | | PATRIC.13 | PATRIC | CDS | 4643914 | 4646583 | 2670 | + | FIG000045 | PLF_544_ | PGF_0003 | AHY14237 | 889 | | Phosphotransferase RcsD | |
| fig\|133384 | RcsB | Citrobacter freundii CFNIH1 | 1333848,3 | CP007557 | CFNIH1_22710 | | PATRIC.13 | PATRIC | CDS | 4646600 | 4647250 | 651 | + | FIG000041 | PLF_544_ | PGF_0042 | AHY14238 | 216 | | DNA-binding capsular synthesis response regulator RcsB | |
| fig\|133384 | RcsC | Citrobacter freundii CFNIH1 | 1333848,3 | CP007557 | CFNIH1_22715 | | PATRIC.13 | PATRIC | CDS | 4647338 | 4650184 | 2847 | - | FIG000042 | PLF_544_ | PGF_0005 | AHY14239 | 948 | | Sensor hist | GO:0004673\|protein histidine kinase activity |
| fig\|133384 | IgaA | Citrobacter freundii CFNIH1 | 1333848,3 | CP007557 | CFNIH1_04655 | | PATRIC.13 | PATRIC | CDS | 937621 | 939756 | 2136 | + | FIG000040 | PLF_544_ | PGF_0001 | AHY10830 | 711 | | IgaA: a membrane protein that prevents overactivation of the Rcs regulatory system | |
| fig\|290338 | DnaK | Citrobacter koseri ATCC BAA-895 | 290338.8 | NC_009792 | CKO_0337 | VBICitKos | PATRIC.29 | PATRIC | CDS | 3143094 | 3145010 | 1917 | - | FIG000233 | PLF_544_ | PGF_1035 | YP_00145 | 638 | dnaK | Chaperone protein DnaK | |
| fig\|290338 | RcsF | Citrobacter koseri ATCC BAA-895 | 290338,8 | NC_009792 | CKO_0316 | VBICitKos | PATRIC.29 | PATRIC | CDS | 2935339 | 2935692 | 354 | + | FIG013040 | PLF_544_ | PGF_0003 | YP_00145 | 117 | rcsF | Protein RcsF | |
| fig\|290338 | IgaA | Citrobacter koseri ATCC BAA-895 | 290338,8 | NC_009792 | CKO_0482 | VBICitKos | PATRIC.29 | PATRIC | CDS | 4419050 | 4421182 | 2133 | + | FIG000040 | PLF_544_ | PGF_0001 | YP_00145 | 710 | | IgaA: a membrane protein that prevents overactivation of the Rcs regulatory system | |
| fig\|290338 | RcsC | Citrobacter koseri ATCC BAA-895 | 290338,8 | NC_009792 | CKO_0055 | VBICitKos | PATRIC.29 | PATRIC | CDS | 535953 | 538799 | 2847 | + | FIG000042 | PLF_544_ | PGF_0005 | YP_00145 | 948 | | Sensor hist | GO:0004673\|protein histidine kinase activity |
| fig\|290338 | RcsB | Citrobacter koseri ATCC BAA-895 | 290338,8 | NC_009792 | CKO_0055 | VBICitKos | PATRIC.29 | PATRIC | CDS | 538893 | 539543 | 651 | - | FIG000041 | PLF_544_ | PGF_0042 | YP_00145 | 216 | | DNA-binding capsular synthesis response regulator RcsB | |
| fig\|290338 | RcsD | Citrobacter koseri ATCC BAA-895 | 290338,8 | NC_009792 | CKO_0055 | VBICitKos | PATRIC.29 | PATRIC | CDS | 539560 | 542229 | 2670 | - | FIG000045 | PLF_544_ | PGF_0003 | YP_00145 | 889 | | Phosphotransferase RcsD | |
| fig\|637910 | DnaK | Citrobacter rodentium ICC168 | 637910.3 | NC_013716 | ROD_0010 | VBICitRod | PATRIC.63 | PATRIC | CDS | 11731 | 13647 | 1917 | + | FIG000233 | PLF_544_ | PGF_1035 | YP_00336 | 638 | dnaK | Chaperone protein DnaK | |
| fig\|637910 | RcsF | Citrobacter rodentium ICC168 | 637910,3 | NC_013716 | ROD_0206 | VBICitRod | PATRIC.63 | PATRIC | CDS | 243731 | 244027 | 297 | - | FIG013040 | PLF_544_ | PGF_0003 | YP_00336 | 98 | rcsF | Protein RcsF | |
| fig\|637910 | RcsD | Citrobacter rodentium ICC168 | 637910,3 | NC_013716 | ROD_2349 | VBICitRod | PATRIC.63 | PATRIC | CDS | 2466916 | 2469513 | 2598 | + | FIG000045 | PLF_544_ | PGF_0003 | YP_00336 | 865 | rcsD | Phosphotransferase RcsD | |
| fig\|637910 | RcsB | Citrobacter rodentium ICC168 | 637910,3 | NC_013716 | ROD_2350 | VBICitRod | PATRIC.63 | PATRIC | CDS | 2469530 | 2470180 | 651 | + | FIG000041 | PLF_544_ | PGF_0042 | YP_00336 | 216 | rcsB | DNA-binding capsular synthesis response regulator RcsB | |
| fig\|637910 | RcsC | Citrobacter rodentium ICC168 | 637910,3 | NC_013716 | ROD_2352 | VBICitRod | PATRIC.63 | PATRIC | CDS | 2470529 | 2473375 | 2847 | - | FIG000042 | PLF_544_ | PGF_0005 | YP_00336 | 948 | rcsC | Sensor hist | GO:0004673\|protein histidine kinase activity |
| fig\|637910 | IgaA | Citrobacter rodentium ICC168 | 637910,3 | NC_013716 | ROD_4425 | VBICitRod | PATRIC.63 | PATRIC | CDS | 4678571 | 4680691 | 2121 | - | FIG000040 | PLF_544_ | PGF_0001 | YP_00336 | 706 | | IgaA: a membrane protein that prevents overactivation of the Rcs regulatory system | |
| fig\|500640 | DnaK | Citrobacter youngae ATCC 29220 | 500640.5 | NZ_DS995323 | CIT292_00 | VBICitYou | PATRIC.50 | PATRIC | CDS | 373620 | 375539 | 1920 | + | FIG000233 | PLF_544_ | PGF_1035 | ZP_03835 | 639 | | Chaperone protein DnaK | |
| fig\|500640 | RcsD | Citrobacter youngae ATCC 29220 | 500640,5 | NZ_DS995324 | CIT292_03 | VBICitYou | PATRIC.50 | PATRIC | CDS | 2368837 | 2371506 | 2670 | + | FIG000045 | PLF_544_ | PGF_0003 | ZP_03837 | 889 | | Phosphotransferase RcsD | |
| fig\|500640 | RcsB | Citrobacter youngae ATCC 29220 | 500640,5 | NZ_DS995324 | CIT292_03 | VBICitYou | PATRIC.50 | PATRIC | CDS | 2371523 | 2372173 | 651 | + | FIG000041 | PLF_544_ | PGF_0042 | ZP_03837 | 216 | | DNA-binding capsular synthesis response regulator RcsB | |
| fig\|500640 | RcsC | Citrobacter youngae ATCC 29220 | 500640,5 | NZ_DS995324 | CIT292_03 | VBICitYou | PATRIC.50 | PATRIC | CDS | 2372275 | 2375121 | 2847 | - | FIG000042 | PLF_544_ | PGF_0005 | ZP_03837 | 948 | | Sensor hist | GO:0004673\|protein histidine kinase activity |
| fig\|500640 | IgaA | Citrobacter youngae ATCC 29220 | 500640,5 | NZ_DS995322 | CIT292_00 | VBICitYou | PATRIC.50 | PATRIC | CDS | 416804 | 418939 | 2136 | - | FIG000040 | PLF_544_ | PGF_0001 | ZP_03834 | 711 | | IgaA: a membrane protein that prevents overactivation of the Rcs regulatory system | |
| fig\|500640 | RcsF | Citrobacter youngae ATCC 29220 | 500640,5 | NZ_DS995323 | | VBICitYou | PATRIC.50 | PATRIC | CDS | 584339 | 584638 | 300 | - | FIG013040 | PLF_544_ | PGF_00037945 | | 99 | | Protein RcsF | |
| fig\|107399 | DnaK | Cronobacter condimenti 1330 | 1073999.4 | CAKW01000072 | BN137_20 | VBICroCon | PATRIC.10 | PATRIC | CDS | 18161 | 18316 | 156 | - | FIG000233 | PLF_4134 | PGF_1035 | CCJ72662. | 51 | | Chaperone protein DnaK | |
| fig\|107399 | IgaA | Cronobacter condimenti 1330 | 1073999,4 | CAKW01000068 | BN137_19 | VBICroCon | PATRIC.10 | PATRIC | CDS | 69355 | 71493 | 2139 | - | FIG000040 | PLF_4134 | PGF_0001 | CCJ72549. | 712 | | IgaA: a membrane protein that prevents overactivation of the Rcs regulatory system | |
| fig\|107399 | RcsD | Cronobacter condimenti 1330 | 1073999,4 | CAKW01000106 | | VBICroCon | PATRIC.10 | PATRIC | CDS | 13239 | 15770 | 2532 | + | FIG000045 | PLF_4134 | PGF_00034034 | | 843 | | Phosphotransferase RcsD | |
| fig\|107399 | RcsD | Cronobacter condimenti 1330 | 1073999,4 | CAKW01000106 | BN137_29 | VBICroCon | PATRIC.10 | PATRIC | CDS | 15785 | 15910 | 126 | + | FIG000045 | PLF_4134 | PGF_0003 | CCJ73597. | 41 | | Phosphotransferase RcsD | |
| fig\|107399 | RcsB | Cronobacter condimenti 1330 | 1073999,4 | CAKW01000106 | BN137_29 | VBICroCon | PATRIC.10 | PATRIC | CDS | 15927 | 16577 | 651 | + | FIG000041 | PLF_4134 | PGF_0042 | CCJ73598. | 216 | | DNA-binding capsular synthesis response regulator RcsB | |
| fig\|107399 | RcsC | Cronobacter condimenti 1330 | 1073999,4 | CAKW01000106 | BN137_29 | VBICroCon | PATRIC.10 | PATRIC | CDS | 16743 | 19592 | 2850 | - | FIG000042 | PLF_4134 | PGF_0005 | CCJ73599. | 949 | | Sensor hist | GO:0004673\|protein histidine kinase activity |
| fig\|120859 | DnaK | Cronobacter malonaticus 681 | 1208591.3 | CALC01000087 | BN131_34 | VBICroMal | PATRIC.12 | PATRIC | CDS | 86977 | 88893 | 1917 | + | FIG000233 | PLF_4134 | PGF_1035 | CCJ95778. | 638 | | Chaperone protein DnaK | |
| fig\|120859 | RcsD | Cronobacter malonaticus 681 | 1208591,3 | CALC01000087 | BN131_10 | VBICroMal | PATRIC.12 | PATRIC | CDS | 2448419 | 2450275 | 1857 | + | FIG000045 | PLF_4134 | PGF_0003 | CCJ93410. | 618 | | Phosphotransferase RcsD | |
| fig\|120859 | RcsD | Cronobacter malonaticus 681 | 1208591,3 | CALC01000087 | BN131_10 | VBICroMal | PATRIC.12 | PATRIC | CDS | 2450257 | 2451222 | 966 | + | FIG000045 | PLF_4134 | PGF_0003 | CCJ93411. | 321 | | Phosphotransferase RcsD | |
| fig\|120859 | RcsB | Cronobacter malonaticus 681 | 1208591,3 | CALC01000087 | BN131_10 | VBICroMal | PATRIC.12 | PATRIC | CDS | 2451239 | 2451889 | 651 | + | FIG000041 | PLF_4134 | PGF_0042 | CCJ93412. | 216 | | DNA-binding capsular synthesis response regulator RcsB | |
| fig\|120859 | RcsC | Cronobacter malonaticus 681 | 1208591,3 | CALC01000087 | BN131_10 | VBICroMal | PATRIC.12 | PATRIC | CDS | 2452176 | 2455016 | 2841 | - | FIG000042 | PLF_4134 | PGF_0005 | CCJ93416. | 946 | | Sensor hist | GO:0004673\|protein histidine kinase activity |
| fig\|120859 | RcsF | Cronobacter malonaticus 681 | 1208591,3 | CALC01000087 | BN131_12 | VBICroMal | PATRIC.12 | PATRIC | CDS | 310663 | 311067 | 405 | - | FIG013040 | PLF_4134 | PGF_0003 | CCJ93573. | 134 | | Protein RcsF | |
| fig\|120859 | IgaA | Cronobacter malonaticus 681 | 1208591,3 | CALC01000087 | BN131_21 | VBICroMal | PATRIC.12 | PATRIC | CDS | 3432365 | 3434503 | 2139 | + | FIG000040 | PLF_4134 | PGF_0001 | CCJ94439. | 712 | | IgaA: a membrane protein that prevents overactivation of the Rcs regulatory system | |
| fig\|290339 | DnaK | Cronobacter sakazakii ATCC BAA-894 | 290339.8 | NC_009778 | ESA_0332 | VBICroSak | PATRIC.29 | PATRIC | CDS | 3259878 | 3261794 | 1917 | - | FIG000233 | PLF_4134 | PGF_1035 | YP_00143 | 638 | dnaK | Chaperone protein DnaK | |
| fig\|290339 | RcsF | Cronobacter sakazakii ATCC BAA-894 | 290339,8 | NC_009778 | ESA_0314 | VBICroSak | PATRIC.29 | PATRIC | CDS | 3057860 | 3058264 | 405 | + | FIG013040 | PLF_4134 | PGF_0003 | YP_00143 | 134 | rcsF | Protein RcsF | |
| fig\|290339 | IgaA | Cronobacter sakazakii ATCC BAA-894 | 290339,8 | NC_009778 | ESA_0434 | VBICroSak | PATRIC.29 | PATRIC | CDS | 4311090 | 4313228 | 2139 | - | FIG000040 | PLF_4134 | PGF_0001 | YP_00144 | 712 | | IgaA: a membrane protein that prevents overactivation of the Rcs regulatory system | |
| fig\|290339 | RcsC | Cronobacter sakazakii ATCC BAA-894 | 290339,8 | NC_009778 | ESA_0097 | VBICroSak | PATRIC.29 | PATRIC | CDS | 941473 | 944322 | 2850 | + | FIG000042 | PLF_4134 | PGF_0005 | YP_00143 | 949 | | Sensor hist | GO:0004673\|protein histidine kinase activity |
| fig\|290339 | RcsB | Cronobacter sakazakii ATCC BAA-894 | 290339,8 | NC_009778 | ESA_0097 | VBICroSak | PATRIC.29 | PATRIC | CDS | 944494 | 945144 | 651 | - | FIG000041 | PLF_4134 | PGF_0042 | YP_00143 | 216 | | DNA-binding capsular synthesis response regulator RcsB | |
| fig\|290339 | RcsD | Cronobacter sakazakii ATCC BAA-894 | 290339,8 | NC_009778 | ESA_0097 | VBICroSak | PATRIC.29 | PATRIC | CDS | 945161 | 947791 | 2631 | - | FIG000045 | PLF_4134 | PGF_0003 | YP_00143 | 876 | | Phosphotransferase RcsD | |
| fig\|693216 | DnaK | Cronobacter turicensis z3032 | 693216.3 | FN543093 | Ctu_06390 | VBICroTur | PATRIC.69 | PATRIC | CDS | 690310 | 692226 | 1917 | + | FIG000233 | PLF_4134 | PGF_1035 | CBA27860 | 638 | dnaK | Chaperone protein DnaK | |
| fig\|693216 | RcsD | Cronobacter turicensis z3032 | 693216,3 | FN543093 | Ctu_28730 | VBICroTur | PATRIC.69 | PATRIC | CDS | 2999033 | 3001687 | 2655 | + | FIG000045 | PLF_4134 | PGF_0003 | CBA32353 | 884 | yojN | Phosphotransferase RcsD | |

| | | | | | | | | | | | | | | | | | | | | |
|---|---|---|---|---|---|---|---|---|---|---|---|---|---|---|---|---|---|---|---|---|
| fig\|693216 RcsB | Cronobacter turicensis z3032 | 693216,3 | FN543093 | Ctu_28740 | VBICroTur | PATRIC.69 | PATRIC | CDS | 3001704 | 3002354 | 651 | + | FIG000041 | PLF_4134 | PGF_0042 | CBA32355 | 216 | rcsB | DNA-binding capsular synthesis response regulator RcsB | |
| fig\|693216 RcsC | Cronobacter turicensis z3032 | 693216,3 | FN543093 | Ctu_28750 | VBICroTur | PATRIC.69 | PATRIC | CDS | 3002533 | 3005382 | 2850 | - | FIG000042 | PLF_4134 | PGF_0005 | CBA32357 | 949 | rcsC | Sensor hist | GO:0004673\|protein histidine kinase activity |
| fig\|693216 IgaA | Cronobacter turicensis z3032 | 693216,3 | FN543093 | Ctu_39070 | VBICroTur | PATRIC.69 | PATRIC | CDS | 4051923 | 4054031 | 2109 | + | FIG000040 | PLF_4134 | PGF_0001 | CBA34231 | 702 | yrfF | IgaA: a membrane protein that prevents overactivation of the Rcs regulatory system | |
| fig\|693216 RcsF | Cronobacter turicensis z3032 | 693216,3 | FN543093 | Ctu_08260 | VBICroTur | PATRIC.69 | PATRIC | CDS | 894274 | 894678 | 405 | - | FIG01304C | PLF_4134 | PGF_0003 | CBA28233 | 134 | rcsF | Protein RcsF | |
| fig\|107400 DnaK | Cronobacter universalis NCTC 9529 | 1074000.3 | CAKX01000221 | BN136_24 | VBICroUni | PATRIC.10 | PATRIC | CDS | 12497 | 13927 | 1431 | + | FIG000233 | PLF_4134 | PGF_1035 | CCK14233 | 476 | | Chaperone protein DnaK | |
| fig\|107400 RcsD | Cronobacter universalis NCTC 9529 | 1074000,3 | CAKX01000221 | BN136_12 | VBICroUni | PATRIC.10 | PATRIC | CDS | 2292614 | 2295439 | 2826 | + | FIG000045 | PLF_4134 | PGF_0003 | CCK15253 | 941 | | Phosphotransferase RcsD | |
| fig\|107400 RcsB | Cronobacter universalis NCTC 9529 | 1074000,3 | CAKX01000221 | BN136_12 | VBICroUni | PATRIC.10 | PATRIC | CDS | 2295456 | 2296106 | 651 | + | FIG000041 | PLF_4134 | PGF_0042 | CCK15254 | 216 | | DNA-binding capsular synthesis response regulator RcsB | |
| fig\|107400 RcsC | Cronobacter universalis NCTC 9529 | 1074000,3 | CAKX01000221 | BN136_12 | VBICroUni | PATRIC.10 | PATRIC | CDS | 2296359 | 2299205 | 2847 | - | FIG000042 | PLF_4134 | PGF_0005 | CCK15255 | 948 | | Sensor hist | GO:0004673\|protein histidine kinase activity |
| fig\|107400 RcsC | Cronobacter universalis NCTC 9529 | 1074000,3 | CAKX01000221 | BN136_12 | VBICroUni | PATRIC.10 | PATRIC | CDS | 2299171 | 2299641 | 471 | - | FIG000042 | PLF_4134 | PGF_0005 | CCK15256 | 156 | | Sensor hist | GO:0004673\|protein histidine kinase activity |
| fig\|107400 IgaA | Cronobacter universalis NCTC 9529 | 1074000,3 | CAKX01000221 | BN136_23 | VBICroUni | PATRIC.10 | PATRIC | CDS | 3300680 | 3302788 | 2109 | + | FIG000040 | PLF_4134 | PGF_0001 | CCK16301 | 702 | | IgaA: a membrane protein that prevents overactivation of the Rcs regulatory system | |
| fig\|107400 RcsF | Cronobacter universalis NCTC 9529 | 1074000,3 | CAKX01000221 | BN136_47 | VBICroUni | PATRIC.10 | PATRIC | CDS | 212076 | 212480 | 405 | - | FIG01304C | PLF_4134 | PGF_0003 | CCK14463 | 134 | | Protein RcsF | |
| fig\|138874 DnaK | Cronobacter zurichensis LMG 23730 | 1388748.3 | AWFZ01000017 | | VBICroZur | PATRIC.13 | PATRIC | CDS | 195582 | 197498 | 1917 | - | FIG00023369 | | PGF_10357457 | | 638 | | Chaperone protein DnaK | |
| fig\|138874 RcsD | Cronobacter zurichensis LMG 23730 | 1388748,3 | AWFZ01000030 | | VBICroZur | PATRIC.13 | PATRIC | CDS | 204660 | 207311 | 2652 | + | FIG00004592 | | PGF_00034034 | | 883 | | Phosphotransferase RcsD | |
| fig\|138874 RcsB | Cronobacter zurichensis LMG 23730 | 1388748,3 | AWFZ01000030 | | VBICroZur | PATRIC.13 | PATRIC | CDS | 207328 | 207978 | 651 | + | FIG00004153 | | PGF_00421954 | | 216 | | DNA-binding capsular synthesis response regulator RcsB | |
| fig\|138874 RcsC | Cronobacter zurichensis LMG 23730 | 1388748,3 | AWFZ01000030 | | VBICroZur | PATRIC.13 | PATRIC | CDS | 208027 | 210876 | 2850 | - | FIG00004208 | | PGF_00050577 | | 949 | | Sensor hist | GO:0004673\|protein histidine kinase activity |
| fig\|138874 RcsF | Cronobacter zurichensis LMG 23730 | 1388748,3 | AWFZ01000017 | | VBICroZur | PATRIC.13 | PATRIC | CDS | 3940 | 4200 | 261 | + | FIG01304060 | | PGF_00037945 | | 86 | | Protein RcsF | |
| fig\|138874 IgaA | Cronobacter zurichensis LMG 23730 | 1388748,3 | AWFZ01000061 | | VBICroZur | PATRIC.13 | PATRIC | CDS | 333064 | 334971 | 1908 | - | FIG00004077 | | PGF_00013790 | | 635 | | IgaA: a membrane protein that prevents overactivation of the Rcs regulatory system | |
| fig\|198628 DnaK | Dickeya dadantii 3937 | 198628.6 | NC_014500 | Dda3937_ | VBIDicDad | PATRIC.19 | PATRIC | CDS | 4195728 | 4197638 | 1911 | - | FIG000233 | PLF_2040 | PGF_1035 | YP_00388 | 636 | dnaK | Chaperone protein DnaK | |
| fig\|198628 RcsC | Dickeya dadantii 3937 | 198628,6 | NC_014500 | Dda3937_ | VBIDicDad | PATRIC.19 | PATRIC | CDS | 1322946 | 1325801 | 2856 | + | FIG000042 | PLF_2040 | PGF_0005 | YP_00388 | 951 | rcsC | Sensor hist | GO:0004673\|protein histidine kinase activity |
| fig\|198628 RcsB | Dickeya dadantii 3937 | 198628,6 | NC_014500 | Dda3937_ | VBIDicDad | PATRIC.19 | PATRIC | CDS | 1325862 | 1326512 | 651 | - | FIG000041 | PLF_2040 | PGF_0042 | YP_00388 | 216 | rcsB | DNA-binding capsular synthesis response regulator RcsB | |
| fig\|198628 RcsD | Dickeya dadantii 3937 | 198628,6 | NC_014500 | Dda3937_ | VBIDicDad | PATRIC.19 | PATRIC | CDS | 1326519 | 1329188 | 2670 | - | FIG000045 | PLF_2040 | PGF_0003 | YP_00388 | 889 | yojN | Phosphotransferase RcsD | |
| fig\|198628 RcsF | Dickeya dadantii 3937 | 198628,6 | NC_014500 | Dda3937_ | VBIDicDad | PATRIC.19 | PATRIC | CDS | 3833633 | 3834040 | 408 | + | FIG01304C | PLF_2040 | PGF_0003 | YP_00388 | 135 | rcsF | Protein RcsF | |
| fig\|198628 IgaA | Dickeya dadantii 3937 | 198628,6 | NC_014500 | Dda3937_ | VBIDicDad | PATRIC.19 | PATRIC | CDS | 4454965 | 4457094 | 2130 | + | FIG000040 | PLF_2040 | PGF_0001 | YP_00388 | 709 | yrfF | IgaA: a membrane protein that prevents overactivation of the Rcs regulatory system | |
| fig\|122415 DnaK | Dickeya paradisiaca NCPPB 2511 | 1224150.8 | CM001857 | | | PATRIC.12 | PATRIC | CDS | 641178 | 643097 | 1920 | + | FIG000233 | PLF_2040 | PGF_10357457 | | 639 | | Chaperone protein DnaK | |
| fig\|122415 RcsC | Dickeya paradisiaca NCPPB 2511 | 1224150,8 | CM001857 | | | PATRIC.12 | PATRIC | CDS | 1175848 | 1178700 | 2853 | + | FIG000042 | PLF_2040 | PGF_00050577 | | 950 | | Sensor hist | GO:0004673\|protein histidine kinase activity |
| fig\|122415 RcsB | Dickeya paradisiaca NCPPB 2511 | 1224150,8 | CM001857 | | | PATRIC.12 | PATRIC | CDS | 1178759 | 1179415 | 657 | - | FIG000041 | PLF_2040 | PGF_00421954 | | 218 | | DNA-binding capsular synthesis response regulator RcsB | |
| fig\|122415 RcsD | Dickeya paradisiaca NCPPB 2511 | 1224150,8 | CM001857 | | | PATRIC.12 | PATRIC | CDS | 1179422 | 1182091 | 2670 | - | FIG000045 | PLF_2040 | PGF_00034034 | | 889 | | Phosphotransferase RcsD | |
| fig\|122415 IgaA | Dickeya paradisiaca NCPPB 2511 | 1224150,8 | CM001857 | | | PATRIC.12 | PATRIC | CDS | 372216 | 374360 | 2145 | - | FIG000040 | PLF_2040 | PGF_00013790 | | 714 | | IgaA: a membrane protein that prevents overactivation of the Rcs regulatory system | |
| fig\|122415 RcsF | Dickeya paradisiaca NCPPB 2511 | 1224150,8 | CM001857 | | | PATRIC.12 | PATRIC | CDS | 986878 | 987285 | 408 | - | FIG01304C | PLF_2040 | PGF_00037945 | | 135 | | Protein RcsF | |
| fig\|561229 DnaK | Dickeya zeae Ech1591 | 561229.3 | NC_012912 | Dd1591_0 | VBIDicZea | PATRIC.56 | PATRIC | CDS | 616690 | 618600 | 1911 | + | FIG000233 | PLF_2040 | PGF_1035 | YP_00300 | 636 | | Chaperone protein DnaK | |
| fig\|561229 IgaA | Dickeya zeae Ech1591 | 561229,3 | NC_012912 | Dd1591_0 | VBIDicZea | PATRIC.56 | PATRIC | CDS | 303835 | 305970 | 2136 | - | FIG000040 | PLF_2040 | PGF_0001 | YP_00300 | 711 | | IgaA: a membrane protein that prevents overactivation of the Rcs regulatory system | |
| fig\|561229 RcsD | Dickeya zeae Ech1591 | 561229,3 | NC_012912 | Dd1591_3 | VBIDicZea | PATRIC.56 | PATRIC | CDS | 3431059 | 3433728 | 2670 | + | FIG000045 | PLF_2040 | PGF_0003 | YP_00300 | 889 | | Phosphotransferase RcsD | |
| fig\|561229 RcsB | Dickeya zeae Ech1591 | 561229,3 | NC_012912 | Dd1591_3 | VBIDicZea | PATRIC.56 | PATRIC | CDS | 3433734 | 3434384 | 651 | + | FIG000041 | PLF_2040 | PGF_0042 | YP_00300 | 216 | | DNA-binding capsular synthesis response regulator RcsB | |
| fig\|561229 RcsC | Dickeya zeae Ech1591 | 561229,3 | NC_012912 | Dd1591_3 | VBIDicZea | PATRIC.56 | PATRIC | CDS | 3434445 | 3437342 | 2898 | - | FIG000042 | PLF_2040 | PGF_0005 | YP_00300 | 965 | | Sensor hist | GO:0004673\|protein histidine kinase activity |
| fig\|561229 RcsF | Dickeya zeae Ech1591 | 561229,3 | NC_012912 | Dd1591_0 | VBIDicZea | PATRIC.56 | PATRIC | CDS | 978302 | 978559 | 258 | - | FIG01304C | PLF_2040 | PGF_0003 | YP_00300 | 85 | | Protein RcsF | |
| fig\|634503 DnaK | Edwardsiella ictaluri 93-146 | 634503.3 | NC_012779 | NT01EI_06 | VBIEdwIct | PATRIC.63 | PATRIC | CDS | 657138 | 659045 | 1908 | + | FIG000233 | PLF_635_ | PGF_1035 | YP_00293 | 635 | | Chaperone protein DnaK | |
| fig\|634503 RcsD | Edwardsiella ictaluri 93-146 | 634503,3 | NC_012779 | NT01EI_26 | VBIEdwIct | PATRIC.63 | PATRIC | CDS | 2515147 | 2517828 | 2682 | + | FIG000045 | PLF_635_ | PGF_0003 | YP_00293 | 893 | | Phosphotransferase RcsD | |
| fig\|634503 RcsB | Edwardsiella ictaluri 93-146 | 634503,3 | NC_012779 | NT01EI_26 | VBIEdwIct | PATRIC.63 | PATRIC | CDS | 2517821 | 2518474 | 654 | + | FIG000041 | PLF_635_ | PGF_0042 | YP_00293 | 217 | | DNA-binding capsular synthesis response regulator RcsB | |
| fig\|634503 RcsC | Edwardsiella ictaluri 93-146 | 634503,3 | NC_012779 | NT01EI_26 | VBIEdwIct | PATRIC.63 | PATRIC | CDS | 2518506 | 2521460 | 2955 | - | FIG000042 | PLF_635_ | PGF_0005 | YP_00293 | 984 | | Sensor hist | GO:0004673\|protein histidine kinase activity |
| fig\|634503 IgaA | Edwardsiella ictaluri 93-146 | 634503,3 | NC_012779 | NT01EI_36 | VBIEdwIct | PATRIC.63 | PATRIC | CDS | 3519949 | 3521970 | 2022 | + | FIG000040 | PLF_635_ | PGF_0001 | YP_00293 | 673 | | IgaA: a membrane protein that prevents overactivation of the Rcs regulatory system | |
| fig\|634503 RcsF | Edwardsiella ictaluri 93-146 | 634503,3 | NC_012779 | NT01EI_06 | VBIEdwIct | PATRIC.63 | PATRIC | CDS | 618377 | 618799 | 423 | - | FIG01304C | PLF_635_ | PGF_0003 | YP_00293 | 140 | | Protein RcsF | |
| fig\|498217 DnaK | Edwardsiella tarda EIB202 | 498217.4 | NC_013508 | ETAE_057 | VBIEdwTa | PATRIC.49 | PATRIC | CDS | 631172 | 633079 | 1908 | + | FIG000233 | PLF_635_ | PGF_1035 | YP_00329 | 635 | dnaK | Chaperone protein DnaK | |
| fig\|498217 RcsD | Edwardsiella tarda EIB202 | 498217,4 | NC_013508 | ETAE_233 | VBIEdwTa | PATRIC.49 | PATRIC | CDS | 2451108 | 2453804 | 2697 | + | FIG000045 | PLF_635_ | PGF_0003 | YP_00329 | 898 | | Phosphotransferase RcsD | |
| fig\|498217 RcsB | Edwardsiella tarda EIB202 | 498217,4 | NC_013508 | ETAE_233 | VBIEdwTa | PATRIC.49 | PATRIC | CDS | 2453797 | 2454450 | 654 | + | FIG000041 | PLF_635_ | PGF_0042 | YP_00329 | 217 | | DNA-binding capsular synthesis response regulator RcsB | |
| fig\|498217 RcsC | Edwardsiella tarda EIB202 | 498217,4 | NC_013508 | ETAE_233 | VBIEdwTa | PATRIC.49 | PATRIC | CDS | 2454483 | 2457353 | 2871 | - | FIG000042 | PLF_635_ | PGF_0005 | YP_00329 | 956 | | Sensor hist | GO:0004673\|protein histidine kinase activity |
| fig\|498217 IgaA | Edwardsiella tarda EIB202 | 498217,4 | NC_013508 | ETAE_326 | VBIEdwTa | PATRIC.49 | PATRIC | CDS | 3439751 | 3441772 | 2022 | + | FIG000040 | PLF_635_ | PGF_0001 | YP_00329 | 673 | | IgaA: a membrane protein that prevents overactivation of the Rcs regulatory system | |
| fig\|498217 RcsF | Edwardsiella tarda EIB202 | 498217,4 | NC_013508 | ETAE_055 | VBIEdwTa | PATRIC.49 | PATRIC | CDS | 596827 | 597249 | 423 | - | FIG01304C | PLF_635_ | PGF_0003 | YP_00329 | 140 | rcsF | Protein RcsF | |
| fig\|102830 DnaK | Enterobacter aerogenes KCTC 2190 | 1028307.3 | NC_015663 | EAE_1073 | VBIEntAer | PATRIC.10 | PATRIC | CDS | 2286596 | 2288512 | 1917 | + | FIG000233 | PLF_570_ | PGF_1035 | YP_00459 | 638 | dnaK | Chaperone protein DnaK | |
| fig\|102830 IgaA | Enterobacter aerogenes KCTC 2190 | 1028307,3 | NC_015663 | EAE_0521 | VBIEntAer | PATRIC.10 | PATRIC | CDS | 1069090 | 1071225 | 2136 | + | FIG000040 | PLF_570_ | PGF_0001 | YP_00459 | 711 | | IgaA: a membrane protein that prevents overactivation of the Rcs regulatory system | |
| fig\|102830 RcsF | Enterobacter aerogenes KCTC 2190 | 1028307,3 | NC_015663 | EAE_1178 | VBIEntAer | PATRIC.10 | PATRIC | CDS | 2532803 | 2533159 | 357 | - | FIG01304C | PLF_570_ | PGF_0003 | YP_00459 | 118 | rcsF | Protein RcsF | |
| fig\|102830 RcsD | Enterobacter aerogenes KCTC 2190 | 1028307,3 | NC_015663 | EAE_2421 | VBIEntAer | PATRIC.10 | PATRIC | CDS | 5155786 | 5158389 | 2604 | + | FIG000045 | PLF_570_ | PGF_0003 | YP_00459 | 867 | | Phosphotransferase RcsD | |
| fig\|102830 RcsB | Enterobacter aerogenes KCTC 2190 | 1028307,3 | NC_015663 | EAE_2422 | VBIEntAer | PATRIC.10 | PATRIC | CDS | 5158406 | 5159056 | 651 | + | FIG000041 | PLF_570_ | PGF_0042 | YP_00459 | 216 | | DNA-binding capsular synthesis response regulator RcsB | |
| fig\|102830 RcsC | Enterobacter aerogenes KCTC 2190 | 1028307,3 | NC_015663 | EAE_2422 | VBIEntAer | PATRIC.10 | PATRIC | CDS | 5159169 | 5162021 | 2853 | - | FIG000042 | PLF_570_ | PGF_0005 | YP_00459 | 950 | | Sensor hist | GO:0004673\|protein histidine kinase activity |
| fig\|640513 DnaK | Enterobacter asburiae LF7a | 640513.3 | NC_015968 | Entas_062 | VBIEntAsb | PATRIC.64 | PATRIC | CDS | 667932 | 669845 | 1914 | + | FIG000233 | PLF_547_ | PGF_1035 | YP_00482 | 637 | | Chaperone protein DnaK | |
| fig\|640513 RcsD | Enterobacter asburiae LF7a | 640513,3 | NC_015968 | Entas_296 | VBIEntAsb | PATRIC.64 | PATRIC | CDS | 3181614 | 3184391 | 2778 | + | FIG000045 | PLF_547_ | PGF_0003 | YP_00482 | 925 | | Phosphotransferase RcsD | |
| fig\|640513 RcsB | Enterobacter asburiae LF7a | 640513,3 | NC_015968 | Entas_296 | VBIEntAsb | PATRIC.64 | PATRIC | CDS | 3184408 | 3185058 | 651 | + | FIG000041 | PLF_547_ | PGF_0042 | YP_00482 | 216 | | DNA-binding capsular synthesis response regulator RcsB | |
| fig\|640513 RcsC | Enterobacter asburiae LF7a | 640513,3 | NC_015968 | Entas_296 | VBIEntAsb | PATRIC.64 | PATRIC | CDS | 3185134 | 3187911 | 2778 | - | FIG000042 | PLF_547_ | PGF_0005 | YP_00482 | 925 | | Sensor hist | GO:0004673\|protein histidine kinase activity |
| fig\|640513 IgaA | Enterobacter asburiae LF7a | 640513,3 | NC_015968 | Entas_409 | VBIEntAsb | PATRIC.64 | PATRIC | CDS | 4365875 | 4368013 | 2139 | + | FIG000040 | PLF_547_ | PGF_0001 | YP_00483 | 712 | | IgaA: a membrane protein that prevents overactivation of the Rcs regulatory system | |
| fig\|640513 RcsF | Enterobacter asburiae LF7a | 640513,3 | NC_015968 | Entas_079 | VBIEntAsb | PATRIC.64 | PATRIC | CDS | 860937 | 861341 | 405 | - | FIG01304C | PLF_547_ | PGF_0003 | YP_00482 | 134 | | Protein RcsF | |
| fig\|500639 DnaK | Enterobacter cancerogenus ATCC 35316 | 500639.8 | NZ_GG704863 | EcanA3_0 | VBIEntCan | PATRIC.50 | PATRIC | CDS | 210548 | 212461 | 1914 | - | FIG000233 | PLF_547_ | PGF_1035 | ZP_05966 | 637 | dnaK | Chaperone protein DnaK | |
| fig\|500639 IgaA | Enterobacter cancerogenus ATCC 35316 | 500639,8 | NZ_GG704865 | EcanA3_0 | VBIEntCan | PATRIC.50 | PATRIC | CDS | 52071 | 54209 | 2139 | + | FIG000040 | PLF_547_ | PGF_0001 | ZP_05969 | 712 | | IgaA: a membrane protein that prevents overactivation of the Rcs regulatory system | |
| fig\|500639 RcsF | Enterobacter cancerogenus ATCC 35316 | 500639,8 | NZ_GG704863 | EcanA3_0 | VBIEntCan | PATRIC.50 | PATRIC | CDS | 5573 | 5926 | 354 | + | FIG01304C | PLF_547_ | PGF_0003 | ZP_05966 | 117 | rcsF | Protein RcsF | |
| fig\|500639 RcsD | Enterobacter cancerogenus ATCC 35316 | 500639,8 | NZ_GG704863 | EcanA3_0 | VBIEntCan | PATRIC.50 | PATRIC | CDS | 1944181 | 1946757 | 2577 | + | FIG000045 | PLF_547_ | PGF_0003 | ZP_05968 | 858 | | Phosphotransferase RcsD | |
| fig\|500639 RcsB | Enterobacter cancerogenus ATCC 35316 | 500639,8 | NZ_GG704863 | EcanA3_0 | VBIEntCan | PATRIC.50 | PATRIC | CDS | 1946774 | 1947424 | 651 | + | FIG000041 | PLF_547_ | PGF_0042 | ZP_05968 | 216 | | DNA-binding capsular synthesis response regulator RcsB | |
| fig\|500639 RcsC | Enterobacter cancerogenus ATCC 35316 | 500639,8 | NZ_GG704863 | EcanA3_0 | VBIEntCan | PATRIC.50 | PATRIC | CDS | 1947536 | 1950382 | 2847 | - | FIG000042 | PLF_547_ | PGF_0005 | ZP_05968 | 948 | | Sensor hist | GO:0004673\|protein histidine kinase activity |
| fig\|716541 DnaK | Enterobacter cloacae subsp. cloacae ATCC 13047 | 716541.4 | NC_014121 | ECL_0082 | VBIEntClo | PATRIC.71 | PATRIC | CDS | 816378 | 818291 | 1914 | + | FIG000233 | PLF_547_ | PGF_1035 | YP_00361 | 637 | dnaK | Chaperone protein DnaK | |
| fig\|716541 RcsF | Enterobacter cloacae subsp. cloacae ATCC 13047 | 716541,4 | NC_014121 | ECL_0100 | VBIEntClo | PATRIC.71 | PATRIC | CDS | 1011586 | 1011816 | 231 | - | FIG01304C | PLF_547_ | PGF_0003 | YP_00361 | 76 | | Protein RcsF | |
| fig\|716541 RcsD | Enterobacter cloacae subsp. cloacae ATCC 13047 | 716541,4 | NC_014121 | ECL_0352 | VBIEntClo | PATRIC.71 | PATRIC | CDS | 3603890 | 3606454 | 2565 | + | FIG000045 | PLF_547_ | PGF_0003 | YP_00361 | 854 | rcsD | Phosphotransferase RcsD | |
| fig\|716541 RcsB | Enterobacter cloacae subsp. cloacae ATCC 13047 | 716541,4 | NC_014121 | ECL_0352 | VBIEntClo | PATRIC.71 | PATRIC | CDS | 3606471 | 3607121 | 651 | + | FIG000041 | PLF_547_ | PGF_0042 | YP_00361 | 216 | rcsB | DNA-binding capsular synthesis response regulator RcsB | |
| fig\|716541 RcsC | Enterobacter cloacae subsp. cloacae ATCC 13047 | 716541,4 | NC_014121 | ECL_0352 | VBIEntClo | PATRIC.71 | PATRIC | CDS | 3607211 | 3610057 | 2847 | - | FIG000042 | PLF_547_ | PGF_0005 | YP_00361 | 948 | rcsC | Sensor hist | GO:0004673\|protein histidine kinase activity |
| fig\|716541 IgaA | Enterobacter cloacae subsp. cloacae ATCC 13047 | 716541,4 | NC_014121 | ECL_0476 | VBIEntClo | PATRIC.71 | PATRIC | CDS | 4869325 | 4871463 | 2139 | + | FIG000040 | PLF_547_ | PGF_0001 | YP_00361 | 712 | | IgaA: a membrane protein that prevents overactivation of the Rcs regulatory system | |
| fig\|888063 DnaK | Enterobacter hormaechei ATCC 49162 | 888063.3 | NZ_AFHR01000027 | HMPREF9 | VBIEntHor | PATRIC.88 | PATRIC | CDS | 202449 | 204362 | 1914 | - | FIG000233 | PLF_547_ | PGF_1035 | ZP_08497 | 637 | dnaK | Chaperone protein DnaK | |
| fig\|888063 RcsF | Enterobacter hormaechei ATCC 49162 | 888063,3 | NZ_AFHR01000027 | HMPREF9 | VBIEntHor | PATRIC.88 | PATRIC | CDS | 3464 | 3868 | 405 | + | FIG01304C | PLF_547_ | PGF_0003 | ZP_08497 | 134 | rcsF | Protein RcsF | |
| fig\|888063 RcsD | Enterobacter hormaechei ATCC 49162 | 888063,3 | NZ_AFHR01000057 | HMPREF9 | VBIEntHor | PATRIC.88 | PATRIC | CDS | 150217 | 152790 | 2574 | + | FIG000045 | PLF_547_ | PGF_0003 | ZP_08498 | 857 | yojN | Phosphotransferase RcsD | |
| fig\|888063 RcsB | Enterobacter hormaechei ATCC 49162 | 888063,3 | NZ_AFHR01000057 | HMPREF9 | VBIEntHor | PATRIC.88 | PATRIC | CDS | 152807 | 153457 | 651 | + | FIG000041 | PLF_547_ | PGF_0042 | ZP_08498 | 216 | rcsB | DNA-binding capsular synthesis response regulator RcsB | |
| fig\|888063 RcsC | Enterobacter hormaechei ATCC 49162 | 888063,3 | NZ_AFHR01000057 | HMPREF9 | VBIEntHor | PATRIC.88 | PATRIC | CDS | 153570 | 156416 | 2847 | - | FIG000042 | PLF_547_ | PGF_0005 | ZP_08498 | 948 | rcsC | Sensor hist | GO:0004673\|protein histidine kinase activity |
| fig\|888063 IgaA | Enterobacter hormaechei ATCC 49162 | 888063,3 | NZ_AFHR01000069 | HMPREF9 | VBIEntHor | PATRIC.88 | PATRIC | CDS | 49279 | 51417 | 2139 | + | FIG000040 | PLF_547_ | PGF_0001 | ZP_08499 | 712 | igaA | IgaA: a membrane protein that prevents overactivation of the Rcs regulatory system | |
| fig\|888063 RcsB | Enterobacter hormaechei ATCC 49162 | 888063,3 | NZ_AFHR01000073 | HMPREF9 | VBIEntHor | PATRIC.88 | PATRIC | CDS | 172743 | 173408 | 666 | - | FIG013087 | PLF_547_ | PGF_0822 | ZP_08500 | 221 | rcsB2 | hypothetical protein | |
| fig\|701347 DnaK | Enterobacter lignolyticus SCF1 | 701347.4 | NC_014618 | Entcl_3705 | VBIEntClo | PATRIC.70 | PATRIC | CDS | 3982710 | 3984626 | 1917 | - | FIG000233 | PLF_1330 | PGF_1035 | YP_00394 | 638 | | Chaperone protein DnaK | |
| fig\|701347 RcsC | Enterobacter lignolyticus SCF1 | 701347,4 | NC_014618 | Entcl_1499 | VBIEntClo | PATRIC.70 | PATRIC | CDS | 1583927 | 1586776 | 2850 | + | FIG000042 | PLF_1330 | PGF_0005 | YP_00394 | 949 | | Sensor hist | GO:0004673\|protein histidine kinase activity |
| fig\|701347 RcsB | Enterobacter lignolyticus SCF1 | 701347,4 | NC_014618 | Entcl_1500 | VBIEntClo | PATRIC.70 | PATRIC | CDS | 1586830 | 1587480 | 651 | - | FIG000041 | PLF_1330 | PGF_0042 | YP_00394 | 216 | | DNA-binding capsular synthesis response regulator RcsB | |
| fig\|701347 RcsD | Enterobacter lignolyticus SCF1 | 701347,4 | NC_014618 | Entcl_1501 | VBIEntClo | PATRIC.70 | PATRIC | CDS | 1587497 | 1590172 | 2676 | - | FIG000045 | PLF_1330 | PGF_0003 | YP_00394 | 891 | | Phosphotransferase RcsD | |
| fig\|701347 IgaA | Enterobacter lignolyticus SCF1 | 701347,4 | NC_014618 | Entcl_0336 | VBIEntClo | PATRIC.70 | PATRIC | CDS | 369964 | 372105 | 2142 | - | FIG000040 | PLF_1330 | PGF_0001 | YP_00393 | 713 | | IgaA: a membrane protein that prevents overactivation of the Rcs regulatory system | |
| fig\|701347 RcsF | Enterobacter lignolyticus SCF1 | 701347,4 | NC_014618 | Entcl_3539 | VBIEntClo | PATRIC.70 | PATRIC | CDS | 3793000 | 3793404 | 405 | + | FIG01304C | PLF_1330 | PGF_0003 | YP_00394 | 134 | | Protein RcsF | |
| fig\|112186 DnaK | Enterobacter pulveris DSM 19144 | 1121863.3 | JHYZ01000006 | | | PATRIC.11 | PATRIC | CDS | 257784 | 259700 | 1917 | + | FIG000233 | PLF_1649 | PGF_10357457 | | 638 | | Chaperone protein DnaK | |
| fig\|112186 RcsF | Enterobacter pulveris DSM 19144 | 1121863,3 | JHYZ01000006 | | | PATRIC.11 | PATRIC | CDS | 448743 | 448973 | 231 | - | FIG01304C | PLF_1649 | PGF_00037945 | | 76 | | Protein RcsF | |

| | | | | | | | | | | | | | | | | | | | | | |
|---|---|---|---|---|---|---|---|---|---|---|---|---|---|---|---|---|---|---|---|---|---|
| fig\|112186 | RcsD | Enterobacter pulveris DSM 19144 | 1121863,3 | JHYZ01000010 | | | PATRIC.11 | PATRIC | CDS | 30272 | 32941 | 2670 | + | FIG000045 | PLF_1649 | PGF_00034034 | | 889 | | Phosphotransferase RcsD | |
| fig\|112186 | RcsB | Enterobacter pulveris DSM 19144 | 1121863,3 | JHYZ01000010 | | | PATRIC.11 | PATRIC | CDS | 32958 | 33608 | 651 | + | FIG000041 | PLF_1649 | PGF_00421954 | | 216 | | DNA-binding capsular synthesis response regulator RcsB | |
| fig\|112186 | RcsC | Enterobacter pulveris DSM 19144 | 1121863,3 | JHYZ01000010 | | | PATRIC.11 | PATRIC | CDS | 33687 | 36536 | 2850 | - | FIG000042 | PLF_1649 | PGF_00050577 | | 949 | | Sensor hist | GO:0004673\|protein histidine kinase activity |
| fig\|112186 | IgaA | Enterobacter pulveris DSM 19144 | 1121863,3 | JHYZ01000014 | | | PATRIC.11 | PATRIC | CDS | 51575 | 53713 | 2139 | + | FIG000040 | PLF_1649 | PGF_00013790 | | 712 | | IgaA: a membrane protein that prevents overactivation of the Rcs regulatory system | |
| fig\|716540 | DnaK | Erwinia amylovora ATCC 49946 | 716540.3 | NC_013971 | EAM_0649 | VBIErwAm | PATRIC.71 | PATRIC | CDS | 727747 | 729660 | 1914 | + | FIG000233 | PLF_551_ | PGF_1035 | YP_00353 | 637 | dnaK | Chaperone protein DnaK | |
| fig\|716540 | RcsD | Erwinia amylovora ATCC 49946 | 716540,3 | NC_013971 | EAM_2261 | VBIErwAm | PATRIC.71 | PATRIC | CDS | 2439304 | 2441961 | 2658 | + | FIG000045 | PLF_551_ | PGF_0003 | YP_00353 | 885 | | Phosphotransferase RcsD | |
| fig\|716540 | RcsB | Erwinia amylovora ATCC 49946 | 716540,3 | NC_013971 | EAM_2262 | VBIErwAm | PATRIC.71 | PATRIC | CDS | 2441963 | 2442610 | 648 | + | FIG000041 | PLF_551_ | PGF_0042 | YP_00353 | 215 | rcsB | DNA-binding capsular synthesis response regulator RcsB | |
| fig\|716540 | RcsC | Erwinia amylovora ATCC 49946 | 716540,3 | NC_013971 | EAM_2263 | VBIErwAm | PATRIC.71 | PATRIC | CDS | 2442740 | 2445541 | 2802 | - | FIG000042 | PLF_551_ | PGF_0005 | YP_00353 | 933 | rcsC | Sensor hist | GO:0004673\|protein histidine kinase activity |
| fig\|716540 | IgaA | Erwinia amylovora ATCC 49946 | 716540,3 | NC_013971 | EAM_3248 | VBIErwAm | PATRIC.71 | PATRIC | CDS | 3532541 | 3534649 | 2109 | + | FIG000040 | PLF_551_ | PGF_0001 | YP_00354 | 702 | | IgaA: a membrane protein that prevents overactivation of the Rcs regulatory system | |
| fig\|716540 | RcsF | Erwinia amylovora ATCC 49946 | 716540,3 | NC_013971 | EAM_0846 | VBIErwAm | PATRIC.71 | PATRIC | CDS | 953179 | 953583 | 405 | - | FIG013040 | PLF_551_ | PGF_0003 | YP_00353 | 134 | rcsF | Protein RcsF | |
| fig\|665029 | DnaK | Erwinia amylovora CFBP1430 | 665029.3 | NC_013961 | EAMY_294 | VBIErwAm | PATRIC.66 | PATRIC | CDS | 3041722 | 3043635 | 1914 | - | FIG000233 | PLF_551_ | PGF_1035 | YP_00353 | 637 | dnaK | Chaperone protein DnaK | |
| fig\|665029 | RcsD | Erwinia amylovora CFBP1430 | 665029,3 | NC_013961 | EAMY_234 | VBIErwAm | PATRIC.66 | PATRIC | CDS | 2401776 | 2404433 | 2658 | + | FIG000045 | PLF_551_ | PGF_0003 | YP_00353 | 885 | yojN | Phosphotransferase RcsD | |
| fig\|665029 | RcsB | Erwinia amylovora CFBP1430 | 665029,3 | NC_013961 | EAMY_234 | VBIErwAm | PATRIC.66 | PATRIC | CDS | 2404435 | 2405082 | 648 | + | FIG000041 | PLF_551_ | PGF_0042 | YP_00353 | 215 | rcsB3 | DNA-binding capsular synthesis response regulator RcsB | |
| fig\|665029 | RcsC | Erwinia amylovora CFBP1430 | 665029,3 | NC_013961 | EAMY_234 | VBIErwAm | PATRIC.66 | PATRIC | CDS | 2405212 | 2408103 | 2892 | - | FIG000042 | PLF_551_ | PGF_0005 | YP_00353 | 963 | rcsC1 | Sensor hist | GO:0004673\|protein histidine kinase activity |
| fig\|665029 | RcsF | Erwinia amylovora CFBP1430 | 665029,3 | NC_013961 | EAMY_273 | VBIErwAm | PATRIC.66 | PATRIC | CDS | 2817616 | 2818020 | 405 | + | FIG013040 | PLF_551_ | PGF_0003 | YP_00353 | 134 | rcsF | Protein RcsF | |
| fig\|665029 | IgaA | Erwinia amylovora CFBP1430 | 665029,3 | NC_013961 | EAMY_343 | VBIErwAm | PATRIC.66 | PATRIC | CDS | 3532289 | 3534397 | 2109 | + | FIG000040 | PLF_551_ | PGF_0001 | YP_00353 | 702 | yrfF | IgaA: a membrane protein that prevents overactivation of the Rcs regulatory system | |
| fig\|634500 | DnaK | Erwinia billingiae Eb661 | 634500.5 | NC_014306 | EbC_0668 | VBIErwBil9 | PATRIC.63 | PATRIC | CDS | 801028 | 802944 | 1917 | + | FIG000233 | PLF_551_ | PGF_1035 | YP_00374 | 638 | dnaK | Chaperone protein DnaK | |
| fig\|634500 | RcsF | Erwinia billingiae Eb661 | 634500,5 | NC_014306 | EbC_0835 | VBIErwBil9 | PATRIC.63 | PATRIC | CDS | 1000564 | 1000914 | 351 | - | FIG013040 | PLF_551_ | PGF_0003 | YP_00374 | 116 | rcsF | Protein RcsF | |
| fig\|634500 | RcsD | Erwinia billingiae Eb661 | 634500,5 | NC_014306 | EbC_3037 | VBIErwBil9 | PATRIC.63 | PATRIC | CDS | 3408515 | 3411175 | 2661 | + | FIG000045 | PLF_551_ | PGF_0003 | YP_00374 | 886 | rcsD | Phosphotransferase RcsD | |
| fig\|634500 | RcsB | Erwinia billingiae Eb661 | 634500,5 | NC_014306 | EbC_3038 | VBIErwBil9 | PATRIC.63 | PATRIC | CDS | 3411177 | 3411827 | 651 | + | FIG000041 | PLF_551_ | PGF_0042 | YP_00374 | 216 | rcsB | DNA-binding capsular synthesis response regulator RcsB | |
| fig\|634500 | RcsC | Erwinia billingiae Eb661 | 634500,5 | NC_014306 | EbC_3039 | VBIErwBil9 | PATRIC.63 | PATRIC | CDS | 3411886 | 3414735 | 2850 | - | FIG000042 | PLF_551_ | PGF_0005 | YP_00374 | 949 | rcsC | Sensor hist | GO:0004673\|protein histidine kinase activity |
| fig\|634500 | IgaA | Erwinia billingiae Eb661 | 634500,5 | NC_014306 | EbC_4198 | VBIErwBil9 | PATRIC.63 | PATRIC | CDS | 4652613 | 4654751 | 2139 | + | FIG000040 | PLF_551_ | PGF_0001 | YP_00374 | 712 | IgaA | IgaA: a membrane protein that prevents overactivation of the Rcs regulatory system | |
| fig\|634499 | DnaK | Erwinia pyrifoliae Ep1/96 | 634499.3 | NC_012214 | EpC_0695 | VBIErwPyr | PATRIC.63 | PATRIC | CDS | 805934 | 807847 | 1914 | + | FIG000233 | PLF_551_ | PGF_1035 | YP_00264 | 637 | dnaK | Chaperone protein DnaK | |
| fig\|634499 | RcsC | Erwinia pyrifoliae Ep1/96 | 634499,3 | NC_012214 | EpC_1287 | VBIErwPyr | PATRIC.63 | PATRIC | CDS | 1488266 | 1491244 | 2979 | + | FIG000042 | PLF_551_ | PGF_0005 | YP_00264 | 992 | rcsC | Sensor hist | GO:0004673\|protein histidine kinase activity |
| fig\|634499 | RcsB | Erwinia pyrifoliae Ep1/96 | 634499,3 | NC_012214 | EpC_1288 | VBIErwPyr | PATRIC.63 | PATRIC | CDS | 1491397 | 1492044 | 648 | - | FIG000041 | PLF_551_ | PGF_0042 | YP_00264 | 215 | rcsB | DNA-binding capsular synthesis response regulator RcsB | |
| fig\|634499 | RcsD | Erwinia pyrifoliae Ep1/96 | 634499,3 | NC_012214 | EpC_1289 | VBIErwPyr | PATRIC.63 | PATRIC | CDS | 1492046 | 1494703 | 2658 | - | FIG000045 | PLF_551_ | PGF_0003 | YP_00264 | 885 | yojN | Phosphotransferase RcsD | |
| fig\|634499 | IgaA | Erwinia pyrifoliae Ep1/96 | 634499,3 | NC_012214 | EpC_3423 | VBIErwPyr | PATRIC.63 | PATRIC | CDS | 3727998 | 3730136 | 2139 | + | FIG000040 | PLF_551_ | PGF_0001 | YP_00265 | 712 | | IgaA: a membrane protein that prevents overactivation of the Rcs regulatory system | |
| fig\|634499 | RcsF | Erwinia pyrifoliae Ep1/96 | 634499,3 | NC_012214 | EpC_0896 | VBIErwPyr | PATRIC.63 | PATRIC | CDS | 1034850 | 1035050 | 201 | - | FIG013040 | PLF_551_ | PGF_0003 | YP_00264 | 66 | rcsF | Protein RcsF | |
| fig\|465817 | DnaK | Erwinia tasmaniensis Et1/99 | 465817.9 | NC_010694 | ETA_0705 | VBIErwTas | PATRIC.46 | PATRIC | CDS | 785209 | 787122 | 1914 | + | FIG000233 | PLF_551_ | PGF_1035 | YP_00190 | 637 | dnaK | Chaperone protein DnaK | |
| fig\|465817 | RcsC | Erwinia tasmaniensis Et1/99 | 465817,9 | NC_010694 | ETA_1229 | VBIErwTas | PATRIC.46 | PATRIC | CDS | 1377580 | 1380435 | 2856 | + | FIG000042 | PLF_551_ | PGF_0005 | YP_00190 | 951 | rcsC | Sensor hist | GO:0004673\|protein histidine kinase activity |
| fig\|465817 | RcsB | Erwinia tasmaniensis Et1/99 | 465817,9 | NC_010694 | ETA_1230 | VBIErwTas | PATRIC.46 | PATRIC | CDS | 1380576 | 1381223 | 648 | - | FIG000041 | PLF_551_ | PGF_0042 | YP_00190 | 215 | rcsB | DNA-binding capsular synthesis response regulator RcsB | |
| fig\|465817 | RcsD | Erwinia tasmaniensis Et1/99 | 465817,9 | NC_010694 | ETA_1231 | VBIErwTas | PATRIC.46 | PATRIC | CDS | 1381225 | 1383882 | 2658 | - | FIG000045 | PLF_551_ | PGF_0003 | YP_00190 | 885 | yojN | Phosphotransferase RcsD | |
| fig\|465817 | IgaA | Erwinia tasmaniensis Et1/99 | 465817,9 | NC_010694 | ETA_3214 | VBIErwTas | PATRIC.46 | PATRIC | CDS | 3577766 | 3579829 | 2064 | + | FIG000040 | PLF_551_ | PGF_0001 | YP_00190 | 687 | | IgaA: a membrane protein that prevents overactivation of the Rcs regulatory system | |
| fig\|465817 | RcsF | Erwinia tasmaniensis Et1/99 | 465817,9 | NC_010694 | ETA_0917 | VBIErwTas | PATRIC.46 | PATRIC | CDS | 1024741 | 1025274 | 534 | - | FIG013040 | PLF_551_ | PGF_0003 | YP_00190 | 177 | rcsF | Protein RcsF | |
| fig\|502347 | DnaK | Escherichia albertii TW07627 | 502347.3 | NZ_CH991859 | ESCAB762 | VBIEscAlb | PATRIC.50 | PATRIC | CDS | 819621 | 821537 | 1917 | - | FIG000233 | PLF_561_ | PGF_1035 | ZP_02903 | 638 | dnaK | Chaperone protein DnaK | |
| fig\|502347 | RcsC | Escherichia albertii TW07627 | 502347,3 | NZ_CH991859 | ESCAB762 | VBIEscAlb | PATRIC.50 | PATRIC | CDS | 3249730 | 3252579 | 2850 | + | FIG000042 | PLF_561_ | PGF_0005 | ZP_02902 | 949 | rcsC | Sensor hist | GO:0004673\|protein histidine kinase activity |
| fig\|502347 | RcsB | Escherichia albertii TW07627 | 502347,3 | NZ_CH991859 | ESCAB762 | VBIEscAlb | PATRIC.50 | PATRIC | CDS | 3253121 | 3253771 | 651 | - | FIG000041 | PLF_561_ | PGF_0042 | ZP_02901 | 216 | rcsB | DNA-binding capsular synthesis response regulator RcsB | |
| fig\|502347 | RcsD | Escherichia albertii TW07627 | 502347,3 | NZ_CH991859 | ESCAB762 | VBIEscAlb | PATRIC.50 | PATRIC | CDS | 3253788 | 3256460 | 2673 | - | FIG000045 | PLF_561_ | PGF_0003 | ZP_02901 | 890 | yojN | Phosphotransferase RcsD | |
| fig\|502347 | RcsF | Escherichia albertii TW07627 | 502347,3 | NZ_CH991859 | ESCAB762 | VBIEscAlb | PATRIC.50 | PATRIC | CDS | 617279 | 617683 | 405 | + | FIG013040 | PLF_561_ | PGF_0003 | ZP_02903 | 134 | rcsF | Protein RcsF | |
| fig\|502347 | IgaA | Escherichia albertii TW07627 | 502347,3 | NZ_CH991859 | ESCAB762 | VBIEscAlb | PATRIC.50 | PATRIC | CDS | 1158511 | 1160556 | 2046 | + | FIG000040 | PLF_561_ | PGF_0001 | ZP_02901 | 681 | | IgaA: a membrane protein that prevents overactivation of the Rcs regulatory system | |
| fig\|113385 | DnaK | Escherichia coli O104:H4 str. 2011C-3493 | 1133852.3 | CP003289 | O3K_2147 | VBIEscCol | PATRIC.11 | PATRIC | CDS | 4436455 | 4438371 | 1917 | - | FIG000233 | PLF_561_ | PGF_1035 | AFS76158 | 638 | dnaK | Chaperone protein DnaK | |
| fig\|113385 | RcsC | Escherichia coli O104:H4 str. 2011C-3493 | 1133852,3 | CP003289 | O3K_0835 | VBIEscCol | PATRIC.11 | PATRIC | CDS | 1737849 | 1740698 | 2850 | + | FIG000042 | PLF_561_ | PGF_0005 | AFS73575 | 949 | | Sensor hist | GO:0004673\|protein histidine kinase activity |
| fig\|113385 | RcsB | Escherichia coli O104:H4 str. 2011C-3493 | 1133852,3 | CP003289 | O3K_0836 | VBIEscCol | PATRIC.11 | PATRIC | CDS | 1740898 | 1741548 | 651 | - | FIG000041 | PLF_561_ | PGF_0042 | AFS73576 | 216 | | DNA-binding capsular synthesis response regulator RcsB | |
| fig\|113385 | RcsD | Escherichia coli O104:H4 str. 2011C-3493 | 1133852,3 | CP003289 | O3K_0836 | VBIEscCol | PATRIC.11 | PATRIC | CDS | 1741565 | 1744237 | 2673 | - | FIG000045 | PLF_561_ | PGF_0003 | AFS73577 | 890 | | Phosphotransferase RcsD | |
| fig\|113385 | RcsF | Escherichia coli O104:H4 str. 2011C-3493 | 1133852,3 | CP003289 | O3K_2057 | VBIEscCol | PATRIC.11 | PATRIC | CDS | 4230105 | 4230458 | 354 | + | FIG013040 | PLF_561_ | PGF_0003 | AFS75978 | 117 | rcsF | Protein RcsF | |
| fig\|113385 | IgaA | Escherichia coli O104:H4 str. 2011C-3493 | 1133852,3 | CP003289 | O3K_0208 | VBIEscCol | PATRIC.11 | PATRIC | CDS | 440064 | 442199 | 2136 | - | FIG000040 | PLF_561_ | PGF_0001 | AFS72345 | 711 | | IgaA: a membrane protein that prevents overactivation of the Rcs regulatory system | |
| fig\|386585 | DnaK | Escherichia coli O157:H7 str. Sakai | 386585.9 | NC_002695 | ECs0014 | VBIEscCol | PATRIC.38 | PATRIC | CDS | 12180 | 14096 | 1917 | + | FIG000233 | PLF_561_ | PGF_1035 | NP_30804 | 638 | dnaK | Chaperone protein DnaK | |
| fig\|386585 | RcsF | Escherichia coli O157:H7 str. Sakai | 386585,9 | NC_002695 | ECs0198 | VBIEscCol | PATRIC.38 | PATRIC | CDS | 222921 | 223274 | 354 | - | FIG013040 | PLF_561_ | PGF_0003 | NP_30822 | 117 | rcsF | Protein RcsF | |
| fig\|386585 | RcsD | Escherichia coli O157:H7 str. Sakai | 386585,9 | NC_002695 | ECs3105 | VBIEscCol | PATRIC.38 | PATRIC | CDS | 3046475 | 3049147 | 2673 | + | FIG000045 | PLF_561_ | PGF_0003 | NP_31113 | 890 | | Phosphotransferase RcsD | |
| fig\|386585 | RcsB | Escherichia coli O157:H7 str. Sakai | 386585,9 | NC_002695 | ECs3106 | VBIEscCol | PATRIC.38 | PATRIC | CDS | 3049164 | 3049814 | 651 | + | FIG000041 | PLF_561_ | PGF_0042 | NP_31113 | 216 | | DNA-binding capsular synthesis response regulator RcsB | |
| fig\|386585 | RcsC | Escherichia coli O157:H7 str. Sakai | 386585,9 | NC_002695 | ECs3107 | VBIEscCol | PATRIC.38 | PATRIC | CDS | 3050014 | 3052863 | 2850 | - | FIG000042 | PLF_561_ | PGF_0005 | NP_31113 | 949 | | Sensor hist | GO:0004673\|protein histidine kinase activity |
| fig\|386585 | IgaA | Escherichia coli O157:H7 str. Sakai | 386585,9 | NC_002695 | ECs4240 | VBIEscCol | PATRIC.38 | PATRIC | CDS | 4238663 | 4240798 | 2136 | + | FIG000040 | PLF_561_ | PGF_0001 | NP_31226 | 711 | | IgaA: a membrane protein that prevents overactivation of the Rcs regulatory system | |
| fig\|685038 | DnaK | Escherichia coli O83:H1 str. NRG 857C | 685038.3 | CP001855 | NRG857_0 | VBIEscCol | PATRIC.68 | PATRIC | CDS | 12231 | 14147 | 1917 | + | FIG000233 | PLF_561_ | PGF_1035 | ADR25446 | 638 | dnaK | Chaperone protein DnaK | |
| fig\|685038 | RcsF | Escherichia coli O83:H1 str. NRG 857C | 685038,3 | CP001855 | NRG857_0 | VBIEscCol | PATRIC.68 | PATRIC | CDS | 223658 | 224011 | 354 | - | FIG013040 | PLF_561_ | PGF_0003 | ADR25632 | 117 | rcsF | Protein RcsF | |
| fig\|685038 | RcsD | Escherichia coli O83:H1 str. NRG 857C | 685038,3 | CP001855 | NRG857_1 | VBIEscCol | PATRIC.68 | PATRIC | CDS | 2335689 | 2338361 | 2673 | + | FIG000045 | PLF_561_ | PGF_0003 | ADR27665 | 890 | | Phosphotransferase RcsD | |
| fig\|685038 | RcsB | Escherichia coli O83:H1 str. NRG 857C | 685038,3 | CP001855 | NRG857_1 | VBIEscCol | PATRIC.68 | PATRIC | CDS | 2338378 | 2339028 | 651 | + | FIG000041 | PLF_561_ | PGF_0042 | ADR27666 | 216 | | DNA-binding capsular synthesis response regulator RcsB | |
| fig\|685038 | RcsC | Escherichia coli O83:H1 str. NRG 857C | 685038,3 | CP001855 | NRG857_1 | VBIEscCol | PATRIC.68 | PATRIC | CDS | 2339228 | 2342077 | 2850 | - | FIG000042 | PLF_561_ | PGF_0005 | ADR27667 | 949 | | Sensor hist | GO:0004673\|protein histidine kinase activity |
| fig\|685038 | IgaA | Escherichia coli O83:H1 str. NRG 857C | 685038,3 | CP001855 | NRG857_1 | VBIEscCol | PATRIC.68 | PATRIC | CDS | 3538492 | 3540627 | 2136 | + | FIG000040 | PLF_561_ | PGF_0001 | ADR28773 | 711 | | IgaA: a membrane protein that prevents overactivation of the Rcs regulatory system | |
| fig\|511145 | DnaK | Escherichia coli str. K-12 substr. MG1655 | 511145.12 | NC_000913 | b0014 | VBIEscCol | PATRIC.51 | PATRIC | CDS | 12163 | 14079 | 1917 | + | FIG000233 | PLF_561_ | PGF_1035 | NP_41455 | 638 | dnaK | Chaperone protein DnaK | |
| fig\|511145 | RcsF | Escherichia coli str. K-12 substr. MG1655 | 511145,12 | NC_000913 | b0196 | VBIEscCol | PATRIC.51 | PATRIC | CDS | 219591 | 219944 | 354 | - | FIG013040 | PLF_561_ | PGF_0003 | NP_41473 | 117 | rcsF | Protein RcsF | |
| fig\|511145 | RcsD | Escherichia coli str. K-12 substr. MG1655 | 511145,12 | NC_000913 | b2216 | VBIEscCol | PATRIC.51 | PATRIC | CDS | 2311510 | 2314182 | 2673 | + | FIG000045 | PLF_561_ | PGF_0003 | NP_41672 | 890 | rcsD | Phosphotransferase RcsD | |
| fig\|511145 | RcsB | Escherichia coli str. K-12 substr. MG1655 | 511145,12 | NC_000913 | b2217 | VBIEscCol | PATRIC.51 | PATRIC | CDS | 2314199 | 2314849 | 651 | + | FIG000041 | PLF_561_ | PGF_0042 | NP_41672 | 216 | rcsB | DNA-binding capsular synthesis response regulator RcsB | |
| fig\|511145 | RcsC | Escherichia coli str. K-12 substr. MG1655 | 511145,12 | NC_000913 | b2218 | VBIEscCol | PATRIC.51 | PATRIC | CDS | 2315049 | 2317898 | 2850 | - | FIG000042 | PLF_561_ | PGF_0005 | NP_41672 | 949 | rcsC | Sensor hist | GO:0004673\|protein histidine kinase activity |
| fig\|511145 | IgaA | Escherichia coli str. K-12 substr. MG1655 | 511145,12 | NC_000913 | b3398 | VBIEscCol | PATRIC.51 | PATRIC | CDS | 3524491 | 3526626 | 2136 | + | FIG000040 | PLF_561_ | PGF_0001 | NP_41785 | 711 | yrfF | IgaA: a membrane protein that prevents overactivation of the Rcs regulatory system | |
| fig\|585056 | DnaK | Escherichia coli UMN026 | 585056.7 | NC_011751 | ECUMN_0 | VBIEscCol | PATRIC.58 | PATRIC | CDS | 12120 | 14036 | 1917 | + | FIG000233 | PLF_561_ | PGF_1035 | YP_00241 | 638 | dnaK | Chaperone protein DnaK | |
| fig\|585056 | RcsD | Escherichia coli UMN026 | 585056,7 | NC_011751 | ECUMN_2 | VBIEscCol | PATRIC.58 | PATRIC | CDS | 2613960 | 2616632 | 2673 | + | FIG000045 | PLF_561_ | PGF_0003 | YP_00241 | 890 | rcsD | Phosphotransferase RcsD | |
| fig\|585056 | RcsB | Escherichia coli UMN026 | 585056,7 | NC_011751 | ECUMN_2 | VBIEscCol | PATRIC.58 | PATRIC | CDS | 2616649 | 2617299 | 651 | + | FIG000041 | PLF_561_ | PGF_0042 | YP_00241 | 216 | rcsB | DNA-binding capsular synthesis response regulator RcsB | |
| fig\|585056 | RcsC | Escherichia coli UMN026 | 585056,7 | NC_011751 | ECUMN_2 | VBIEscCol | PATRIC.58 | PATRIC | CDS | 2617499 | 2620348 | 2850 | - | FIG000042 | PLF_561_ | PGF_0005 | YP_00241 | 949 | rcsC | Sensor hist | GO:0004673\|protein histidine kinase activity |
| fig\|585056 | RcsF | Escherichia coli UMN026 | 585056,7 | NC_011751 | ECUMN_0 | VBIEscCol | PATRIC.58 | PATRIC | CDS | 222441 | 222845 | 405 | - | FIG013040 | PLF_561_ | PGF_0003 | YP_00241 | 134 | rcsF | Protein RcsF | |
| fig\|585056 | IgaA | Escherichia coli UMN026 | 585056,7 | NC_011751 | ECUMN_3 | VBIEscCol | PATRIC.58 | PATRIC | CDS | 3952286 | 3954421 | 2136 | + | FIG000040 | PLF_561_ | PGF_0001 | YP_00241 | 711 | igaA | IgaA: a membrane protein that prevents overactivation of the Rcs regulatory system | |
| fig\|585054 | DnaK | Escherichia fergusonii ATCC 35469 | 585054.5 | NC_011740 | EFER_001 | VBIEscFer | PATRIC.58 | PATRIC | CDS | 10750 | 12666 | 1917 | + | FIG000233 | PLF_561_ | PGF_1035 | YP_00238 | 638 | dnaK | Chaperone protein DnaK | |
| fig\|585054 | RcsF | Escherichia fergusonii ATCC 35469 | 585054,5 | NC_011740 | EFER_022 | VBIEscFer | PATRIC.58 | PATRIC | CDS | 241009 | 241362 | 354 | - | FIG013040 | PLF_561_ | PGF_0003 | YP_00238 | 117 | rcsF | Protein RcsF | |
| fig\|585054 | IgaA | Escherichia fergusonii ATCC 35469 | 585054,5 | NC_011740 | EFER_336 | VBIEscFer | PATRIC.58 | PATRIC | CDS | 3420865 | 3423000 | 2136 | + | FIG000040 | PLF_561_ | PGF_0001 | YP_00238 | 711 | igaA | IgaA: a membrane protein that prevents overactivation of the Rcs regulatory system | |
| fig\|585054 | RcsC | Escherichia fergusonii ATCC 35469 | 585054,5 | NC_011740 | EFER_094 | VBIEscFer | PATRIC.58 | PATRIC | CDS | 965524 | 968373 | 2850 | + | FIG000042 | PLF_561_ | PGF_0005 | YP_00238 | 949 | rcsC | Sensor hist | GO:0004673\|protein histidine kinase activity |
| fig\|585054 | RcsB | Escherichia fergusonii ATCC 35469 | 585054,5 | NC_011740 | EFER_094 | VBIEscFer | PATRIC.58 | PATRIC | CDS | 968457 | 969107 | 651 | - | FIG000041 | PLF_561_ | PGF_0042 | YP_00238 | 216 | rcsB | DNA-binding capsular synthesis response regulator RcsB | |
| fig\|585054 | RcsD | Escherichia fergusonii ATCC 35469 | 585054,5 | NC_011740 | EFER_094 | VBIEscFer | PATRIC.58 | PATRIC | CDS | 969124 | 971793 | 2670 | - | FIG000045 | PLF_561_ | PGF_0003 | YP_00238 | 889 | rcsD | Phosphotransferase RcsD | |
| fig\|111551 | DnaK | Escherichia hermannii NBRC 105704 | 1115512.3 | BAFF01000002 | EH105704 | VBIEscHer | PATRIC.11 | PATRIC | CDS | 290358 | 292274 | 1917 | + | FIG00023369 | | PGF_1035 | GAB51236 | 638 | dnaK | Chaperone protein DnaK | |
| fig\|111551 | IgaA | Escherichia hermannii NBRC 105704 | 1115512,3 | BAFF01000007 | EH105704 | VBIEscHer | PATRIC.11 | PATRIC | CDS | 51775 | 53913 | 2139 | + | FIG00004077 | | PGF_0001 | GAB52488 | 712 | yrfF | IgaA: a membrane protein that prevents overactivation of the Rcs regulatory system | |
| fig\|111551 | RcsF | Escherichia hermannii NBRC 105704 | 1115512,3 | BAFF01000009 | EH105704 | VBIEscHer | PATRIC.11 | PATRIC | CDS | 125016 | 125420 | 405 | - | FIG01304060 | | PGF_0003 | GAB52889 | 134 | rcsF | Protein RcsF | |
| fig\|111551 | RcsB | Escherichia hermannii NBRC 105704 | 1115512,3 | BAFF01000001 | EH105704 | VBIEscHer | PATRIC.11 | PATRIC | CDS | 676101 | 676751 | 651 | - | FIG00004153 | | PGF_0042 | GAB50655 | 216 | rcsB | DNA-binding capsular synthesis response regulator RcsB | |
| fig\|111551 | RcsD | Escherichia hermannii NBRC 105704 | 1115512,3 | BAFF01000001 | EH105704 | VBIEscHer | PATRIC.11 | PATRIC | CDS | 676768 | 679332 | 2565 | - | FIG00004592 | | PGF_0003 | GAB50656 | 854 | rcsD | Phosphotransferase RcsD | |
| fig\|111551 | RcsC | Escherichia hermannii NBRC 105704 | 1115512,3 | BAFF01000001 | EH105704 | VBIEscHer | PATRIC.11 | PATRIC | CDS | 687787 | 690636 | 2850 | - | FIG00004208 | | PGF_0005 | GAB50663 | 949 | rcsC | Sensor hist | GO:0004673\|protein histidine kinase activity |
| fig\|111551 | DnaK | Escherichia vulneris NBRC 102420 | 1115515.3 | BBMZ01000002 | EV102420_02_03460 | | PATRIC.11 | PATRIC | CDS | 369488 | 371404 | 1917 | - | FIG00023369 | | PGF_1035 | GAL56741 | 638 | | Chaperone protein DnaK | |
| fig\|111551 | RcsF | Escherichia vulneris NBRC 102420 | 1115515,3 | BBMZ01000008 | EV102420_08_04270 | | PATRIC.11 | PATRIC | CDS | 453254 | 453661 | 408 | - | FIG01304060 | | PGF_0003 | GAL57964 | 135 | | Protein RcsF | |
| fig\|111551 | IgaA | Escherichia vulneris NBRC 102420 | 1115515,3 | BBMZ01000016 | EV102420_16_00570 | | PATRIC.11 | PATRIC | CDS | 50925 | 53072 | 2148 | + | FIG00004077 | | PGF_0001 | GAL59338 | 715 | | IgaA: a membrane protein that prevents overactivation of the Rcs regulatory system | |
| fig\|111551 | RcsC | Escherichia vulneris NBRC 102420 | 1115515,3 | BBMZ01000007 | EV102420_07_01740 | | PATRIC.11 | PATRIC | CDS | 188703 | 191546 | 2844 | + | FIG00004208 | | PGF_0005 | GAL57355 | 947 | | Sensor hist | GO:0004673\|protein histidine kinase activity |

Table-S2

| | | | | | | | | | | | | | | | | | | | | |
|---|---|---|---|---|---|---|---|---|---|---|---|---|---|---|---|---|---|---|---|---|
| fig\|111551 | RcsB | Escherichia vulneris NBRC 102420 | 1115515,3 | BBMZ01000007 | EV102420_07_01750 | | PATRIC.11 | PATRIC | CDS | 191606 | 192256 | 651 | - | FIG00004153 | | PGF_0042 | GAL57356 | 216 | | DNA-binding capsular synthesis response regulator RcsB |
| fig\|111551 | RcsD | Escherichia vulneris NBRC 102420 | 1115515,3 | BBMZ01000007 | EV102420_07_01760 | | PATRIC.11 | PATRIC | CDS | 192273 | 194921 | 2649 | - | FIG00004592 | | PGF_0003 | GAL57357 | 882 | | Phosphotransferase RcsD |
| fig\|145349 | DnaK | Hafnia alvei FB1 | 1453496.5 | CP009706 | AT03_18495 | | PATRIC.14 | PATRIC | CDS | 4024526 | 4026436 | 1911 | - | FIG000233 | PLF_568_ | PGF_10357457 | | 636 | dnaK | Chaperone protein DnaK |
| fig\|145349 | RcsD | Hafnia alvei FB1 | 1453496,5 | CP009706 | AT03_14320 | | PATRIC.14 | PATRIC | CDS | 3083389 | 3086094 | 2706 | + | FIG000045 | PLF_568_ | PGF_00034034 | | 901 | | Phosphotransferase RcsD |
| fig\|145349 | RcsB | Hafnia alvei FB1 | 1453496,5 | CP009706 | AT03_14325 | | PATRIC.14 | PATRIC | CDS | 3086087 | 3086737 | 651 | + | FIG000041 | PLF_568_ | PGF_00421954 | | 216 | | DNA-binding capsular synthesis response regulator RcsB |
| fig\|145349 | RcsC | Hafnia alvei FB1 | 1453496,5 | CP009706 | AT03_14330 | | PATRIC.14 | PATRIC | CDS | 3086862 | 3089726 | 2865 | - | FIG000042 | PLF_568_ | PGF_00050577 | | 954 | | Sensor his GO:0004673\|protein histidine kinase activity |
| fig\|145349 | RcsF | Hafnia alvei FB1 | 1453496,5 | CP009706 | AT03_18680 | | PATRIC.14 | PATRIC | CDS | 4070294 | 4070713 | 420 | + | FIG013040 | PLF_568_ | PGF_00037945 | | 139 | rcsF | Protein RcsF |
| fig\|145349 | IgaA | Hafnia alvei FB1 | 1453496,5 | CP009706 | AT03_20610 | | PATRIC.14 | PATRIC | CDS | 4502863 | 4504977 | 2115 | + | FIG000040 | PLF_568_ | PGF_00013790 | | 704 | | IgaA: a membrane protein that prevents overactivation of the Rcs regulatory system |
| fig\|100655 | DnaK | Klebsiella oxytoca KCTC 1686 | 1006551.4 | CP003218 | KOX_1046 | VBIKleOxy | PATRIC.10 | PATRIC | CDS | 2231183 | 2233099 | 1917 | + | FIG000233 | PLF_570_ | PGF_1035 | AEX03819 | 638 | dnaK | Chaperone protein DnaK |
| fig\|100655 | RcsF | Klebsiella oxytoca KCTC 1686 | 1006551,4 | CP003218 | KOX_1159 | VBIKleOxy | PATRIC.10 | PATRIC | CDS | 2496787 | 2496918 | 132 | - | FIG013040 | PLF_570_ | PGF_0003 | AEX04044 | 43 | rcsF | Protein RcsF |
| fig\|100655 | RcsD | Klebsiella oxytoca KCTC 1686 | 1006551,4 | CP003218 | KOX_2611 | VBIKleOxy | PATRIC.10 | PATRIC | CDS | 5636876 | 5639536 | 2661 | + | FIG000045 | PLF_570_ | PGF_0003 | AEX06931 | 886 | | Phosphotransferase RcsD |
| fig\|100655 | RcsB | Klebsiella oxytoca KCTC 1686 | 1006551,4 | CP003218 | KOX_2611 | VBIKleOxy | PATRIC.10 | PATRIC | CDS | 5639553 | 5640203 | 651 | + | FIG000041 | PLF_570_ | PGF_0042 | AEX06932 | 216 | | DNA-binding capsular synthesis response regulator RcsB |
| fig\|100655 | RcsC | Klebsiella oxytoca KCTC 1686 | 1006551,4 | CP003218 | KOX_2612 | VBIKleOxy | PATRIC.10 | PATRIC | CDS | 5640248 | 5643094 | 2847 | - | FIG000042 | PLF_570_ | PGF_0005 | AEX06933 | 948 | | Sensor his GO:0004673\|protein histidine kinase activity |
| fig\|100655 | IgaA | Klebsiella oxytoca KCTC 1686 | 1006551,4 | CP003218 | KOX_0465 | VBIKleOxy | PATRIC.10 | PATRIC | CDS | 950091 | 952226 | 2136 | + | FIG000040 | PLF_570_ | PGF_0001 | AEX02667 | 711 | | IgaA: a membrane protein that prevents overactivation of the Rcs regulatory system |
| fig\|272620 | DnaK | Klebsiella pneumoniae subsp. pneumoniae MGH 78578 | 272620.9 | NC_009648 | KPN_0001 | VBIKlePne | PATRIC.27 | PATRIC | CDS | 13612 | 15528 | 1917 | + | FIG000233 | PLF_570_ | PGF_1035 | YP_00133 | 638 | dnaK | Chaperone protein DnaK |
| fig\|272620 | RcsF | Klebsiella pneumoniae subsp. pneumoniae MGH 78578 | 272620,9 | NC_009648 | KPN_0021 | VBIKlePne | PATRIC.27 | PATRIC | CDS | 245325 | 245681 | 357 | - | FIG013040 | PLF_570_ | PGF_0003 | YP_00133 | 118 | rcsF | Protein RcsF |
| fig\|272620 | RcsD | Klebsiella pneumoniae subsp. pneumoniae MGH 78578 | 272620,9 | NC_009648 | KPN_0263 | VBIKlePne | PATRIC.27 | PATRIC | CDS | 2896683 | 2899340 | 2658 | + | FIG000045 | PLF_570_ | PGF_0003 | YP_00133 | 885 | yojN | Phosphotransferase RcsD |
| fig\|272620 | RcsB | Klebsiella pneumoniae subsp. pneumoniae MGH 78578 | 272620,9 | NC_009648 | KPN_0263 | VBIKlePne | PATRIC.27 | PATRIC | CDS | 2899356 | 2900006 | 651 | + | FIG000041 | PLF_570_ | PGF_0042 | YP_00133 | 216 | rcsB | DNA-binding capsular synthesis response regulator RcsB |
| fig\|272620 | RcsC | Klebsiella pneumoniae subsp. pneumoniae MGH 78578 | 272620,9 | NC_009648 | KPN_0263 | VBIKlePne | PATRIC.27 | PATRIC | CDS | 2900052 | 2902892 | 2841 | - | FIG000042 | PLF_570_ | PGF_0005 | YP_00133 | 946 | rcsC | Sensor his GO:0004673\|protein histidine kinase activity |
| fig\|272620 | IgaA | Klebsiella pneumoniae subsp. pneumoniae MGH 78578 | 272620,9 | NC_009648 | KPN_0376 | VBIKlePne | PATRIC.27 | PATRIC | CDS | 4113123 | 4115252 | 2130 | + | FIG000040 | PLF_570_ | PGF_0001 | YP_00133 | 709 | yrfF | IgaA: a membrane protein that prevents overactivation of the Rcs regulatory system |
| fig\|640131 | DnaK | Klebsiella variicola At-22 | 640131.3 | NC_013850 | Kvar_4380 | VBIKleVar | PATRIC.64 | PATRIC | CDS | 4639093 | 4641009 | 1917 | - | FIG000233 | PLF_570_ | PGF_1035 | YP_00344 | 638 | | Chaperone protein DnaK |
| fig\|640131 | RcsC | Klebsiella variicola At-22 | 640131,3 | NC_013850 | Kvar_1414 | VBIKleVar | PATRIC.64 | PATRIC | CDS | 1498611 | 1501451 | 2841 | + | FIG000042 | PLF_570_ | PGF_0005 | YP_00343 | 946 | | Sensor his GO:0004673\|protein histidine kinase activity |
| fig\|640131 | RcsB | Klebsiella variicola At-22 | 640131,3 | NC_013850 | Kvar_1415 | VBIKleVar | PATRIC.64 | PATRIC | CDS | 1501497 | 1502147 | 651 | - | FIG000041 | PLF_570_ | PGF_0042 | YP_00343 | 216 | | DNA-binding capsular synthesis response regulator RcsB |
| fig\|640131 | RcsD | Klebsiella variicola At-22 | 640131,3 | NC_013850 | Kvar_1416 | VBIKleVar | PATRIC.64 | PATRIC | CDS | 1502163 | 1504820 | 2658 | - | FIG000045 | PLF_570_ | PGF_0003 | YP_00343 | 885 | | Phosphotransferase RcsD |
| fig\|640131 | IgaA | Klebsiella variicola At-22 | 640131,3 | NC_013850 | Kvar_0332 | VBIKleVar | PATRIC.64 | PATRIC | CDS | 370828 | 372963 | 2136 | - | FIG000040 | PLF_570_ | PGF_0001 | YP_00343 | 711 | | IgaA: a membrane protein that prevents overactivation of the Rcs regulatory system |
| fig\|640131 | RcsF | Klebsiella variicola At-22 | 640131,3 | NC_013850 | Kvar_4171 | VBIKleVar | PATRIC.64 | PATRIC | CDS | 4395842 | 4396198 | 357 | + | FIG013040 | PLF_570_ | PGF_0003 | YP_00344 | 118 | | Protein RcsF |
| fig\|100600 | DnaK | Kluyvera ascorbata ATCC 33433 | 1006000.3 | JMPL01000091 | GKAS_03565 | | PATRIC.10 | PATRIC | CDS | 15199 | 17121 | 1923 | + | FIG000233 | PLF_579_ | PGF_1035 | KFC99038 | 640 | dnaK | Chaperone protein DnaK |
| fig\|100600 | RcsC | Kluyvera ascorbata ATCC 33433 | 1006000,3 | JMPL01000014 | GKAS_01200 | | PATRIC.10 | PATRIC | CDS | 20367 | 23210 | 2844 | + | FIG000042 | PLF_579_ | PGF_0005 | KFD06865 | 947 | rcsC | Sensor his GO:0004673\|protein histidine kinase activity |
| fig\|100600 | RcsB | Kluyvera ascorbata ATCC 33433 | 1006000,3 | JMPL01000014 | GKAS_01201 | | PATRIC.10 | PATRIC | CDS | 23283 | 23933 | 651 | - | FIG000041 | PLF_579_ | PGF_0042 | KFD06866 | 216 | rcsB | DNA-binding capsular synthesis response regulator RcsB |
| fig\|100600 | RcsD | Kluyvera ascorbata ATCC 33433 | 1006000,3 | JMPL01000014 | GKAS_01202 | | PATRIC.10 | PATRIC | CDS | 23949 | 26549 | 2601 | - | FIG000045 | PLF_579_ | PGF_0003 | KFD06867 | 866 | rcsD | Phosphotransferase RcsD |
| fig\|100600 | RcsF | Kluyvera ascorbata ATCC 33433 | 1006000,3 | JMPL01000006 | GKAS_00678 | | PATRIC.10 | PATRIC | CDS | 86301 | 86705 | 405 | - | FIG013040 | PLF_579_ | PGF_0003 | KFD08094 | 134 | rcsF | Protein RcsF |
| fig\|100600 | IgaA | Kluyvera ascorbata ATCC 33433 | 1006000,3 | JMPL01000008 | GKAS_00783 | | PATRIC.10 | PATRIC | CDS | 13763 | 15910 | 2148 | + | FIG000040 | PLF_579_ | PGF_0001 | KFD07630 | 715 | yrfF | IgaA: a membrane protein that prevents overactivation of the Rcs regulatory system |
| fig\|121811 | DnaK | Kluyvera cryocrescens NBRC 102467 | 1218112.3 | BCTM01000015 | | | PATRIC.12 | PATRIC | CDS | 23856 | 25778 | 1923 | - | FIG000233 | PLF_579_ | PGF_10357457 | | 640 | | Chaperone protein DnaK |
| fig\|121811 | RcsF | Kluyvera cryocrescens NBRC 102467 | 1218112,3 | BCTM01000011 | | | PATRIC.12 | PATRIC | CDS | 187477 | 187830 | 354 | - | FIG013040 | PLF_579_ | PGF_00037945 | | 117 | | Protein RcsF |
| fig\|121811 | IgaA | Kluyvera cryocrescens NBRC 102467 | 1218112,3 | BCTM01000012 | | | PATRIC.12 | PATRIC | CDS | 48439 | 50586 | 2148 | + | FIG000040 | PLF_579_ | PGF_00013790 | | 715 | | IgaA: a membrane protein that prevents overactivation of the Rcs regulatory system |
| fig\|121811 | RcsD | Kluyvera cryocrescens NBRC 102467 | 1218112,3 | BCTM01000023 | | | PATRIC.12 | PATRIC | CDS | 500 | 3175 | 2676 | + | FIG000045 | PLF_579_ | PGF_00034034 | | 891 | | Phosphotransferase RcsD |
| fig\|121811 | RcsB | Kluyvera cryocrescens NBRC 102467 | 1218112,3 | BCTM01000023 | | | PATRIC.12 | PATRIC | CDS | 3191 | 3841 | 651 | + | FIG000041 | PLF_579_ | PGF_00421954 | | 216 | | DNA-binding capsular synthesis response regulator RcsB |
| fig\|121811 | RcsC | Kluyvera cryocrescens NBRC 102467 | 1218112,3 | BCTM01000023 | | | PATRIC.12 | PATRIC | CDS | 3961 | 6807 | 2847 | - | FIG000042 | PLF_579_ | PGF_00050577 | | 948 | | Sensor his GO:0004673\|protein histidine kinase activity |
| fig\|61648.5 | DnaK | Kluyvera intermedia strain CAV1151 | 61648.5 | CP011602 | AB182_21655 | | PATRIC.61 | PATRIC | CDS | 3846253 | 3848169 | 1917 | - | FIG000233 | PLF_579_ | PGF_10357457 | | 638 | dnaK | Chaperone protein DnaK |
| fig\|61648.5 | RcsF | Kluyvera intermedia strain CAV1151 | 61648,5 | CP011602 | AB182_20785 | | PATRIC.61 | PATRIC | CDS | 3640870 | 3641274 | 405 | + | FIG013040 | PLF_579_ | PGF_00037945 | | 134 | rcsF | Protein RcsF |
| fig\|61648.5 | IgaA | Kluyvera intermedia strain CAV1151 | 61648,5 | CP011602 | AB182_27415 | | PATRIC.61 | PATRIC | CDS | 5073430 | 5075568 | 2139 | - | FIG000040 | PLF_579_ | PGF_00013790 | | 712 | | IgaA: a membrane protein that prevents overactivation of the Rcs regulatory system |
| fig\|61648.5 | RcsC | Kluyvera intermedia strain CAV1151 | 61648,5 | CP011602 | AB182_07135 | | PATRIC.61 | PATRIC | CDS | 794406 | 797252 | 2847 | + | FIG000042 | PLF_579_ | PGF_00050577 | | 948 | | Sensor his GO:0004673\|protein histidine kinase activity |
| fig\|61648.5 | RcsB | Kluyvera intermedia strain CAV1151 | 61648,5 | CP011602 | AB182_07140 | | PATRIC.61 | PATRIC | CDS | 797287 | 797937 | 651 | - | FIG000041 | PLF_579_ | PGF_00421954 | | 216 | | DNA-binding capsular synthesis response regulator RcsB |
| fig\|61648.5 | RcsD | Kluyvera intermedia strain CAV1151 | 61648,5 | CP011602 | AB182_07145 | | PATRIC.61 | PATRIC | CDS | 797954 | 800581 | 2628 | - | FIG000045 | PLF_579_ | PGF_00034034 | | 875 | | Phosphotransferase RcsD |
| fig\|123583 | DnaK | Kosakonia sacchari SP1 | 1235834.6 | CP007215 | | | PATRIC.12 | PATRIC | CDS | 2340637 | 2342550 | 1914 | - | FIG000233 | PLF_13305 | PGF_10357457 | | 637 | | Chaperone protein DnaK |
| fig\|123583 | RcsF | Kosakonia sacchari SP1 | 1235834,6 | CP007215 | | | PATRIC.12 | PATRIC | CDS | 2156009 | 2156362 | 354 | + | FIG013040 | PLF_13305 | PGF_00037945 | | 117 | | Protein RcsF |
| fig\|123583 | IgaA | Kosakonia sacchari SP1 | 1235834,6 | CP007215 | | | PATRIC.12 | PATRIC | CDS | 3495401 | 3497539 | 2139 | - | FIG000040 | PLF_13305 | PGF_00013790 | | 712 | | IgaA: a membrane protein that prevents overactivation of the Rcs regulatory system |
| fig\|123583 | RcsC | Kosakonia sacchari SP1 | 1235834,6 | CP007215 | | | PATRIC.12 | PATRIC | CDS | 4707155 | 4710001 | 2847 | + | FIG000042 | PLF_13305 | PGF_00050577 | | 948 | | Sensor his GO:0004673\|protein histidine kinase activity |
| fig\|123583 | RcsB | Kosakonia sacchari SP1 | 1235834,6 | CP007215 | | | PATRIC.12 | PATRIC | CDS | 4710034 | 4710684 | 651 | - | FIG000041 | PLF_13305 | PGF_00421954 | | 216 | | DNA-binding capsular synthesis response regulator RcsB |
| fig\|123583 | RcsD | Kosakonia sacchari SP1 | 1235834,6 | CP007215 | | | PATRIC.12 | PATRIC | CDS | 4710701 | 4713352 | 2652 | - | FIG000045 | PLF_13305 | PGF_00034034 | | 883 | | Phosphotransferase RcsD |
| fig\|83655.1 | DnaK | Leclercia adecarboxylata strain USDA-ARS-USMARC-602 | 83655.10 | CP013990 | APT61_19000 | | PATRIC.83 | PATRIC | CDS | 4101474 | 4103393 | 1920 | - | FIG000233 | PLF_83654 | PGF_10357457 | | 639 | dnaK | Chaperone protein DnaK |
| fig\|83655.1 | RcsC | Leclercia adecarboxylata strain USDA-ARS-USMARC-602 | 83655,1 | CP013990 | APT61_07005 | | PATRIC.83 | PATRIC | CDS | 1502907 | 1505753 | 2847 | + | FIG000042 | PLF_83654 | PGF_00050577 | | 948 | | Sensor his GO:0004673\|protein histidine kinase activity |
| fig\|83655.1 | RcsB | Leclercia adecarboxylata strain USDA-ARS-USMARC-602 | 83655,1 | CP013990 | APT61_07010 | | PATRIC.83 | PATRIC | CDS | 1506572 | 1507222 | 651 | - | FIG000041 | PLF_83654 | PGF_00421954 | | 216 | | DNA-binding capsular synthesis response regulator RcsB |
| fig\|83655.1 | RcsD | Leclercia adecarboxylata strain USDA-ARS-USMARC-602 | 83655,1 | CP013990 | APT61_07015 | | PATRIC.83 | PATRIC | CDS | 1507239 | 1509911 | 2673 | - | FIG000045 | PLF_83654 | PGF_00034034 | | 890 | | Phosphotransferase RcsD |
| fig\|83655.1 | IgaA | Leclercia adecarboxylata strain USDA-ARS-USMARC-602 | 83655,1 | CP013990 | APT61_01765 | | PATRIC.83 | PATRIC | CDS | 388434 | 390569 | 2136 | - | FIG000040 | PLF_83654 | PGF_00013790 | | 711 | | IgaA: a membrane protein that prevents overactivation of the Rcs regulatory system |
| fig\|83655.1 | RcsF | Leclercia adecarboxylata strain USDA-ARS-USMARC-602 | 83655,1 | CP013990 | APT61_18125 | | PATRIC.83 | PATRIC | CDS | 3896988 | 3897392 | 405 | + | FIG013040 | PLF_83654 | PGF_00037945 | | 134 | rcsF | Protein RcsF |
| fig\|112499 | DnaK | Morganella morganii subsp. morganii KT | 1124991.3 | ALJX01000008 | MU9_1996 | VBIMorMor | PATRIC.11 | PATRIC | CDS | 77470 | 79392 | 1923 | + | FIG000233 | PLF_581_ | PGF_1035 | EJO25360 | 640 | dnaK | Chaperone protein DnaK |
| fig\|112499 | RcsD | Morganella morganii subsp. morganii KT | 1124991,3 | ALJX01000007 | MU9_1705 | VBIMorMor | PATRIC.11 | PATRIC | CDS | 5214 | 7928 | 2715 | + | FIG000045 | PLF_581_ | PGF_0003 | EJO25485 | 904 | rcsD | Phosphotransferase RcsD |
| fig\|112499 | RcsB | Morganella morganii subsp. morganii KT | 1124991,3 | ALJX01000007 | MU9_1706 | VBIMorMor | PATRIC.11 | PATRIC | CDS | 7921 | 8571 | 651 | + | FIG000041 | PLF_581_ | PGF_0042 | EJO25486 | 216 | rcsB | DNA-binding capsular synthesis response regulator RcsB |
| fig\|112499 | RcsC | Morganella morganii subsp. morganii KT | 1124991,3 | ALJX01000007 | MU9_1707 | VBIMorMor | PATRIC.11 | PATRIC | CDS | 8641 | 11487 | 2847 | - | FIG000042 | PLF_581_ | PGF_0005 | EJO25487 | 948 | rcsC | Sensor his GO:0004673\|protein histidine kinase activity |
| fig\|112499 | RcsF | Morganella morganii subsp. morganii KT | 1124991,3 | ALJX01000008 | MU9_2111 | VBIMorMor | PATRIC.11 | PATRIC | CDS | 212100 | 212510 | 411 | - | | PLF_581_ | PGF_0238 | EJO25475 | 136 | rcsF | hypothetical protein |
| fig\|112499 | IgaA | Morganella morganii subsp. morganii KT | 1124991,3 | ALJX01000025 | MU9_3381 | VBIMorMor | PATRIC.11 | PATRIC | CDS | 28870 | 30993 | 2124 | - | FIG000040 | PLF_581_ | PGF_0001 | EJO24042 | 707 | umoB | IgaA: a membrane protein that prevents overactivation of the Rcs regulatory system |
| fig\|549.24. | DnaK | Pantoea agglomerans strain FDAARGOS_160 | 549.24 | CP014129 | AL522_07335 | | PATRIC.54 | PATRIC | CDS | 853317 | 855227 | 1911 | + | FIG000233 | PLF_53335 | PGF_10357457 | | 636 | dnaK | Chaperone protein DnaK |
| fig\|549.24. | RcsD | Pantoea agglomerans strain FDAARGOS_160 | 549,24 | CP014129 | AL522_17440 | | PATRIC.54 | PATRIC | CDS | 2991214 | 2993871 | 2658 | + | FIG000045 | PLF_53335 | PGF_00034034 | | 885 | | Phosphotransferase RcsD |
| fig\|549.24. | RcsB | Pantoea agglomerans strain FDAARGOS_160 | 549,24 | CP014129 | AL522_17445 | | PATRIC.54 | PATRIC | CDS | 2993873 | 2994523 | 651 | + | FIG000041 | PLF_53335 | PGF_00421954 | | 216 | | DNA-binding capsular synthesis response regulator RcsB |
| fig\|549.24. | RcsC | Pantoea agglomerans strain FDAARGOS_160 | 549,24 | CP014129 | AL522_17450 | | PATRIC.54 | PATRIC | CDS | 2994577 | 2997429 | 2853 | - | FIG000042 | PLF_53335 | PGF_00050577 | | 950 | | Sensor his GO:0004673\|protein histidine kinase activity |
| fig\|549.24. | IgaA | Pantoea agglomerans strain FDAARGOS_160 | 549,24 | CP014129 | AL522_21885 | | PATRIC.54 | PATRIC | CDS | 3911963 | 3914098 | 2136 | + | FIG000040 | PLF_53335 | PGF_00013790 | | 711 | | IgaA: a membrane protein that prevents overactivation of the Rcs regulatory system |
| fig\|549.24. | RcsF | Pantoea agglomerans strain FDAARGOS_160 | 549,24 | CP014129 | AL522_08090 | | PATRIC.54 | PATRIC | CDS | 1033618 | 1034016 | 399 | - | FIG013040 | PLF_53335 | PGF_00037945 | | 132 | | Protein RcsF |
| fig\|706191 | DnaK | Pantoea ananatis LMG 20103 | 706191.3 | NC_013956 | PANA_066 | VBIPanAn | PATRIC.70 | PATRIC | CDS | 768793 | 770703 | 1911 | + | FIG000233 | PLF_53335 | PGF_1035 | YP_00351 | 636 | dnaK | Chaperone protein DnaK |
| fig\|706191 | RcsD | Pantoea ananatis LMG 20103 | 706191,3 | NC_013956 | PANA_260 | VBIPanAn | PATRIC.70 | PATRIC | CDS | 2927654 | 2930314 | 2661 | + | FIG000045 | PLF_53335 | PGF_0003 | YP_00352 | 886 | yojN | Phosphotransferase RcsD |
| fig\|706191 | RcsB | Pantoea ananatis LMG 20103 | 706191,3 | NC_013956 | PANA_261 | VBIPanAn | PATRIC.70 | PATRIC | CDS | 2930316 | 2930966 | 651 | + | FIG000041 | PLF_53335 | PGF_0042 | YP_00352 | 216 | rcsB | DNA-binding capsular synthesis response regulator RcsB |
| fig\|706191 | RcsC | Pantoea ananatis LMG 20103 | 706191,3 | NC_013956 | PANA_261 | VBIPanAn | PATRIC.70 | PATRIC | CDS | 2931006 | 2933855 | 2850 | - | FIG000042 | PLF_53335 | PGF_0005 | YP_00352 | 949 | rcsC | Sensor his GO:0004673\|protein histidine kinase activity |
| fig\|706191 | IgaA | Pantoea ananatis LMG 20103 | 706191,3 | NC_013956 | PANA_367 | VBIPanAn | PATRIC.70 | PATRIC | CDS | 4055580 | 4057640 | 2061 | + | FIG000040 | PLF_53335 | PGF_0001 | YP_00352 | 686 | yrfF | IgaA: a membrane protein that prevents overactivation of the Rcs regulatory system |
| fig\|706191 | RcsF | Pantoea ananatis LMG 20103 | 706191,3 | NC_013956 | PANA_081 | VBIPanAn | PATRIC.70 | PATRIC | CDS | 955428 | 955559 | 132 | - | FIG013040 | PLF_53335 | PGF_0003 | YP_00351 | 43 | rcsF | Protein RcsF |
| fig\|107655 | DnaK | Pantoea rwandensis ND04 | 1076550.3 | CP009454 | | | PATRIC.10 | PATRIC | CDS | 4052067 | 4053923 | 1857 | - | FIG000233 | PLF_53335 | PGF_10357457 | | 618 | | Chaperone protein DnaK |
| fig\|107655 | RcsC | Pantoea rwandensis ND04 | 1076550,3 | CP009454 | | | PATRIC.10 | PATRIC | CDS | 1932667 | 1935519 | 2853 | + | FIG000042 | PLF_53335 | PGF_00050577 | | 950 | | Sensor his GO:0004673\|protein histidine kinase activity |
| fig\|107655 | RcsB | Pantoea rwandensis ND04 | 1076550,3 | CP009454 | | | PATRIC.10 | PATRIC | CDS | 1935561 | 1936211 | 651 | - | FIG000041 | PLF_53335 | PGF_00421954 | | 216 | | DNA-binding capsular synthesis response regulator RcsB |
| fig\|107655 | RcsD | Pantoea rwandensis ND04 | 1076550,3 | CP009454 | | | PATRIC.10 | PATRIC | CDS | 1936213 | 1938873 | 2661 | - | FIG000045 | PLF_53335 | PGF_00034034 | | 886 | | Phosphotransferase RcsD |
| fig\|107655 | RcsF | Pantoea rwandensis ND04 | 1076550,3 | CP009454 | | | PATRIC.10 | PATRIC | CDS | 3873991 | 3874392 | 402 | + | FIG013040 | PLF_53335 | PGF_00037945 | | 133 | | Protein RcsF |
| fig\|107655 | IgaA | Pantoea rwandensis ND04 | 1076550,3 | CP009454 | | | PATRIC.10 | PATRIC | CDS | 845127 | 847268 | 2142 | - | FIG000040 | PLF_53335 | PGF_00013790 | | 713 | | IgaA: a membrane protein that prevents overactivation of the Rcs regulatory system |
| fig\|218491 | DnaK | Pectobacterium atrosepticum SCRI1043 | 218491.5 | NC_004547 | ECA3882 | VBIPecAtr | PATRIC.21 | PATRIC | CDS | 4343607 | 4345514 | 1908 | - | FIG000233 | PLF_12227 | PGF_1035 | YP_05197 | 635 | dnaK | Chaperone protein DnaK |
| fig\|218491 | RcsC | Pectobacterium atrosepticum SCRI1043 | 218491,5 | NC_004547 | ECA1202 | VBIPecAtr | PATRIC.21 | PATRIC | CDS | 1366323 | 1369109 | 2787 | + | FIG000042 | PLF_12227 | PGF_0005 | YP_04930 | 928 | rcsC | Sensor his GO:0004673\|protein histidine kinase activity |
| fig\|218491 | RcsB | Pectobacterium atrosepticum SCRI1043 | 218491,5 | NC_004547 | ECA1203 | VBIPecAtr | PATRIC.21 | PATRIC | CDS | 1369147 | 1369791 | 645 | - | FIG000041 | PLF_12227 | PGF_0042 | YP_04930 | 214 | rcsB | DNA-binding capsular synthesis response regulator RcsB |
| fig\|218491 | RcsD | Pectobacterium atrosepticum SCRI1043 | 218491,5 | NC_004547 | ECA1204 | VBIPecAtr | PATRIC.21 | PATRIC | CDS | 1369800 | 1372481 | 2682 | - | FIG000045 | PLF_12227 | PGF_0003 | YP_04931 | 893 | | Phosphotransferase RcsD |
| fig\|218491 | RcsF | Pectobacterium atrosepticum SCRI1043 | 218491,5 | NC_004547 | ECA3527 | VBIPecAtr | PATRIC.21 | PATRIC | CDS | 3955024 | 3955434 | 411 | + | FIG013040 | PLF_12227 | PGF_0003 | YP_05161 | 136 | rcsF | Protein RcsF |
| fig\|218491 | IgaA | Pectobacterium atrosepticum SCRI1043 | 218491,5 | NC_004547 | ECA4102 | VBIPecAtr | PATRIC.21 | PATRIC | CDS | 4570831 | 4572978 | 2148 | + | FIG000040 | PLF_12227 | PGF_0001 | YP_05218 | 715 | igaA | IgaA: a membrane protein that prevents overactivation of the Rcs regulatory system |

| | | | | | | | | | | | | | | | | | | | | | |
|---|---|---|---|---|---|---|---|---|---|---|---|---|---|---|---|---|---|---|---|---|---|
| fig\|561230 | DnaK | Pectobacterium carotovorum subsp. carotovorum PC1 | 561230.3 | NC_012917 | PC1_3659 | VBIPecCar | PATRIC.56 | PATRIC | CDS | 4128700 | 4130607 | 1908 | - | FIG000233 | PLF_1222 | PGF_1035 | YP_00301 | 635 | | Chaperone protein DnaK | |
| fig\|561230 | RcsC | Pectobacterium carotovorum subsp. carotovorum PC1 | 561230,3 | NC_012917 | PC1_1106 | VBIPecCar | PATRIC.56 | PATRIC | CDS | 1292123 | 1294978 | 2856 | + | FIG000042 | PLF_1222 | PGF_0005 | YP_00301 | 951 | | Sensor his | GO:0004673\|protein histidine kinase activity |
| fig\|561230 | RcsB | Pectobacterium carotovorum subsp. carotovorum PC1 | 561230,3 | NC_012917 | PC1_1107 | VBIPecCar | PATRIC.56 | PATRIC | CDS | 1295016 | 1295666 | 651 | - | FIG000041 | PLF_1222 | PGF_0042 | YP_00301 | 216 | | DNA-binding capsular synthesis response regulator RcsB | |
| fig\|561230 | RcsD | Pectobacterium carotovorum subsp. carotovorum PC1 | 561230,3 | NC_012917 | PC1_1108 | VBIPecCar | PATRIC.56 | PATRIC | CDS | 1295675 | 1298362 | 2688 | - | FIG000045 | PLF_1222 | PGF_0003 | YP_00301 | 895 | | Phosphotransferase RcsD | |
| fig\|561230 | RcsF | Pectobacterium carotovorum subsp. carotovorum PC1 | 561230,3 | NC_012917 | PC1_3345 | VBIPecCar | PATRIC.56 | PATRIC | CDS | 3773329 | 3773739 | 411 | + | FIG013040 | PLF_1222 | PGF_0003 | YP_00301 | 136 | | Protein RcsF | |
| fig\|561230 | IgaA | Pectobacterium carotovorum subsp. carotovorum PC1 | 561230,3 | NC_012917 | PC1_3892 | VBIPecCar | PATRIC.56 | PATRIC | CDS | 4377679 | 4379826 | 2148 | + | FIG000040 | PLF_1222 | PGF_0001 | YP_00301 | 715 | | IgaA: a membrane protein that prevents overactivation of the Rcs regulatory system | |
| fig\|561231 | DnaK | Pectobacterium wasabiae WPP163 | 561231.5 | NC_013421 | Pecwa_38 | VBIPecWa | PATRIC.56 | PATRIC | CDS | 4254388 | 4256295 | 1908 | - | FIG000233 | PLF_1222 | PGF_1035 | YP_00326 | 635 | | Chaperone protein DnaK | |
| fig\|561231 | RcsD | Pectobacterium wasabiae WPP163 | 561231,5 | NC_013421 | Pecwa_32 | VBIPecWa | PATRIC.56 | PATRIC | CDS | 3496409 | 3499096 | 2688 | + | FIG000045 | PLF_1222 | PGF_0003 | YP_00326 | 895 | | Phosphotransferase RcsD | |
| fig\|561231 | RcsB | Pectobacterium wasabiae WPP163 | 561231,5 | NC_013421 | Pecwa_32 | VBIPecWa | PATRIC.56 | PATRIC | CDS | 3499105 | 3499755 | 651 | + | FIG000041 | PLF_1222 | PGF_0042 | YP_00326 | 216 | | DNA-binding capsular synthesis response regulator RcsB | |
| fig\|561231 | RcsC | Pectobacterium wasabiae WPP163 | 561231,5 | NC_013421 | Pecwa_32 | VBIPecWa | PATRIC.56 | PATRIC | CDS | 3499793 | 3502648 | 2856 | - | FIG000042 | PLF_1222 | PGF_0005 | YP_00326 | 951 | | Sensor his | GO:0004673\|protein histidine kinase activity |
| fig\|561231 | RcsF | Pectobacterium wasabiae WPP163 | 561231,5 | NC_013421 | Pecwa_35 | VBIPecWa | PATRIC.56 | PATRIC | CDS | 3852200 | 3852610 | 411 | + | FIG013040 | PLF_1222 | PGF_0003 | YP_00326 | 136 | | Protein RcsF | |
| fig\|561231 | IgaA | Pectobacterium wasabiae WPP163 | 561231,5 | NC_013421 | Pecwa_40 | VBIPecWa | PATRIC.56 | PATRIC | CDS | 4477957 | 4480104 | 2148 | + | FIG000040 | PLF_1222 | PGF_0001 | YP_00326 | 715 | | IgaA: a membrane protein that prevents overactivation of the Rcs regulatory system | |
| fig\|291112 | DnaK | Photorhabdus asymbiotica strain ATCC 43949 | 291112.3 | NC_012962 | PAU_0054 | VBIPhoAs | PATRIC.29 | PATRIC | CDS | 609902 | 611812 | 1911 | + | FIG000233 | PLF_2948 | PGF_1035 | YP_00303 | 636 | dnaK | Chaperone protein DnaK | |
| fig\|291112 | IgaA | Photorhabdus asymbiotica strain ATCC 43949 | 291112,3 | NC_012962 | PAU_0007 | VBIPhoAs | PATRIC.29 | PATRIC | CDS | 81871 | 82884 | 1014 | + | FIG000040 | PLF_2948 | PGF_0001 | YP_00303 | 337 | yrfF | IgaA: a membrane protein that prevents overactivation of the Rcs regulatory system | |
| fig\|291112 | IgaA | Photorhabdus asymbiotica strain ATCC 43949 | 291112,3 | NC_012962 | PAU_0008 | VBIPhoAs | PATRIC.29 | PATRIC | CDS | 82884 | 83981 | 1098 | + | FIG000040 | PLF_2948 | PGF_0001 | YP_00303 | 365 | yrfF | IgaA: a membrane protein that prevents overactivation of the Rcs regulatory system | |
| fig\|291112 | RcsC | Photorhabdus asymbiotica strain ATCC 43949 | 291112,3 | NC_012962 | PAU_0155 | VBIPhoAs | PATRIC.29 | PATRIC | CDS | 1790947 | 1793778 | 2832 | + | FIG000042 | PLF_2948 | PGF_0005 | YP_00304 | 943 | rcsC | Sensor his | GO:0004673\|protein histidine kinase activity |
| fig\|291112 | RcsB | Photorhabdus asymbiotica strain ATCC 43949 | 291112,3 | NC_012962 | PAU_0156 | VBIPhoAs | PATRIC.29 | PATRIC | CDS | 1793928 | 1794581 | 654 | - | FIG000041 | PLF_2948 | PGF_0042 | YP_00304 | 217 | rcsB | DNA-binding capsular synthesis response regulator RcsB | |
| fig\|291112 | RcsD | Photorhabdus asymbiotica strain ATCC 43949 | 291112,3 | NC_012962 | PAU_0156 | VBIPhoAs | PATRIC.29 | PATRIC | CDS | 1794585 | 1797275 | 2691 | - | FIG000045 | PLF_2948 | PGF_0003 | YP_00304 | 896 | yojN | Phosphotransferase RcsD | |
| fig\|291112 | RcsF | Photorhabdus asymbiotica strain ATCC 43949 | 291112,3 | NC_012962 | PAU_0066 | VBIPhoAs | PATRIC.29 | PATRIC | CDS | 755599 | 755946 | 348 | - | FIG013040 | PLF_2948 | PGF_0003 | YP_00303 | 115 | rcsF | Protein RcsF | |
| fig\|243265 | DnaK | Photorhabdus luminescens subsp. laumondii TTO1 | 243265.5 | NC_005126 | plu0579 | VBIPhoLu | PATRIC.24 | PATRIC | CDS | 655919 | 657829 | 1911 | + | FIG000233 | PLF_2948 | PGF_1035 | NP_92792 | 636 | dnaK | Chaperone protein DnaK | |
| fig\|243265 | RcsD | Photorhabdus luminescens subsp. laumondii TTO1 | 243265,5 | NC_005126 | plu3047 | VBIPhoLu | PATRIC.24 | PATRIC | CDS | 3546077 | 3548767 | 2691 | + | FIG000045 | PLF_2948 | PGF_0003 | NP_93027 | 896 | | Phosphotransferase RcsD | |
| fig\|243265 | RcsB | Photorhabdus luminescens subsp. laumondii TTO1 | 243265,5 | NC_005126 | plu3048 | VBIPhoLu | PATRIC.24 | PATRIC | CDS | 3548772 | 3549425 | 654 | + | FIG000041 | PLF_2948 | PGF_0042 | NP_93028 | 217 | rcsB | DNA-binding capsular synthesis response regulator RcsB | |
| fig\|243265 | RcsC | Photorhabdus luminescens subsp. laumondii TTO1 | 243265,5 | NC_005126 | plu3049 | VBIPhoLu | PATRIC.24 | PATRIC | CDS | 3549510 | 3552257 | 2748 | - | FIG000042 | PLF_2948 | PGF_0005 | NP_93028 | 915 | rcsC | Sensor his | GO:0004673\|protein histidine kinase activity |
| fig\|243265 | RcsF | Photorhabdus luminescens subsp. laumondii TTO1 | 243265,5 | NC_005126 | plu0694 | VBIPhoLu | PATRIC.24 | PATRIC | CDS | 797711 | 798109 | 399 | - | FIG013040 | PLF_2948 | PGF_0003 | NP_92803 | 132 | rcsF | Protein RcsF | |
| fig\|243265 | IgaA | Photorhabdus luminescens subsp. laumondii TTO1 | 243265,5 | NC_005126 | plu0097 | VBIPhoLu | PATRIC.24 | PATRIC | CDS | 92664 | 94796 | 2133 | + | FIG000040 | PLF_2948 | PGF_0001 | NP_92746 | 710 | | IgaA: a membrane protein that prevents overactivation of the Rcs regulatory system | |
| fig\|230089 | DnaK | Photorhabdus temperata subsp. thracensis strain DSM 1 | 230089.6 | CP011104 | VY86_04980 | | PATRIC.23 | PATRIC | CDS | 1268611 | 1270518 | 1908 | - | FIG000233 | PLF_2948 | PGF_10357457 | | 635 | dnaK | Chaperone protein DnaK | |
| fig\|230089 | IgaA | Photorhabdus temperata subsp. thracensis strain DSM 1 | 230089,6 | CP011104 | VY86_08385 | | PATRIC.23 | PATRIC | CDS | 1971098 | 1973212 | 2115 | + | FIG000040 | PLF_2948 | PGF_00013790 | | 704 | | IgaA: a membrane protein that prevents overactivation of the Rcs regulatory system | |
| fig\|230089 | RcsC | Photorhabdus temperata subsp. thracensis strain DSM 1 | 230089,6 | CP011104 | VY86_16435 | | PATRIC.23 | PATRIC | CDS | 3827334 | 3830171 | 2838 | + | FIG000042 | PLF_2948 | PGF_00050577 | | 945 | | Sensor his | GO:0004673\|protein histidine kinase activity |
| fig\|230089 | RcsB | Photorhabdus temperata subsp. thracensis strain DSM 1 | 230089,6 | CP011104 | VY86_16440 | | PATRIC.23 | PATRIC | CDS | 3830240 | 3830893 | 654 | - | FIG000041 | PLF_2948 | PGF_00421954 | | 217 | | DNA-binding capsular synthesis response regulator RcsB | |
| fig\|230089 | RcsD | Photorhabdus temperata subsp. thracensis strain DSM 1 | 230089,6 | CP011104 | VY86_16445 | | PATRIC.23 | PATRIC | CDS | 3830899 | 3833589 | 2691 | - | FIG000045 | PLF_2948 | PGF_00034034 | | 896 | | Phosphotransferase RcsD | |
| fig\|230089 | RcsF | Photorhabdus temperata subsp. thracensis strain DSM 1 | 230089,6 | CP011104 | VY86_04255 | | PATRIC.23 | PATRIC | CDS | 1066043 | 1066441 | 399 | + | FIG013040 | PLF_2948 | PGF_00037945 | | 132 | | Protein RcsF | |
| fig\|703.8.p | DnaK | Plesiomonas shigelloides strain NCTC10360 | 703.8 | LT575468 | SAMEA2665130_024 | | PATRIC.70 | PATRIC | CDS | 2679898 | 2681817 | 1920 | - | FIG000233 | PLF_702_ | PGF_10357457 | | 639 | dnaK_2 | Chaperone protein DnaK | |
| fig\|703.8.p | RcsF | Plesiomonas shigelloides strain NCTC10360 | 703,8 | LT575468 | SAMEA2665130_024 | | PATRIC.70 | PATRIC | CDS | 2700025 | 2700585 | 561 | + | | PLF_702_ | PGF_04774977 | | 186 | rcsF | hypothetical protein | |
| fig\|703.8.p | RcsC | Plesiomonas shigelloides strain NCTC10360 | 703,8 | LT575468 | SAMEA2665130_028 | | PATRIC.70 | PATRIC | CDS | 3090011 | 3093493 | 3483 | - | FIG004509 | PLF_702_ | PGF_02620852 | | 1160 | rcsC | Two-component system sensor histidine kinase | |
| fig\|61647.1 | DnaK | Pluralibacter gergoviae FB2 | 61647.10 | CP009450 | | | PATRIC.61 | PATRIC | CDS | 5342210 | 5344126 | 1917 | - | FIG000233 | PLF_1330 | PGF_10357457 | | 638 | | Chaperone protein DnaK | |
| fig\|61647.1 | RcsC | Pluralibacter gergoviae FB2 | 61647,1 | CP009450 | | | PATRIC.61 | PATRIC | CDS | 2544790 | 2547624 | 2835 | + | FIG000042 | PLF_1330 | PGF_00050577 | | 944 | | Sensor his | GO:0004673\|protein histidine kinase activity |
| fig\|61647.1 | RcsB | Pluralibacter gergoviae FB2 | 61647,1 | CP009450 | | | PATRIC.61 | PATRIC | CDS | 2547683 | 2548333 | 651 | - | FIG000041 | PLF_1330 | PGF_00421954 | | 216 | | DNA-binding capsular synthesis response regulator RcsB | |
| fig\|61647.1 | RcsD | Pluralibacter gergoviae FB2 | 61647,1 | CP009450 | | | PATRIC.61 | PATRIC | CDS | 2548350 | 2551025 | 2676 | - | FIG000045 | PLF_1330 | PGF_00034034 | | 891 | | Phosphotransferase RcsD | |
| fig\|61647.1 | RcsF | Pluralibacter gergoviae FB2 | 61647,1 | CP009450 | | | PATRIC.61 | PATRIC | CDS | 5142063 | 5142467 | 405 | + | FIG013040 | PLF_1330 | PGF_00037945 | | 134 | | Protein RcsF | |
| fig\|61647.1 | IgaA | Pluralibacter gergoviae FB2 | 61647,1 | CP009450 | | | PATRIC.61 | PATRIC | CDS | 609555 | 611702 | 2148 | + | FIG000040 | PLF_1330 | PGF_00013790 | | 715 | | IgaA: a membrane protein that prevents overactivation of the Rcs regulatory system | |
| fig\|529507 | DnaK | Proteus mirabilis HI4320 | 529507.6 | NC_010554 | PMI0009 | VBIProMir1 | PATRIC.52 | PATRIC | CDS | 18101 | 20026 | 1926 | + | FIG000233 | PLF_583_ | PGF_1035 | YP_00214 | 641 | dnaK | Chaperone protein DnaK | |
| fig\|529507 | RcsD | Proteus mirabilis HI4320 | 529507,6 | NC_010554 | PMI1729 | VBIProMir1 | PATRIC.52 | PATRIC | CDS | 1845004 | 1847697 | 2694 | + | FIG000045 | PLF_583_ | PGF_0003 | YP_00215 | 897 | rsbA | Phosphotransferase RcsD | |
| fig\|529507 | RcsB | Proteus mirabilis HI4320 | 529507,6 | NC_010554 | PMI1730 | VBIProMir1 | PATRIC.52 | PATRIC | CDS | 1847715 | 1848371 | 657 | + | FIG000041 | PLF_583_ | PGF_0042 | YP_00215 | 218 | rcsB | DNA-binding capsular synthesis response regulator RcsB | |
| fig\|529507 | RcsC | Proteus mirabilis HI4320 | 529507,6 | NC_010554 | PMI1731 | VBIProMir1 | PATRIC.52 | PATRIC | CDS | 1848619 | 1851378 | 2760 | - | FIG000042 | PLF_583_ | PGF_0005 | YP_00215 | 919 | rcsC | Sensor his | GO:0004673\|protein histidine kinase activity |
| fig\|529507 | RcsF | Proteus mirabilis HI4320 | 529507,6 | NC_010554 | PMI2262 | VBIProMir1 | PATRIC.52 | PATRIC | CDS | 2457939 | 2458343 | 405 | + | FIG013040 | PLF_583_ | PGF_0003 | YP_00215 | 134 | rcsF | Protein RcsF | |
| fig\|529507 | IgaA | Proteus mirabilis HI4320 | 529507,6 | NC_010554 | PMI3018 | VBIProMir1 | PATRIC.52 | PATRIC | CDS | 3312777 | 3314834 | 2058 | - | FIG000040 | PLF_583_ | PGF_0001 | YP_00215 | 685 | umoB | IgaA: a membrane protein that prevents overactivation of the Rcs regulatory system | |
| fig\|520999 | DnaK | Providencia alcalifaciens DSM 30120 | 520999.6 | NZ_ABXW0100004 | PROVALC | VBIProAlc1 | PATRIC.52 | PATRIC | CDS | 73469 | 75382 | 1914 | + | FIG000233 | PLF_586_ | PGF_1035 | ZP_03317 | 637 | | Chaperone protein DnaK | |
| fig\|520999 | RcsF | Providencia alcalifaciens DSM 30120 | 520999,6 | NZ_ABXW0100004 | PROVALC | VBIProAlc1 | PATRIC.52 | PATRIC | CDS | 3582 | 3893 | 312 | + | FIG013040 | PLF_586_ | PGF_0003 | ZP_03318 | 103 | | Protein RcsF | |
| fig\|520999 | RcsC | Providencia alcalifaciens DSM 30120 | 520999,6 | NZ_ABXW0100006 | PROVALC | VBIProAlc1 | PATRIC.52 | PATRIC | CDS | 128789 | 131575 | 2787 | + | FIG000042 | PLF_586_ | PGF_0005 | ZP_03320 | 928 | | Sensor his | GO:0004673\|protein histidine kinase activity |
| fig\|520999 | RcsB | Providencia alcalifaciens DSM 30120 | 520999,6 | NZ_ABXW0100006 | PROVALC | VBIProAlc1 | PATRIC.52 | PATRIC | CDS | 131630 | 132280 | 651 | - | FIG000041 | PLF_586_ | PGF_0042 | ZP_03320 | 216 | | DNA-binding capsular synthesis response regulator RcsB | |
| fig\|520999 | RcsD | Providencia alcalifaciens DSM 30120 | 520999,6 | NZ_ABXW0100006 | PROVALC | VBIProAlc1 | PATRIC.52 | PATRIC | CDS | 132282 | 134975 | 2694 | - | FIG000045 | PLF_586_ | PGF_0003 | ZP_03320 | 897 | | Phosphotransferase RcsD | |
| fig\|520999 | IgaA | Providencia alcalifaciens DSM 30120 | 520999,6 | NZ_ABXW0100005 | PROVALC | VBIProAlc1 | PATRIC.52 | PATRIC | CDS | 23068 | 24903 | 1836 | + | FIG000040 | PLF_586_ | PGF_0001 | ZP_03317 | 611 | | IgaA: a membrane protein that prevents overactivation of the Rcs regulatory system | |
| fig\|114166 | DnaK | Providencia burhodogranariea DSM 19968 | 1141662.3 | AKKL01000046 | OOA_1657 | VBIProBur | PATRIC.11 | PATRIC | CDS | 117475 | 119388 | 1914 | + | FIG000233 | PLF_586_ | PGF_1035 | EKT55424 | 637 | dnaK | Chaperone protein DnaK | |
| fig\|114166 | RcsF | Providencia burhodogranariea DSM 19968 | 1141662,3 | AKKL01000037 | OOA_1412 | VBIProBur | PATRIC.11 | PATRIC | CDS | 196150 | 196554 | 405 | - | FIG013040 | PLF_586_ | PGF_0003 | EKT58269 | 134 | rcsF | Protein RcsF | |
| fig\|114166 | IgaA | Providencia burhodogranariea DSM 19968 | 1141662,3 | AKKL01000045 | OOA_1597 | VBIProBur | PATRIC.11 | PATRIC | CDS | 130607 | 132565 | 1959 | - | FIG000040 | PLF_586_ | PGF_0001 | EKT56126 | 652 | | IgaA: a membrane protein that prevents overactivation of the Rcs regulatory system | |
| fig\|114166 | RcsC | Providencia burhodogranariea DSM 19968 | 1141662,3 | AKKL01000012 | OOA_0353 | VBIProBur | PATRIC.11 | PATRIC | CDS | 143244 | 146081 | 2838 | + | FIG000042 | PLF_586_ | PGF_0005 | EKT63901 | 945 | | Sensor his | GO:0004673\|protein histidine kinase activity |
| fig\|114166 | RcsB | Providencia burhodogranariea DSM 19968 | 1141662,3 | AKKL01000012 | OOA_0354 | VBIProBur | PATRIC.11 | PATRIC | CDS | 146156 | 146806 | 651 | - | FIG000041 | PLF_586_ | PGF_0042 | EKT63902 | 216 | | DNA-binding capsular synthesis response regulator RcsB | |
| fig\|114166 | RcsD | Providencia burhodogranariea DSM 19968 | 1141662,3 | AKKL01000012 | OOA_0354 | VBIProBur | PATRIC.11 | PATRIC | CDS | 146808 | 149432 | 2625 | - | FIG000045 | PLF_586_ | PGF_0003 | EKT63903 | 874 | | Phosphotransferase RcsD | |
| fig\|114166 | DnaK | Providencia rettgeri Dmel1 | 1141663.3 | AJSB01000008 | OOC_1884 | VBIProRet | PATRIC.11 | PATRIC | CDS | 306372 | 308288 | 1917 | - | FIG000233 | PLF_586_ | PGF_1035 | EKT53968 | 638 | dnaK | Chaperone protein DnaK | |
| fig\|114166 | RcsF | Providencia rettgeri Dmel1 | 1141663,3 | AJSB01000004 | OOC_0676 | VBIProRet | PATRIC.11 | PATRIC | CDS | 809299 | 809700 | 402 | - | FIG013040 | PLF_586_ | PGF_0003 | EKT59351 | 133 | rcsF | Protein RcsF | |
| fig\|114166 | RcsD | Providencia rettgeri Dmel1 | 1141663,3 | AJSB01000007 | OOC_1395 | VBIProRet | PATRIC.11 | PATRIC | CDS | 194918 | 197542 | 2625 | + | FIG000045 | PLF_586_ | PGF_0003 | EKT54662 | 874 | | Phosphotransferase RcsD | |
| fig\|114166 | RcsB | Providencia rettgeri Dmel1 | 1141663,3 | AJSB01000007 | OOC_1396 | VBIProRet | PATRIC.11 | PATRIC | CDS | 197544 | 198194 | 651 | + | FIG000041 | PLF_586_ | PGF_0042 | EKT54663 | 216 | | DNA-binding capsular synthesis response regulator RcsB | |
| fig\|114166 | RcsC | Providencia rettgeri Dmel1 | 1141663,3 | AJSB01000007 | OOC_1396 | VBIProRet | PATRIC.11 | PATRIC | CDS | 198273 | 201116 | 2844 | - | FIG000042 | PLF_586_ | PGF_0005 | EKT54664 | 947 | | Sensor his | GO:0004673\|protein histidine kinase activity |
| fig\|114166 | IgaA | Providencia rettgeri Dmel1 | 1141663,3 | AJSB01000001 | OOC_0151 | VBIProRet | PATRIC.11 | PATRIC | CDS | 301825 | 303783 | 1959 | - | FIG000040 | PLF_586_ | PGF_0001 | EKT60653 | 652 | | IgaA: a membrane protein that prevents overactivation of the Rcs regulatory system | |
| fig\|114166 | DnaK | Providencia sneebia DSM 19967 | 1141660.3 | AKKN01000013 | OO7_1535 | VBIProSne | PATRIC.11 | PATRIC | CDS | 228074 | 229990 | 1917 | - | FIG000233 | PLF_586_ | PGF_1035 | EKT53599 | 638 | dnaK | Chaperone protein DnaK | |
| fig\|114166 | RcsD | Providencia sneebia DSM 19967 | 1141660,3 | AKKN01000010 | OO7_1139 | VBIProSne | PATRIC.11 | PATRIC | CDS | 80418 | 83126 | 2709 | + | FIG000045 | PLF_586_ | PGF_0003 | EKT55585 | 902 | | Phosphotransferase RcsD | |
| fig\|114166 | RcsB | Providencia sneebia DSM 19967 | 1141660,3 | AKKN01000010 | OO7_1139 | VBIProSne | PATRIC.11 | PATRIC | CDS | 83128 | 83790 | 663 | + | FIG000041 | PLF_586_ | PGF_0042 | EKT55586 | 220 | | DNA-binding capsular synthesis response regulator RcsB | |
| fig\|114166 | RcsC | Providencia sneebia DSM 19967 | 1141660,3 | AKKN01000010 | OO7_1140 | VBIProSne | PATRIC.11 | PATRIC | CDS | 83875 | 86712 | 2838 | - | FIG000042 | PLF_586_ | PGF_0005 | EKT55587 | 945 | | Sensor his | GO:0004673\|protein histidine kinase activity |
| fig\|114166 | RcsF | Providencia sneebia DSM 19967 | 1141660,3 | AKKN01000013 | OO7_1429 | VBIProSne | PATRIC.11 | PATRIC | CDS | 4253 | 4657 | 405 | + | FIG013040 | PLF_586_ | PGF_0003 | EKT53396 | 134 | rcsF | Protein RcsF | |
| fig\|114166 | IgaA | Providencia sneebia DSM 19967 | 1141660,3 | AKKN01000003 | OO7_0200 | VBIProSne | PATRIC.11 | PATRIC | CDS | 21322 | 23280 | 1959 | + | FIG000040 | PLF_586_ | PGF_0001 | EKT60831 | 652 | | IgaA: a membrane protein that prevents overactivation of the Rcs regulatory system | |
| fig\|115795 | DnaK | Providencia stuartii MRSN 2154 | 1157951.4 | CP003488 | S70_0724 | VBIProStu | PATRIC.11 | PATRIC | CDS | 1515546 | 1517465 | 1920 | - | FIG000233 | PLF_586_ | PGF_1035 | AFH93319 | 639 | dnaK | Chaperone protein DnaK | |
| fig\|115795 | IgaA | Providencia stuartii MRSN 2154 | 1157951,4 | CP003488 | S70_1004 | VBIProStu | PATRIC.11 | PATRIC | CDS | 2147168 | 2149126 | 1959 | - | FIG000040 | PLF_586_ | PGF_0001 | AFH93868 | 652 | | IgaA: a membrane protein that prevents overactivation of the Rcs regulatory system | |
| fig\|115795 | RcsF | Providencia stuartii MRSN 2154 | 1157951,4 | CP003488 | S70_1518 | VBIProStu | PATRIC.11 | PATRIC | CDS | 3226449 | 3226757 | 309 | - | FIG013040 | PLF_586_ | PGF_0003 | AFH94861 | 102 | rcsF | Protein RcsF | |
| fig\|115795 | RcsD | Providencia stuartii MRSN 2154 | 1157951,4 | CP003488 | S70_0224 | VBIProStu | PATRIC.11 | PATRIC | CDS | 437189 | 439900 | 2712 | + | FIG000045 | PLF_586_ | PGF_0003 | AFH92342 | 903 | | Phosphotransferase RcsD | |
| fig\|115795 | RcsB | Providencia stuartii MRSN 2154 | 1157951,4 | CP003488 | S70_0224 | VBIProStu | PATRIC.11 | PATRIC | CDS | 439902 | 440552 | 651 | + | FIG000041 | PLF_586_ | PGF_0042 | AFH92343 | 216 | | DNA-binding capsular synthesis response regulator RcsB | |
| fig\|115795 | RcsC | Providencia stuartii MRSN 2154 | 1157951,4 | CP003488 | S70_0225 | VBIProStu | PATRIC.11 | PATRIC | CDS | 440631 | 443468 | 2838 | - | FIG000042 | PLF_586_ | PGF_0005 | AFH92344 | 945 | | Sensor his | GO:0004673\|protein histidine kinase activity |
| fig\|133623 | DnaK | Pseudomonas flectens ATCC 12775 | 1336237.3 | JAEE01000001 | | VBIPseFle | PATRIC.13 | PATRIC | CDS | 620507 | 622414 | 1908 | - | FIG00023369 | | PGF_10357457 | | 635 | | Chaperone protein DnaK | |
| fig\|133623 | RcsD | Pseudomonas flectens ATCC 12775 | 1336237,3 | JAEE01000004 | | VBIPseFle | PATRIC.13 | PATRIC | CDS | 60709 | 63375 | 2667 | + | FIG00004592 | | PGF_00034034 | | 888 | | Phosphotransferase RcsD | |
| fig\|133623 | RcsB | Pseudomonas flectens ATCC 12775 | 1336237,3 | JAEE01000004 | | VBIPseFle | PATRIC.13 | PATRIC | CDS | 63368 | 64036 | 669 | + | FIG00004153 | | PGF_00421954 | | 222 | | DNA-binding capsular synthesis response regulator RcsB | |
| fig\|133623 | RcsC | Pseudomonas flectens ATCC 12775 | 1336237,3 | JAEE01000004 | | VBIPseFle | PATRIC.13 | PATRIC | CDS | 64322 | 67135 | 2814 | - | FIG00004208 | | PGF_00050577 | | 937 | | Sensor his | GO:0004673\|protein histidine kinase activity |
| fig\|133623 | IgaA | Pseudomonas flectens ATCC 12775 | 1336237,3 | JAEE01000009 | | VBIPseFle | PATRIC.13 | PATRIC | CDS | 68610 | 70742 | 2133 | - | FIG00004077 | | PGF_00013790 | | 710 | | IgaA: a membrane protein that prevents overactivation of the Rcs regulatory system | |
| fig\|133623 | RcsF | Pseudomonas flectens ATCC 12775 | 1336237,3 | JAEE01000016 | | VBIPseFle | PATRIC.13 | PATRIC | CDS | 39911 | 40315 | 405 | - | | | | | 134 | | Protein RcsF | |
| fig\|745277 | DnaK | Rahnella aquatilis CIP 78.65 = ATCC 33071 | 745277.3 | CP003244 | Rahaq2_3 | VBIRahAq | PATRIC.74 | PATRIC | CDS | 4134990 | 4136891 | 1902 | - | FIG000233 | PLF_3403 | PGF_1035 | AEX53667 | 633 | | Chaperone protein DnaK | |
| fig\|745277 | RcsC | Rahnella aquatilis CIP 78.65 = ATCC 33071 | 745277,3 | CP003244 | Rahaq2_1 | VBIRahAq | PATRIC.74 | PATRIC | CDS | 1416776 | 1419562 | 2787 | + | FIG000042 | PLF_3403 | PGF_0005 | AEX51246 | 928 | | Sensor his | GO:0004673\|protein histidine kinase activity |
| fig\|745277 | RcsB | Rahnella aquatilis CIP 78.65 = ATCC 33071 | 745277,3 | CP003244 | Rahaq2_1 | VBIRahAq | PATRIC.74 | PATRIC | CDS | 1420543 | 1421193 | 651 | - | FIG000041 | PLF_3403 | PGF_0042 | AEX51248 | 216 | | DNA-binding capsular synthesis response regulator RcsB | |
| fig\|745277 | RcsD | Rahnella aquatilis CIP 78.65 = ATCC 33071 | 745277,3 | CP003244 | Rahaq2_1 | VBIRahAq | PATRIC.74 | PATRIC | CDS | 1421198 | 1423891 | 2694 | - | FIG000045 | PLF_3403 | PGF_0003 | AEX51249 | 897 | | Phosphotransferase RcsD | |

| | | | | | | | | | | | | | | | | | | | | | |
|---|---|---|---|---|---|---|---|---|---|---|---|---|---|---|---|---|---|---|---|---|---|
| fig\|745277 | IgaA | Rahnella aquatilis CIP 78.65 = ATCC 33071 | 745277,3 | CP003244 | Rahaq2_0 | VBIRahAq | PATRIC.74 | PATRIC | CDS | 303419 | 305566 | 2148 | - | FIG000040 | PLF_3403 | PGF_0001 | AEX50226 | 715 | | IgaA: a membrane protein that prevents overactivation of the Rcs regulatory system | |
| fig\|745277 | RcsF | Rahnella aquatilis CIP 78.65 = ATCC 33071 | 745277,3 | CP003244 | Rahaq2_0 | VBIRahAq | PATRIC.74 | PATRIC | CDS | 998321 | 998728 | 408 | - | FIG013040 | PLF_3403 | PGF_0003 | AEX50854 | 135 | | Protein RcsF | |
| fig\|128617 | DnaK | Raoultella ornithinolytica B6 | 1286170.3 | CP004142 | RORB6_15 | VBIRaoOrn | PATRIC.12 | PATRIC | CDS | 3326621 | 3328537 | 1917 | - | FIG000233 | PLF_1606 | PGF_1035 | AGJ87712 | 638 | dnaK | Chaperone protein DnaK | |
| fig\|128617 | RcsC | Raoultella ornithinolytica B6 | 1286170,3 | CP004142 | RORB6_01 | VBIRaoOrn | PATRIC.12 | PATRIC | CDS | 294828 | 297683 | 2856 | + | FIG000042 | PLF_1606 | PGF_0005 | AGJ84961 | 951 | | Sensor his | GO:0004673\|protein histidine kinase activity |
| fig\|128617 | RcsB | Raoultella ornithinolytica B6 | 1286170,3 | CP004142 | RORB6_01 | VBIRaoOrn | PATRIC.12 | PATRIC | CDS | 297747 | 298397 | 651 | - | FIG000041 | PLF_1606 | PGF_0042 | AGJ84962 | 216 | | DNA-binding capsular synthesis response regulator RcsB | |
| fig\|128617 | RcsD | Raoultella ornithinolytica B6 | 1286170,3 | CP004142 | RORB6_01 | VBIRaoOrn | PATRIC.12 | PATRIC | CDS | 298414 | 301071 | 2658 | - | FIG000045 | PLF_1606 | PGF_0003 | AGJ84963 | 885 | | Phosphotransferase RcsD | |
| fig\|128617 | RcsF | Raoultella ornithinolytica B6 | 1286170,3 | CP004142 | RORB6_14 | VBIRaoOrn | PATRIC.12 | PATRIC | CDS | 3093854 | 3094261 | 408 | + | FIG013040 | PLF_1606 | PGF_0003 | AGJ87507 | 135 | rcsF | Protein RcsF | |
| fig\|128617 | IgaA | Raoultella ornithinolytica B6 | 1286170,3 | CP004142 | RORB6_20 | VBIRaoOrn | PATRIC.12 | PATRIC | CDS | 4556952 | 4559087 | 2136 | - | FIG000040 | PLF_1606 | PGF_0001 | AGJ88797 | 711 | | IgaA: a membrane protein that prevents overactivation of the Rcs regulatory system | |
| fig\|218493 | DnaK | Salmonella bongori NCTC 12419 | 218493.5 | NC_015761 | SBG_0013 | VBISalBon | PATRIC.21 | PATRIC | CDS | 12050 | 13966 | 1917 | + | FIG000233 | PLF_590_ | PGF_1035 | YP_00472 | 638 | dnaK | Chaperone protein DnaK | |
| fig\|218493 | RcsD | Salmonella bongori NCTC 12419 | 218493,5 | NC_015761 | SBG_2066 | VBISalBon | PATRIC.21 | PATRIC | CDS | 2251455 | 2254124 | 2670 | + | FIG000045 | PLF_590_ | PGF_0003 | YP_00473 | 889 | yojN | Phosphotransferase RcsD | |
| fig\|218493 | RcsB | Salmonella bongori NCTC 12419 | 218493,5 | NC_015761 | SBG_2067 | VBISalBon | PATRIC.21 | PATRIC | CDS | 2254141 | 2254791 | 651 | + | FIG000041 | PLF_590_ | PGF_0042 | YP_00473 | 216 | rcsB | DNA-binding capsular synthesis response regulator RcsB | |
| fig\|218493 | RcsC | Salmonella bongori NCTC 12419 | 218493,5 | NC_015761 | SBG_2068 | VBISalBon | PATRIC.21 | PATRIC | CDS | 2254895 | 2257741 | 2847 | - | FIG000042 | PLF_590_ | PGF_0005 | YP_00473 | 948 | rcsC | Sensor his | GO:0004673\|protein histidine kinase activity |
| fig\|218493 | RcsF | Salmonella bongori NCTC 12419 | 218493,5 | NC_015761 | SBG_0235 | VBISalBon | PATRIC.21 | PATRIC | CDS | 268291 | 268422 | 132 | - | FIG013040 | PLF_590_ | PGF_0003 | YP_00472 | 43 | rcsF | Protein RcsF | |
| fig\|218493 | IgaA | Salmonella bongori NCTC 12419 | 218493,5 | NC_015761 | SBG_3100 | VBISalBon | PATRIC.21 | PATRIC | CDS | 3380691 | 3382823 | 2133 | + | FIG000040 | PLF_590_ | PGF_0001 | YP_00473 | 710 | yrfF | IgaA: a membrane protein that prevents overactivation of the Rcs regulatory system | |
| fig\|41514.7 | DnaK | Salmonella enterica subsp. arizonae serovar 62:z4,z23:- | 41514.7 | NC_010067 | SARI_0298 | VBISalEnt | PATRIC.41 | PATRIC | CDS | 2912205 | 2914121 | 1917 | - | FIG000233 | PLF_590_ | PGF_1035 | YP_00157 | 638 | dnaK | Chaperone protein DnaK | |
| fig\|41514.7 | RcsF | Salmonella enterica subsp. arizonae serovar 62:z4,z23:- | 41514,7 | NC_010067 | SARI_0275 | VBISalEnt | PATRIC.41 | PATRIC | CDS | 2668169 | 2668540 | 372 | + | FIG013040 | PLF_590_ | PGF_0003 | YP_00157 | 123 | rcsF | Protein RcsF | |
| fig\|41514.7 | IgaA | Salmonella enterica subsp. arizonae serovar 62:z4,z23:- | 41514,7 | NC_010067 | SARI_0412 | VBISalEnt | PATRIC.41 | PATRIC | CDS | 4071183 | 4073225 | 2043 | - | FIG000040 | PLF_590_ | PGF_0001 | YP_00157 | 680 | | IgaA: a membrane protein that prevents overactivation of the Rcs regulatory system | |
| fig\|41514.7 | RcsC | Salmonella enterica subsp. arizonae serovar 62:z4,z23:- | 41514,7 | NC_010067 | SARI_006 | VBISalEnt | PATRIC.41 | PATRIC | CDS | 621496 | 624342 | 2847 | + | FIG000042 | PLF_590_ | PGF_0005 | YP_00156 | 948 | | Sensor his | GO:0004673\|protein histidine kinase activity |
| fig\|41514.7 | RcsB | Salmonella enterica subsp. arizonae serovar 62:z4,z23:- | 41514,7 | NC_010067 | SARI_006 | VBISalEnt | PATRIC.41 | PATRIC | CDS | 624444 | 625094 | 651 | - | FIG000041 | PLF_590_ | PGF_0042 | YP_00156 | 216 | | DNA-binding capsular synthesis response regulator RcsB | |
| fig\|41514.7 | RcsD | Salmonella enterica subsp. arizonae serovar 62:z4,z23:- | 41514,7 | NC_010067 | | VBISalEnt | PATRIC.41 | PATRIC | CDS | 625111 | 626268 | 1158 | - | FIG000045 | PLF_590_ | PGF_00034034 | | 385 | | Phosphotransferase RcsD | |
| fig\|41514.7 | RcsD | Salmonella enterica subsp. arizonae serovar 62:z4,z23:- | 41514,7 | NC_010067 | | VBISalEnt | PATRIC.41 | PATRIC | CDS | 626301 | 627779 | 1479 | - | FIG000045 | PLF_590_ | PGF_00034034 | | 492 | | Phosphotransferase RcsD | |
| fig\|295319 | DnaK | Salmonella enterica subsp. enterica serovar Paratyphi A | 295319.15 | NC_006511 | SPA0012 | VBISalEnt | PATRIC.29 | PATRIC | CDS | 11594 | 13510 | 1917 | + | FIG000233 | PLF_590_ | PGF_1035 | YP_14936 | 638 | dnaK | Chaperone protein DnaK | |
| fig\|295319 | RcsF | Salmonella enterica subsp. enterica serovar Paratyphi A | 295319,15 | NC_006511 | SPA0251 | VBISalEnt | PATRIC.29 | PATRIC | CDS | 289140 | 289544 | 405 | - | FIG013040 | PLF_590_ | PGF_0003 | YP_14959 | 134 | rcsF | Protein RcsF | |
| fig\|295319 | IgaA | Salmonella enterica subsp. enterica serovar Paratyphi A | 295319,15 | NC_006511 | SPA3360 | VBISalEnt | PATRIC.29 | PATRIC | CDS | 3455897 | 3458029 | 2133 | + | FIG000040 | PLF_590_ | PGF_0001 | YP_15248 | 710 | yrfF | IgaA: a membrane protein that prevents overactivation of the Rcs regulatory system | |
| fig\|295319 | RcsC | Salmonella enterica subsp. enterica serovar Paratyphi A | 295319,15 | NC_006511 | SPA0593 | VBISalEnt | PATRIC.29 | PATRIC | CDS | 671675 | 674521 | 2847 | + | FIG000042 | PLF_590_ | PGF_0005 | YP_14990 | 948 | rcsC | Sensor his | GO:0004673\|protein histidine kinase activity |
| fig\|295319 | RcsB | Salmonella enterica subsp. enterica serovar Paratyphi A | 295319,15 | NC_006511 | SPA0594 | VBISalEnt | PATRIC.29 | PATRIC | CDS | 674624 | 675274 | 651 | - | FIG000041 | PLF_590_ | PGF_0042 | YP_14990 | 216 | rcsB | DNA-binding capsular synthesis response regulator RcsB | |
| fig\|295319 | RcsD | Salmonella enterica subsp. enterica serovar Paratyphi A | 295319,15 | NC_006511 | SPA0595 | VBISalEnt | PATRIC.29 | PATRIC | CDS | 675291 | 677960 | 2670 | - | FIG000045 | PLF_590_ | PGF_0003 | YP_14990 | 889 | yojN | Phosphotransferase RcsD | |
| fig\|220341 | DnaK | Salmonella enterica subsp. enterica serovar Typhi str. CT | 220341.7 | NC_003198 | STY0012 | VBISalEnt | PATRIC.22 | PATRIC | CDS | 11594 | 13510 | 1917 | + | FIG000233 | PLF_590_ | PGF_1035 | NP_45462 | 638 | dnaK | Chaperone protein DnaK | |
| fig\|220341 | RcsD | Salmonella enterica subsp. enterica serovar Typhi str. CT | 220341,7 | NC_003198 | STY2494 | VBISalEnt | PATRIC.22 | PATRIC | CDS | 2324258 | 2326927 | 2670 | + | FIG000045 | PLF_590_ | PGF_0003 | NP_45681 | 889 | yojN | Phosphotransferase RcsD | |
| fig\|220341 | RcsB | Salmonella enterica subsp. enterica serovar Typhi str. CT | 220341,7 | NC_003198 | STY2495 | VBISalEnt | PATRIC.22 | PATRIC | CDS | 2326953 | 2327594 | 642 | + | FIG000041 | PLF_590_ | PGF_0042 | NP_45681 | 213 | rcsB | DNA-binding capsular synthesis response regulator RcsB | |
| fig\|220341 | RcsC | Salmonella enterica subsp. enterica serovar Typhi str. CT | 220341,7 | NC_003198 | STY2496 | VBISalEnt | PATRIC.22 | PATRIC | CDS | 2327697 | 2330543 | 2847 | - | FIG000042 | PLF_590_ | PGF_0005 | NP_45681 | 948 | rcsC | Sensor his | GO:0004673\|protein histidine kinase activity |
| fig\|220341 | RcsF | Salmonella enterica subsp. enterica serovar Typhi str. CT | 220341,7 | NC_003198 | STY0271 | VBISalEnt | PATRIC.22 | PATRIC | CDS | 283298 | 283702 | 405 | - | FIG013040 | PLF_590_ | PGF_0003 | NP_45485 | 134 | rcsF | Protein RcsF | |
| fig\|220341 | IgaA | Salmonella enterica subsp. enterica serovar Typhi str. CT | 220341,7 | NC_003198 | STY4301 | VBISalEnt | PATRIC.22 | PATRIC | CDS | 4178276 | 4180408 | 2133 | - | FIG000040 | PLF_590_ | PGF_0001 | NP_45840 | 710 | yrfF | IgaA: a membrane protein that prevents overactivation of the Rcs regulatory system | |
| fig\|209261 | DnaK | Salmonella enterica subsp. enterica serovar Typhi str. Ty | 209261.6 | NC_004631 | t0012 | VBISalEnt | PATRIC.20 | PATRIC | CDS | 11594 | 13510 | 1917 | + | FIG000233 | PLF_590_ | PGF_1035 | NP_80389 | 638 | dnaK | Chaperone protein DnaK | |
| fig\|209261 | RcsF | Salmonella enterica subsp. enterica serovar Typhi str. Ty | 209261,6 | NC_004631 | t0247 | VBISalEnt | PATRIC.20 | PATRIC | CDS | 283289 | 283642 | 354 | - | FIG013040 | PLF_590_ | PGF_0003 | NP_80412 | 117 | rcsF | Protein RcsF | |
| fig\|209261 | IgaA | Salmonella enterica subsp. enterica serovar Typhi str. Ty | 209261,6 | NC_004631 | t4011 | VBISalEnt | PATRIC.20 | PATRIC | CDS | 4162923 | 4165055 | 2133 | - | FIG000040 | PLF_590_ | PGF_0001 | NP_80762 | 710 | yrfF | IgaA: a membrane protein that prevents overactivation of the Rcs regulatory system | |
| fig\|209261 | RcsC | Salmonella enterica subsp. enterica serovar Typhi str. Ty | 209261,6 | NC_004631 | t0594 | VBISalEnt | PATRIC.20 | PATRIC | CDS | 674170 | 677016 | 2847 | + | FIG000042 | PLF_590_ | PGF_0005 | NP_80445 | 948 | rcsC | Sensor his | GO:0004673\|protein histidine kinase activity |
| fig\|209261 | RcsB | Salmonella enterica subsp. enterica serovar Typhi str. Ty | 209261,6 | NC_004631 | t0595 | VBISalEnt | PATRIC.20 | PATRIC | CDS | 677119 | 677769 | 651 | - | FIG000041 | PLF_590_ | PGF_0042 | NP_80445 | 216 | rcsB | DNA-binding capsular synthesis response regulator RcsB | |
| fig\|209261 | RcsD | Salmonella enterica subsp. enterica serovar Typhi str. Ty | 209261,6 | NC_004631 | t0596 | VBISalEnt | PATRIC.20 | PATRIC | CDS | 677786 | 680455 | 2670 | - | FIG000045 | PLF_590_ | PGF_0003 | NP_80445 | 889 | yojN | Phosphotransferase RcsD | |
| fig\|588858 | DnaK | Salmonella enterica subsp. enterica serovar Typhimurium | 588858.6 | CP001363 | STM14_00 | VBISalEnt | PATRIC.58 | PATRIC | CDS | 11593 | 13509 | 1917 | + | FIG000233 | PLF_590_ | PGF_1035 | ACY86550 | 638 | dnaK | Chaperone protein DnaK | |
| fig\|588858 | RcsD | Salmonella enterica subsp. enterica serovar Typhimurium | 588858,6 | CP001363 | STM14_28 | VBISalEnt | PATRIC.58 | PATRIC | CDS | 2418861 | 2421530 | 2670 | + | FIG000045 | PLF_590_ | PGF_0003 | ACY89240 | 889 | yojN | Phosphotransferase RcsD | |
| fig\|588858 | RcsB | Salmonella enterica subsp. enterica serovar Typhimurium | 588858,6 | CP001363 | STM14_28 | VBISalEnt | PATRIC.58 | PATRIC | CDS | 2421547 | 2422197 | 651 | + | FIG000041 | PLF_590_ | PGF_0042 | ACY89241 | 216 | rcsB | DNA-binding capsular synthesis response regulator RcsB | |
| fig\|588858 | RcsC | Salmonella enterica subsp. enterica serovar Typhimurium | 588858,6 | CP001363 | STM14_28 | VBISalEnt | PATRIC.58 | PATRIC | CDS | 2422300 | 2425146 | 2847 | - | FIG000042 | PLF_590_ | PGF_0005 | ACY89242 | 948 | rcsC | Sensor his | GO:0004673\|protein histidine kinase activity |
| fig\|588858 | IgaA | Salmonella enterica subsp. enterica serovar Typhimurium | 588858,6 | CP001363 | STM14_42 | VBISalEnt | PATRIC.58 | PATRIC | CDS | 3664082 | 3666214 | 2133 | + | FIG000040 | PLF_590_ | PGF_0001 | ACY90598 | 710 | yrfF | IgaA: a membrane protein that prevents overactivation of the Rcs regulatory system | |
| fig\|588858 | RcsF | Salmonella enterica subsp. enterica serovar Typhimurium | 588858,6 | CP001363 | STM14_02 | VBISalEnt | PATRIC.58 | PATRIC | CDS | 285700 | 285831 | 132 | - | FIG013040 | PLF_590_ | PGF_0003 | ACY86819 | 43 | rcsF | Protein RcsF | |
| fig\|216597 | DnaK | Salmonella enterica subsp. enterica serovar Typhimurium | 216597.6 | FQ312003 | SL1344_0 | VBISalEnt | PATRIC.21 | PATRIC | CDS | 11593 | 13509 | 1917 | + | FIG000233 | PLF_590_ | PGF_1035 | CBW1611 | 638 | dnaK | Chaperone protein DnaK | |
| fig\|216597 | RcsD | Salmonella enterica subsp. enterica serovar Typhimurium | 216597,6 | FQ312003 | SL1344_2 | VBISalEnt | PATRIC.21 | PATRIC | CDS | 2365024 | 2367693 | 2670 | + | FIG000045 | PLF_590_ | PGF_0003 | CBW1834 | 889 | yojN | Phosphotransferase RcsD | |
| fig\|216597 | RcsB | Salmonella enterica subsp. enterica serovar Typhimurium | 216597,6 | FQ312003 | SL1344_2 | VBISalEnt | PATRIC.21 | PATRIC | CDS | 2367710 | 2368360 | 651 | + | FIG000041 | PLF_590_ | PGF_0042 | CBW18341 | 216 | rcsB | DNA-binding capsular synthesis response regulator RcsB | |
| fig\|216597 | RcsC | Salmonella enterica subsp. enterica serovar Typhimurium | 216597,6 | FQ312003 | SL1344_2 | VBISalEnt | PATRIC.21 | PATRIC | CDS | 2368463 | 2371309 | 2847 | - | FIG000042 | PLF_590_ | PGF_0005 | CBW1834 | 948 | rcsC | Sensor his | GO:0004673\|protein histidine kinase activity |
| fig\|216597 | RcsF | Salmonella enterica subsp. enterica serovar Typhimurium | 216597,6 | FQ312003 | SL1344_0 | VBISalEnt | PATRIC.21 | PATRIC | CDS | 284991 | 285122 | 132 | - | FIG013040 | PLF_590_ | PGF_0003 | CBW1634 | 43 | rcsF | Protein RcsF | |
| fig\|216597 | IgaA | Salmonella enterica subsp. enterica serovar Typhimurium | 216597,6 | FQ312003 | SL1344_3 | VBISalEnt | PATRIC.21 | PATRIC | CDS | 3671706 | 3673838 | 2133 | + | FIG000040 | PLF_590_ | PGF_0001 | CBW1955 | 710 | yrfF | IgaA: a membrane protein that prevents overactivation of the Rcs regulatory system | |
| fig\|144193 | DnaK | Serratia fonticola RB-25 | 1441930.4 | CP007044 | Z042_170 | VBISerFon | PATRIC.14 | PATRIC | CDS | 3780875 | 3782788 | 1914 | + | FIG00023369 | | PGF_1035 | AHG2112 | 637 | dnaK | Chaperone protein DnaK | |
| fig\|144193 | RcsF | Serratia fonticola RB-25 | 1441930,4 | CP007044 | Z042_058 | VBISerFon | PATRIC.14 | PATRIC | CDS | 1295717 | 1296001 | 285 | + | FIG01304060 | | PGF_0003 | AHG1918 | 94 | rcsF | Protein RcsF | |
| fig\|144193 | IgaA | Serratia fonticola RB-25 | 1441930,4 | CP007044 | Z042_102 | VBISerFon | PATRIC.14 | PATRIC | CDS | 2241213 | 2243348 | 2136 | + | FIG00004077 | | PGF_0001 | AHG1997 | 711 | | IgaA: a membrane protein that prevents overactivation of the Rcs regulatory system | |
| fig\|144193 | RcsD | Serratia fonticola RB-25 | 1441930,4 | CP007044 | Z042_031 | VBISerFon | PATRIC.14 | PATRIC | CDS | 696266 | 698965 | 2700 | + | FIG00004592 | | PGF_0003 | AHG1871 | 899 | | Phosphotransferase RcsD | |
| fig\|144193 | RcsB | Serratia fonticola RB-25 | 1441930,4 | CP007044 | Z042_031 | VBISerFon | PATRIC.14 | PATRIC | CDS | 698958 | 699608 | 651 | + | FIG00004153 | | PGF_0042 | AHG1871 | 216 | | DNA-binding capsular synthesis response regulator RcsB | |
| fig\|144193 | RcsC | Serratia fonticola RB-25 | 1441930,4 | CP007044 | Z042_031 | VBISerFon | PATRIC.14 | PATRIC | CDS | 699659 | 702538 | 2880 | - | FIG00004208 | | PGF_0005 | AHG1871 | 959 | | Sensor his | GO:0004673\|protein histidine kinase activity |
| fig\|134661 | DnaK | Serratia liquefaciens ATCC 27592 | 1346614.3 | CP006252 | M495_028 | VBISerLiq | PATRIC.13 | PATRIC | CDS | 601690 | 603603 | 1914 | + | FIG000233 | PLF_613_ | PGF_1035 | AGQ2940 | 637 | dnaK | Chaperone protein DnaK | |
| fig\|134661 | RcsD | Serratia liquefaciens ATCC 27592 | 1346614,3 | CP006252 | M495_167 | VBISerLiq | PATRIC.13 | PATRIC | CDS | 3483825 | 3486527 | 2703 | + | FIG000045 | PLF_613_ | PGF_0003 | AGQ3204 | 900 | | Phosphotransferase RcsD | |
| fig\|134661 | RcsB | Serratia liquefaciens ATCC 27592 | 1346614,3 | CP006252 | M495_167 | VBISerLiq | PATRIC.13 | PATRIC | CDS | 3486520 | 3487170 | 651 | + | FIG000041 | PLF_613_ | PGF_0042 | AGQ3204 | 216 | | DNA-binding capsular synthesis response regulator RcsB | |
| fig\|134661 | RcsC | Serratia liquefaciens ATCC 27592 | 1346614,3 | CP006252 | M495_167 | VBISerLiq | PATRIC.13 | PATRIC | CDS | 3487221 | 3490091 | 2871 | - | FIG000042 | PLF_613_ | PGF_0005 | AGQ3205 | 956 | | Sensor his | GO:0004673\|protein histidine kinase activity |
| fig\|134661 | RcsF | Serratia liquefaciens ATCC 27592 | 1346614,3 | CP006252 | M495_195 | VBISerLiq | PATRIC.13 | PATRIC | CDS | 4086874 | 4087278 | 405 | + | FIG013040 | PLF_613_ | PGF_0003 | AGQ3255 | 134 | rcsF | Protein RcsF | |
| fig\|134661 | IgaA | Serratia liquefaciens ATCC 27592 | 1346614,3 | CP006252 | M495_232 | VBISerLiq | PATRIC.13 | PATRIC | CDS | 4883408 | 4885528 | 2121 | + | FIG000040 | PLF_613_ | PGF_0001 | AGQ33247 | 706 | | IgaA: a membrane protein that prevents overactivation of the Rcs regulatory system | |
| fig\|273526 | DnaK | Serratia marcescens subsp. marcescens Db11 | 273526.11 | HG326223 | SMDB11_0010 | | PATRIC.27 | PATRIC | CDS | 11869 | 13782 | 1914 | + | FIG000233 | PLF_613_ | PGF_1035 | CDG1060 | 637 | dnaK | Chaperone protein DnaK | |
| fig\|273526 | RcsD | Serratia marcescens subsp. marcescens Db11 | 273526,11 | HG326223 | SMDB11_2662 | | PATRIC.27 | PATRIC | CDS | 2811670 | 2814372 | 2703 | + | FIG000045 | PLF_613_ | PGF_0003 | CDG13234 | 900 | rcsD | Phosphotransferase RcsD | |
| fig\|273526 | RcsB | Serratia marcescens subsp. marcescens Db11 | 273526,11 | HG326223 | SMDB11_2663 | | PATRIC.27 | PATRIC | CDS | 2814365 | 2815015 | 651 | + | FIG000041 | PLF_613_ | PGF_0042 | CDG1323 | 216 | rcsB | DNA-binding capsular synthesis response regulator RcsB | |
| fig\|273526 | RcsC | Serratia marcescens subsp. marcescens Db11 | 273526,11 | HG326223 | SMDB11_2664 | | PATRIC.27 | PATRIC | CDS | 2815077 | 2817938 | 2862 | - | FIG000042 | PLF_613_ | PGF_0005 | CDG1323 | 953 | rcsC | Sensor his | GO:0004673\|protein histidine kinase activity |
| fig\|273526 | RcsF | Serratia marcescens subsp. marcescens Db11 | 273526,11 | HG326223 | SMDB11_3145 | | PATRIC.27 | PATRIC | CDS | 3322917 | 3323147 | 231 | + | FIG013040 | PLF_613_ | PGF_0003 | CDG1370 | 76 | rcsF | Protein RcsF | |
| fig\|273526 | IgaA | Serratia marcescens subsp. marcescens Db11 | 273526,11 | HG326223 | SMDB11_3859 | | PATRIC.27 | PATRIC | CDS | 4099182 | 4101305 | 2124 | + | FIG000040 | PLF_613_ | PGF_0001 | CDG1442 | 707 | igaA | IgaA: a membrane protein that prevents overactivation of the Rcs regulatory system | |
| fig\|667129 | DnaK | Serratia odorifera DSM 4582 | 667129.3 | NZ_GG753567 | HMPREF0 | VBISerOd | PATRIC.66 | PATRIC | CDS | 590260 | 592173 | 1914 | - | FIG000233 | PLF_613_ | PGF_1035 | ZP_06637 | 637 | dnaK | Chaperone protein DnaK | |
| fig\|667129 | RcsF | Serratia odorifera DSM 4582 | 667129,3 | NZ_GG753567 | HMPREF0 | VBISerOd | PATRIC.66 | PATRIC | CDS | 2481169 | 2481522 | 354 | - | FIG013040 | PLF_613_ | PGF_0003 | ZP_06639 | 117 | rcsF | Protein RcsF | |
| fig\|667129 | RcsC | Serratia odorifera DSM 4582 | 667129,3 | NZ_GG753567 | HMPREF0 | VBISerOd | PATRIC.66 | PATRIC | CDS | 2965128 | 2967989 | 2862 | + | FIG000042 | PLF_613_ | PGF_0005 | ZP_06639 | 953 | rcsC | Sensor his | GO:0004673\|protein histidine kinase activity |
| fig\|667129 | RcsB | Serratia odorifera DSM 4582 | 667129,3 | NZ_GG753567 | HMPREF0 | VBISerOd | PATRIC.66 | PATRIC | CDS | 2968069 | 2968719 | 651 | - | FIG000041 | PLF_613_ | PGF_0042 | ZP_06639 | 216 | | DNA-binding capsular synthesis response regulator RcsB | |
| fig\|667129 | RcsD | Serratia odorifera DSM 4582 | 667129,3 | NZ_GG753567 | HMPREF0 | VBISerOd | PATRIC.66 | PATRIC | CDS | 2968712 | 2971417 | 2706 | - | FIG000045 | PLF_613_ | PGF_0003 | ZP_06639 | 901 | | Phosphotransferase RcsD | |
| fig\|667129 | IgaA | Serratia odorifera DSM 4582 | 667129,3 | NZ_GG753567 | HMPREF0 | VBISerOd | PATRIC.66 | PATRIC | CDS | 1045415 | 1047535 | 2121 | + | FIG000040 | PLF_613_ | PGF_0001 | ZP_06637 | 706 | igaA | IgaA: a membrane protein that prevents overactivation of the Rcs regulatory system | |
| fig\|768492 | DnaK | Serratia plymuthica AS9 | 768492.3 | NC_015567 | SerAS9_0 | VBISerSp1 | PATRIC.76 | PATRIC | CDS | 702184 | 704097 | 1914 | + | FIG000233 | PLF_613_ | PGF_1035 | YP_00450 | 637 | | Chaperone protein DnaK | |
| fig\|768492 | RcsD | Serratia plymuthica AS9 | 768492,3 | NC_015567 | SerAS9_3 | VBISerSp1 | PATRIC.76 | PATRIC | CDS | 3679859 | 3682561 | 2703 | + | FIG000045 | PLF_613_ | PGF_0003 | YP_00450 | 900 | | Phosphotransferase RcsD | |
| fig\|768492 | RcsB | Serratia plymuthica AS9 | 768492,3 | NC_015567 | SerAS9_3 | VBISerSp1 | PATRIC.76 | PATRIC | CDS | 3682554 | 3683204 | 651 | + | FIG000041 | PLF_613_ | PGF_0042 | YP_00450 | 216 | | DNA-binding capsular synthesis response regulator RcsB | |
| fig\|768492 | RcsC | Serratia plymuthica AS9 | 768492,3 | NC_015567 | SerAS9_3 | VBISerSp1 | PATRIC.76 | PATRIC | CDS | 3683258 | 3686128 | 2871 | - | FIG000042 | PLF_613_ | PGF_0005 | YP_00450 | 956 | | Sensor his | GO:0004673\|protein histidine kinase activity |
| fig\|768492 | RcsF | Serratia plymuthica AS9 | 768492,3 | NC_015567 | SerAS9_3 | VBISerSp1 | PATRIC.76 | PATRIC | CDS | 4221839 | 4222069 | 231 | + | FIG013040 | PLF_613_ | PGF_0003 | YP_00450 | 76 | | Protein RcsF | |
| fig\|768492 | IgaA | Serratia plymuthica AS9 | 768492,3 | NC_015567 | SerAS9_4 | VBISerSp1 | PATRIC.76 | PATRIC | CDS | 5072688 | 5074823 | 2136 | + | FIG000040 | PLF_613_ | PGF_0001 | YP_00450 | 711 | | IgaA: a membrane protein that prevents overactivation of the Rcs regulatory system | |
| fig\|399741 | DnaK | Serratia proteamaculans 568 | 399741.7 | NC_009832 | Spro_0692 | VBISerPro | PATRIC.39 | PATRIC | CDS | 762495 | 764408 | 1914 | + | FIG000233 | PLF_613_ | PGF_1035 | YP_00147 | 637 | dnaK | Chaperone protein DnaK | |
| fig\|399741 | RcsD | Serratia proteamaculans 568 | 399741,7 | NC_009832 | Spro_3266 | VBISerPro | PATRIC.39 | PATRIC | CDS | 3602311 | 3605013 | 2703 | + | FIG000045 | PLF_613_ | PGF_0003 | YP_00147 | 900 | | Phosphotransferase RcsD | |
| fig\|399741 | RcsB | Serratia proteamaculans 568 | 399741,7 | NC_009832 | Spro_3267 | VBISerPro | PATRIC.39 | PATRIC | CDS | 3605006 | 3605656 | 651 | + | FIG000041 | PLF_613_ | PGF_0042 | YP_00147 | 216 | | DNA-binding capsular synthesis response regulator RcsB | |
| fig\|399741 | RcsC | Serratia proteamaculans 568 | 399741,7 | NC_009832 | Spro_3268 | VBISerPro | PATRIC.39 | PATRIC | CDS | 3605708 | 3608581 | 2874 | - | FIG000042 | PLF_613_ | PGF_0005 | YP_00147 | 957 | | Sensor his | GO:0004673\|protein histidine kinase activity |
| fig\|399741 | RcsF | Serratia proteamaculans 568 | 399741,7 | NC_009832 | Spro_3756 | VBISerPro | PATRIC.39 | PATRIC | CDS | 4153209 | 4153439 | 231 | + | FIG013040 | PLF_613_ | PGF_0003 | YP_00147 | 76 | rcsF | Protein RcsF | |

Table-S2

| | | | | | | | | | | | | | | | | | | | | | |
|---|---|---|---|---|---|---|---|---|---|---|---|---|---|---|---|---|---|---|---|---|---|
| fig\|399741 | IgaA | Serratia proteamaculans 568 | 399741,7 | NC_009832 | Spro_4613 | VBISerPro | PATRIC.39 | PATRIC | CDS | 5088503 | 5090635 | 2133 | + | FIG000040 | PLF_613_ | PGF_0001 | YP_00148 | 710 | | IgaA: a membrane protein that prevents overactivation of the Rcs regulatory system | |
| fig\|300268 | DnaK | Shigella boydii Sb227 | 300268.11 | NC_007613 | SBO_0015 | VBIShiBoy | PATRIC.30 | PATRIC | CDS | 13301 | 15217 | 1917 | + | FIG000233 | PLF_620_ | PGF_1035 | YP_40657 | 638 | dnaK | Chaperone protein DnaK | |
| fig\|300268 | RcsC | Shigella boydii Sb227 | 300268,11 | NC_007613 | SBO_2089 | VBIShiBoy | PATRIC.30 | PATRIC | CDS | 2065775 | 2068576 | 2802 | + | FIG000042 | PLF_620_ | PGF_0005 | YP_40849 | 933 | rcsC | Sensor hist | GO:0004673\|protein histidine kinase activity |
| fig\|300268 | RcsB | Shigella boydii Sb227 | 300268,11 | NC_007613 | SBO_2090 | VBIShiBoy | PATRIC.30 | PATRIC | CDS | 2068776 | 2069426 | 651 | - | FIG000041 | PLF_620_ | PGF_0042 | YP_40849 | 216 | rcsB | DNA-binding capsular synthesis response regulator RcsB | |
| fig\|300268 | RcsD | Shigella boydii Sb227 | 300268,11 | NC_007613 | SBO_2091 | VBIShiBoy | PATRIC.30 | PATRIC | CDS | 2069443 | 2072115 | 2673 | - | FIG000045 | PLF_620_ | PGF_0003 | YP_40849 | 890 | yojN | Phosphotransferase RcsD | |
| fig\|300268 | RcsF | Shigella boydii Sb227 | 300268,11 | NC_007613 | SBO_0185 | VBIShiBoy | PATRIC.30 | PATRIC | CDS | 205569 | 205922 | 354 | - | FIG013040 | PLF_620_ | PGF_0003 | YP_40674 | 117 | rcsF | Protein RcsF | |
| fig\|300268 | IgaA | Shigella boydii Sb227 | 300268,11 | NC_007613 | SBO_3385 | VBIShiBoy | PATRIC.30 | PATRIC | CDS | 3368493 | 3370628 | 2136 | + | FIG000040 | PLF_620_ | PGF_0001 | YP_40970 | 711 | yrfF | IgaA: a membrane protein that prevents overactivation of the Rcs regulatory system | |
| fig\|300267 | DnaK | Shigella dysenteriae Sd197 | 300267.13 | NC_007606 | SDY_0013 | VBIShiDys | PATRIC.30 | PATRIC | CDS | 12155 | 14071 | 1917 | + | FIG000233 | PLF_620_ | PGF_1035 | YP_40174 | 638 | dnaK | Chaperone protein DnaK | |
| fig\|300267 | RcsF | Shigella dysenteriae Sd197 | 300267,13 | NC_007606 | SDY_0215 | VBIShiDys | PATRIC.30 | PATRIC | CDS | 220562 | 220915 | 354 | - | FIG013040 | PLF_620_ | PGF_0003 | YP_40193 | 117 | rcsF | Protein RcsF | |
| fig\|300267 | IgaA | Shigella dysenteriae Sd197 | 300267,13 | NC_007606 | SDY_3680 | VBIShiDys | PATRIC.30 | PATRIC | CDS | 3411557 | 3413692 | 2136 | - | FIG000040 | PLF_620_ | PGF_0001 | YP_40513 | 711 | yrfF | IgaA: a membrane protein that prevents overactivation of the Rcs regulatory system | |
| fig\|300267 | RcsC | Shigella dysenteriae Sd197 | 300267,13 | NC_007606 | SDY_0856 | VBIShiDys | PATRIC.30 | PATRIC | CDS | 802405 | 805206 | 2802 | + | FIG000042 | PLF_620_ | PGF_0005 | YP_40252 | 933 | rcsC | Sensor hist | GO:0004673\|protein histidine kinase activity |
| fig\|300267 | RcsB | Shigella dysenteriae Sd197 | 300267,13 | NC_007606 | SDY_0857 | VBIShiDys | PATRIC.30 | PATRIC | CDS | 805406 | 806056 | 651 | - | FIG000041 | PLF_620_ | PGF_0042 | YP_40252 | 216 | rcsB | DNA-binding capsular synthesis response regulator RcsB | |
| fig\|300267 | RcsD | Shigella dysenteriae Sd197 | 300267,13 | NC_007606 | SDY_0858 | VBIShiDys | PATRIC.30 | PATRIC | CDS | 806073 | 808661 | 2589 | - | FIG000045 | PLF_620_ | PGF_0003 | YP_40252 | 862 | yojN | Phosphotransferase RcsD | |
| fig\|108603 | RcsF | Shigella flexneri 5a str. M90T | 1086030.18 | CP037923 | EKN05_15770 | | PATRIC.10 | PATRIC | CDS | 3009299 | 3009703 | 405 | + | | PLF_620_ | PGF_00037945 | | 134 | rcsF | Protein RcsF | |
| fig\|108603 | DnaK | Shigella flexneri 5a str. M90T | 1086030.18 | CP037923 | EKN05_16635 | | PATRIC.10 | PATRIC | CDS | 3205706 | 3207622 | 1917 | - | | PLF_620_ | PGF_10357457 | | 638 | dnaK | Chaperone protein DnaK | |
| fig\|108603 | IgaA | Shigella flexneri 5a str. M90T | 1086030.18 | CP037923 | EKN05_22025 | | PATRIC.10 | PATRIC | CDS | 4333643 | 4335778 | 2136 | - | | PLF_620_ | PGF_00013790 | | 711 | igaA | IgaA: a membrane protein that prevents overactivation of the Rcs regulatory system | |
| fig\|108603 | RcsC | Shigella flexneri 5a str. M90T | 1086030.18 | CP037923 | EKN05_04390 | | PATRIC.10 | PATRIC | CDS | 876167 | 879016 | 2850 | + | | PLF_620_ | PGF_00050577 | | 949 | rcsC | Sensor hist | GO:0004673\|protein histidine kinase activity |
| fig\|108603 | RcsB | Shigella flexneri 5a str. M90T | 1086030.18 | CP037923 | EKN05_04395 | | PATRIC.10 | PATRIC | CDS | 879216 | 879866 | 651 | - | | PLF_620_ | PGF_00421954 | | 216 | rcsB | DNA-binding capsular synthesis response regulator RcsB | |
| fig\|108603 | RcsD | Shigella flexneri 5a str. M90T | 1086030.18 | CP037923 | EKN05_04400 | | PATRIC.10 | PATRIC | CDS | 879883 | 882555 | 2673 | - | | PLF_620_ | PGF_00034034 | | 890 | rcsD | Phosphotransferase RcsD | |
| fig\|300269 | DnaK | Shigella sonnei Ss046 | 300269.12 | NC_007384 | SSON_001 | VBIShiSon | PATRIC.30 | PATRIC | CDS | 12939 | 14855 | 1917 | + | FIG000233 | PLF_620_ | PGF_1035 | YP_30904 | 638 | dnaK | Chaperone protein DnaK | |
| fig\|300269 | RcsF | Shigella sonnei Ss046 | 300269,12 | NC_007384 | SSON_021 | VBIShiSon | PATRIC.30 | PATRIC | CDS | 235903 | 236256 | 354 | - | FIG013040 | PLF_620_ | PGF_0003 | YP_30923 | 117 | rcsF | Protein RcsF | |
| fig\|300269 | RcsD | Shigella sonnei Ss046 | 300269,12 | NC_007384 | SSON_227 | VBIShiSon | PATRIC.30 | PATRIC | CDS | 2388560 | 2391133 | 2574 | + | FIG000045 | PLF_620_ | PGF_0003 | YP_31115 | 857 | yojN | Phosphotransferase RcsD | |
| fig\|300269 | RcsB | Shigella sonnei Ss046 | 300269,12 | NC_007384 | SSON_227 | VBIShiSon | PATRIC.30 | PATRIC | CDS | 2391150 | 2391800 | 651 | + | FIG000041 | PLF_620_ | PGF_0042 | YP_31115 | 216 | rcsB | DNA-binding capsular synthesis response regulator RcsB | |
| fig\|300269 | RcsC | Shigella sonnei Ss046 | 300269,12 | NC_007384 | SSON_227 | VBIShiSon | PATRIC.30 | PATRIC | CDS | 2392000 | 2394849 | 2850 | - | FIG000042 | PLF_620_ | PGF_0005 | YP_31115 | 949 | rcsC | Sensor hist | GO:0004673\|protein histidine kinase activity |
| fig\|300269 | IgaA | Shigella sonnei Ss046 | 300269,12 | NC_007384 | SSON_352 | VBIShiSon | PATRIC.30 | PATRIC | CDS | 3689115 | 3691250 | 2136 | + | FIG000040 | PLF_620_ | PGF_0001 | YP_31232 | 711 | yrfF | IgaA: a membrane protein that prevents overactivation of the Rcs regulatory system | |
| fig\|630626 | DnaK | Shimwellia blattae DSM 4481 = NBRC 105725 | 630626.3 | CP001560 | EBL_c333 | VBIEscBla | PATRIC.63 | PATRIC | CDS | 3448110 | 3450026 | 1917 | - | FIG00023369 | | PGF_1035 | AFJ48401. | 638 | dnaK | Chaperone protein DnaK | |
| fig\|630626 | RcsC | Shimwellia blattae DSM 4481 = NBRC 105725 | 630626,3 | CP001560 | EBL_c130 | VBIEscBla | PATRIC.63 | PATRIC | CDS | 1353215 | 1356064 | 2850 | + | FIG00004208 | | PGF_0005 | AFJ46410. | 949 | rcsC | Sensor hist | GO:0004673\|protein histidine kinase activity |
| fig\|630626 | RcsB | Shimwellia blattae DSM 4481 = NBRC 105725 | 630626,3 | CP001560 | EBL_c130 | VBIEscBla | PATRIC.63 | PATRIC | CDS | 1356100 | 1356750 | 651 | - | FIG00004153 | | PGF_0042 | AFJ46411. | 216 | rcsB | DNA-binding capsular synthesis response regulator RcsB | |
| fig\|630626 | RcsD | Shimwellia blattae DSM 4481 = NBRC 105725 | 630626,3 | CP001560 | EBL_c130 | VBIEscBla | PATRIC.63 | PATRIC | CDS | 1356767 | 1359430 | 2664 | - | FIG00004592 | | PGF_0003 | AFJ46412. | 887 | yojN | Phosphotransferase RcsD | |
| fig\|630626 | IgaA | Shimwellia blattae DSM 4481 = NBRC 105725 | 630626,3 | CP001560 | EBL_c023 | VBIEscBla | PATRIC.63 | PATRIC | CDS | 262307 | 264430 | 2124 | - | FIG00004077 | | PGF_0001 | AFJ45366. | 707 | yrfF | IgaA: a membrane protein that prevents overactivation of the Rcs regulatory system | |
| fig\|630626 | RcsF | Shimwellia blattae DSM 4481 = NBRC 105725 | 630626,3 | CP001560 | EBL_c316 | VBIEscBla | PATRIC.63 | PATRIC | CDS | 3251148 | 3251558 | 411 | + | FIG01304060 | | PGF_0003 | AFJ48229. | 136 | rcsF | Protein RcsF | |
| fig\|343509 | DnaK | Sodalis glossinidius str. 'morsitans' | 343509.12 | NC_007712 | SG0409 | VBISodGlo | PATRIC.34 | PATRIC | CDS | 726305 | 728215 | 1911 | + | FIG000233 | PLF_8456 | PGF_1035 | YP_45408 | 636 | dnaK | Chaperone protein DnaK | |
| fig\|343509 | RcsD | Sodalis glossinidius str. 'morsitans' | 343509,12 | NC_007712 | SG1579 | VBISodGlo | PATRIC.34 | PATRIC | CDS | 2708478 | 2711171 | 2694 | + | FIG000045 | PLF_8456 | PGF_0003 | YP_45525 | 897 | | Phosphotransferase RcsD | |
| fig\|343509 | RcsB | Sodalis glossinidius str. 'morsitans' | 343509,12 | NC_007712 | SG1580 | VBISodGlo | PATRIC.34 | PATRIC | CDS | 2711180 | 2711830 | 651 | + | FIG000041 | PLF_8456 | PGF_0042 | YP_45526 | 216 | | DNA-binding capsular synthesis response regulator RcsB | |
| fig\|343509 | RcsC | Sodalis glossinidius str. 'morsitans' | 343509,12 | NC_007712 | SG1581 | VBISodGlo | PATRIC.34 | PATRIC | CDS | 2711872 | 2714802 | 2931 | - | FIG000042 | PLF_8456 | PGF_0005 | YP_45526 | 976 | | Sensor hist | GO:0004673\|protein histidine kinase activity |
| fig\|343509 | RcsF | Sodalis glossinidius str. 'morsitans' | 343509,12 | NC_007712 | SG1918 | VBISodGlo | PATRIC.34 | PATRIC | CDS | 3315048 | 3315341 | 294 | + | FIG013040 | PLF_8456 | PGF_0003 | YP_45559 | 97 | rcsF | Protein RcsF | |
| fig\|343509 | IgaA | Sodalis glossinidius str. 'morsitans' | 343509,12 | NC_007712 | SG2313 | VBISodGlo | PATRIC.34 | PATRIC | CDS | 3975150 | 3977318 | 2169 | + | FIG000040 | PLF_8456 | PGF_0001 | YP_45599 | 722 | | IgaA: a membrane protein that prevents overactivation of the Rcs regulatory system | |
| fig\|642227 | DnaK | Tatumella morbirosei LMG 23360 | 642227.3 | JPKR01000011 | HA49_19370 | | PATRIC.64 | PATRIC | CDS | 161293 | 163206 | 1914 | - | FIG000233 | PLF_8298 | PGF_1035 | KGD70597 | 637 | | Chaperone protein DnaK | |
| fig\|642227 | RcsD | Tatumella morbirosei LMG 23360 | 642227,3 | JPKR01000009 | HA49_16120 | | PATRIC.64 | PATRIC | CDS | 543198 | 545423 | 2226 | + | FIG000045 | PLF_8298 | PGF_00034034 | | 741 | | Phosphotransferase RcsD | |
| fig\|642227 | RcsB | Tatumella morbirosei LMG 23360 | 642227,3 | JPKR01000009 | HA49_16125 | | PATRIC.64 | PATRIC | CDS | 545438 | 546085 | 648 | + | FIG000041 | PLF_8298 | PGF_0042 | KGD7227 | 215 | | DNA-binding capsular synthesis response regulator RcsB | |
| fig\|642227 | RcsC | Tatumella morbirosei LMG 23360 | 642227,3 | JPKR01000009 | HA49_16130 | | PATRIC.64 | PATRIC | CDS | 546135 | 548981 | 2847 | - | FIG000042 | PLF_8298 | PGF_0005 | KGD72271 | 948 | | Sensor hist | GO:0004673\|protein histidine kinase activity |
| fig\|642227 | RcsF | Tatumella morbirosei LMG 23360 | 642227,3 | JPKR01000011 | HA49_18705 | | PATRIC.64 | PATRIC | CDS | 3919 | 4323 | 405 | + | FIG013040 | PLF_8298 | PGF_0003 | KGD70471 | 134 | | Protein RcsF | |
| fig\|642227 | IgaA | Tatumella morbirosei LMG 23360 | 642227,3 | JPKR01000003 | HA49_02800 | | PATRIC.64 | PATRIC | CDS | 219126 | 221258 | 2133 | - | FIG000040 | PLF_8298 | PGF_0001 | KGD79526 | 710 | | IgaA: a membrane protein that prevents overactivation of the Rcs regulatory system | |
| fig\|100599 | DnaK | Tatumella ptyseos ATCC 33301 | 1005995.4 | JMPR01000020 | GTPT_1131 | | PATRIC.10 | PATRIC | CDS | 32393 | 34300 | 1908 | + | FIG000233 | PLF_8298 | PGF_10357457 | | 635 | dnaK | Chaperone protein DnaK | |
| fig\|100599 | RcsF | Tatumella ptyseos ATCC 33301 | 1005995,4 | JMPR01000020 | GTPT_1261 | | PATRIC.10 | PATRIC | CDS | 186612 | 187013 | 402 | - | FIG013040 | PLF_8298 | PGF_00037945 | | 133 | rcsF | Protein RcsF | |
| fig\|100599 | RcsC | Tatumella ptyseos ATCC 33301 | 1005995,4 | JMPR01000028 | GTPT_1728 | | PATRIC.10 | PATRIC | CDS | 128217 | 131060 | 2844 | + | FIG000042 | PLF_8298 | PGF_00050577 | | 947 | rcsC | Sensor hist | GO:0004673\|protein histidine kinase activity |
| fig\|100599 | RcsB | Tatumella ptyseos ATCC 33301 | 1005995,4 | JMPR01000028 | GTPT_1729 | | PATRIC.10 | PATRIC | CDS | 131193 | 131840 | 648 | - | FIG000041 | PLF_8298 | PGF_00421954 | | 215 | rcsB | DNA-binding capsular synthesis response regulator RcsB | |
| fig\|100599 | RcsD | Tatumella ptyseos ATCC 33301 | 1005995,4 | JMPR01000028 | GTPT_1730 | | PATRIC.10 | PATRIC | CDS | 131845 | 134493 | 2649 | - | FIG000045 | PLF_8298 | PGF_00034034 | | 882 | rcsD | Phosphotransferase RcsD | |
| fig\|100599 | IgaA | Tatumella ptyseos ATCC 33301 | 1005995,4 | JMPR01000008 | GTPT_0423 | | PATRIC.10 | PATRIC | CDS | 36981 | 39005 | 2025 | + | FIG000040 | PLF_8298 | PGF_00013790 | | 674 | yrfF | IgaA: a membrane protein that prevents overactivation of the Rcs regulatory system | |
| fig\|379893 | DnaK | Trabulsiella odontotermitis strain TbO2.3 | 379893.5 | LIFV01000002 | | | PATRIC.37 | PATRIC | CDS | 195070 | 196986 | 1917 | - | FIG000233 | PLF_1588 | PGF_10357457 | | 638 | | Chaperone protein DnaK | |
| fig\|379893 | RcsF | Trabulsiella odontotermitis strain TbO2.3 | 379893,5 | LIFV01000002 | | | PATRIC.37 | PATRIC | CDS | 3940 | 4344 | 405 | + | FIG013040 | PLF_1588 | PGF_00037945 | | 134 | | Protein RcsF | |
| fig\|379893 | RcsD | Trabulsiella odontotermitis strain TbO2.3 | 379893,5 | LIFV01000005 | | | PATRIC.37 | PATRIC | CDS | 210746 | 213424 | 2679 | + | FIG000045 | PLF_1588 | PGF_00034034 | | 892 | | Phosphotransferase RcsD | |
| fig\|379893 | RcsB | Trabulsiella odontotermitis strain TbO2.3 | 379893,5 | LIFV01000005 | | | PATRIC.37 | PATRIC | CDS | 213441 | 214091 | 651 | + | FIG000041 | PLF_1588 | PGF_00421954 | | 216 | | DNA-binding capsular synthesis response regulator RcsB | |
| fig\|379893 | RcsC | Trabulsiella odontotermitis strain TbO2.3 | 379893,5 | LIFV01000005 | | | PATRIC.37 | PATRIC | CDS | 214143 | 217091 | 2949 | - | FIG000042 | PLF_1588 | PGF_00050577 | | 982 | | Sensor hist | GO:0004673\|protein histidine kinase activity |
| fig\|379893 | IgaA | Trabulsiella odontotermitis strain TbO2.3 | 379893,5 | LIFV01000001 | | | PATRIC.37 | PATRIC | CDS | 51545 | 53674 | 2130 | + | FIG000040 | PLF_1588 | PGF_00013790 | | 709 | | IgaA: a membrane protein that prevents overactivation of the Rcs regulatory system | |
| fig\|36870.6 | DnaK | Wigglesworthia glossinidia endosymbiont of Glossina brev | 36870.6 | NC_004344 | WGLp299 | VBIWigGlo | PATRIC.36 | PATRIC | CDS | 351424 | 353349 | 1926 | - | FIG00023369 | | PGF_1035 | NP_87130 | 641 | dnaK | Chaperone protein DnaK | |
| fig\|36868.4 | DnaK | Wigglesworthia glossinidia endosymbiont of Glossina mor | 36868.4 | NC_016893 | WIGMOR_ | VBIWigGlo | PATRIC.36 | PATRIC | CDS | 488041 | 489966 | 1926 | + | FIG00023369 | | PGF_1035 | YP_00526 | 641 | dnaK | Chaperone protein DnaK | |
| fig\|406818 | DnaK | Xenorhabdus bovienii SS-2004 | 406818.4 | NC_013892 | XBJ1_174 | VBIXenBo | PATRIC.40 | PATRIC | CDS | 1704793 | 1706697 | 1905 | - | FIG000233 | PLF_626_ | PGF_1035 | YP_00346 | 634 | dnaK | Chaperone protein DnaK | |
| fig\|406818 | RcsF | Xenorhabdus bovienii SS-2004 | 406818,4 | NC_013892 | XBJ1_061 | VBIXenBo | PATRIC.40 | PATRIC | CDS | 632911 | 633036 | 126 | - | | PLF_626_ | PGF_0391 | YP_00346 | 41 | rcsF | hypothetical protein | |
| fig\|406818 | IgaA | Xenorhabdus bovienii SS-2004 | 406818,4 | NC_013892 | XBJ1_006 | VBIXenBo | PATRIC.40 | PATRIC | CDS | 75528 | 77303 | 1776 | + | FIG000040 | PLF_626_ | PGF_0391 | YP_00346 | 591 | yrfF | hypothetical protein | |
| fig\|406818 | RcsC | Xenorhabdus bovienii SS-2004 | 406818,4 | NC_013892 | XBJ1_073 | VBIXenBo | PATRIC.40 | PATRIC | CDS | 718904 | 721669 | 2766 | + | FIG000042 | PLF_626_ | PGF_0005 | YP_00346 | 921 | rcsC | Sensor hist | GO:0004673\|protein histidine kinase activity |
| fig\|406818 | RcsB | Xenorhabdus bovienii SS-2004 | 406818,4 | NC_013892 | XBJ1_073 | VBIXenBo | PATRIC.40 | PATRIC | CDS | 721749 | 722399 | 651 | - | FIG000041 | PLF_626_ | PGF_0042 | YP_00346 | 216 | rcsB | DNA-binding capsular synthesis response regulator RcsB | |
| fig\|406818 | RcsD | Xenorhabdus bovienii SS-2004 | 406818,4 | NC_013892 | XBJ1_073 | VBIXenBo | PATRIC.40 | PATRIC | CDS | 722392 | 724944 | 2553 | - | FIG000045 | PLF_626_ | PGF_0003 | YP_00346 | 850 | rcsD | Phosphotransferase RcsD | |
| fig\|351671 | DnaK | Xenorhabdus doucetiae strain FRM16 = DSM 17909 | 351671.5 | FO704550 | XDD1_0760 | | PATRIC.35 | PATRIC | CDS | 725022 | 726932 | 1911 | - | FIG000233 | PLF_626_ | PGF_10357457 | | 636 | dnaK | Chaperone protein DnaK | |
| fig\|351671 | RcsC | Xenorhabdus doucetiae strain FRM16 = DSM 17909 | 351671,5 | FO704550 | XDD1_2560 | | PATRIC.35 | PATRIC | CDS | 2706968 | 2709739 | 2772 | + | FIG000042 | PLF_626_ | PGF_00050577 | | 923 | rcsC | Sensor hist | GO:0004673\|protein histidine kinase activity |
| fig\|351671 | RcsB | Xenorhabdus doucetiae strain FRM16 = DSM 17909 | 351671,5 | FO704550 | XDD1_2561 | | PATRIC.35 | PATRIC | CDS | 2709828 | 2710478 | 651 | - | FIG000041 | PLF_626_ | PGF_00421954 | | 216 | rcsB | DNA-binding capsular synthesis response regulator RcsB | |
| fig\|351671 | RcsD | Xenorhabdus doucetiae strain FRM16 = DSM 17909 | 351671,5 | FO704550 | XDD1_2562 | | PATRIC.35 | PATRIC | CDS | 2710480 | 2713224 | 2745 | - | FIG000045 | PLF_626_ | PGF_00034034 | | 914 | rcsD | Phosphotransferase RcsD | |
| fig\|351671 | RcsF | Xenorhabdus doucetiae strain FRM16 = DSM 17909 | 351671,5 | FO704550 | XDD1_3282 | | PATRIC.35 | PATRIC | CDS | 3475212 | 3475481 | 270 | + | FIG013040 | PLF_626_ | PGF_00037945 | | 89 | rcsF | Protein RcsF | |
| fig\|351671 | IgaA | Xenorhabdus doucetiae strain FRM16 = DSM 17909 | 351671,5 | FO704550 | XDD1_0078 | | PATRIC.35 | PATRIC | CDS | 90739 | 92841 | 2103 | + | FIG000040 | PLF_626_ | PGF_03915705 | | 700 | umoB | hypothetical protein | |
| fig\|406817 | DnaK | Xenorhabdus nematophila ATCC 19061 | 406817.27 | FN667742 | XNC1_0708 | | PATRIC.40 | PATRIC | CDS | 615061 | 616971 | 1911 | + | FIG000233 | PLF_626_ | PGF_10357457 | | 636 | dnaK | Chaperone protein DnaK | |
| fig\|406817 | RcsD | Xenorhabdus nematophila ATCC 19061 | 406817,27 | FN667742 | XNC1_2852 | | PATRIC.40 | PATRIC | CDS | 2859121 | 2861895 | 2775 | + | FIG000045 | PLF_626_ | PGF_00034034 | | 924 | rcsD | Phosphotransferase RcsD | |
| fig\|406817 | RcsB | Xenorhabdus nematophila ATCC 19061 | 406817,27 | FN667742 | XNC1_2853 | | PATRIC.40 | PATRIC | CDS | 2861897 | 2862547 | 651 | + | FIG000041 | PLF_626_ | PGF_00421954 | | 216 | rcsB | DNA-binding capsular synthesis response regulator RcsB | |
| fig\|406817 | RcsC | Xenorhabdus nematophila ATCC 19061 | 406817,27 | FN667742 | XNC1_2854 | | PATRIC.40 | PATRIC | CDS | 2862615 | 2865407 | 2793 | - | FIG000042 | PLF_626_ | PGF_00050577 | | 930 | rcsC | Sensor hist | GO:0004673\|protein histidine kinase activity |
| fig\|406817 | RcsF | Xenorhabdus nematophila ATCC 19061 | 406817,27 | FN667742 | XNC1_3983 | | PATRIC.40 | PATRIC | CDS | 3843055 | 3843450 | 396 | + | FIG013040 | PLF_626_ | PGF_03915707 | | 131 | rcsF | hypothetical protein | |
| fig\|406817 | IgaA | Xenorhabdus nematophila ATCC 19061 | 406817,27 | FN667742 | XNC1_4593 | | PATRIC.40 | PATRIC | CDS | 4387009 | 4389129 | 2121 | + | FIG000040 | PLF_626_ | PGF_03915705 | | 706 | yrfF | hypothetical protein | |
| fig\|142751 | DnaK | Xenorhabdus szentirmaii DSM 16338 | 1427518.3 | CBXF010000088 | XSR1_302 | VBIXenSz | PATRIC.14 | PATRIC | CDS | 223786 | 225696 | 1911 | + | FIG000233 | PLF_626_ | PGF_1035 | CDL83378 | 636 | dnaK | Chaperone protein DnaK | |
| fig\|142751 | RcsD | Xenorhabdus szentirmaii DSM 16338 | 1427518,3 | CBXF010000068 | XSR1_160 | VBIXenSz | PATRIC.14 | PATRIC | CDS | 24763 | 27492 | 2730 | + | FIG000045 | PLF_626_ | PGF_0003 | CDL81803 | 909 | rcsD | Phosphotransferase RcsD | |
| fig\|142751 | RcsB | Xenorhabdus szentirmaii DSM 16338 | 1427518,3 | CBXF010000068 | XSR1_160 | VBIXenSz | PATRIC.14 | PATRIC | CDS | 27494 | 28138 | 645 | + | FIG000041 | PLF_626_ | PGF_0042 | CDL81804 | 214 | rcsB | DNA-binding capsular synthesis response regulator RcsB | |
| fig\|142751 | RcsC | Xenorhabdus szentirmaii DSM 16338 | 1427518,3 | CBXF010000068 | XSR1_160 | VBIXenSz | PATRIC.14 | PATRIC | CDS | 28224 | 30986 | 2763 | - | FIG000042 | PLF_626_ | PGF_0005 | CDL81805 | 920 | rcsC | Sensor hist | GO:0004673\|protein histidine kinase activity |
| fig\|142751 | RcsF | Xenorhabdus szentirmaii DSM 16338 | 1427518,3 | CBXF010000077 | XSR1_200 | VBIXenSz | PATRIC.14 | PATRIC | CDS | 3699 | 4049 | 351 | + | FIG013040 | PLF_626_ | PGF_0391 | CDL82171 | 116 | rcsF | hypothetical protein | |
| fig\|142751 | IgaA | Xenorhabdus szentirmaii DSM 16338 | 1427518,3 | CBXF010000082 | XSR1_240 | VBIXenSz | PATRIC.14 | PATRIC | CDS | 27977 | 30085 | 2109 | + | FIG000040 | PLF_626_ | PGF_0391 | CDL82777 | 702 | umoB | hypothetical protein | |
| fig\|527002 | DnaK | Yersinia aldovae ATCC 35236 | 527002.3 | NZ_ACCB01000018 | yaldo0001_ | VBIYerAld | PATRIC.52 | PATRIC | CDS | 21459 | 23369 | 1911 | + | FIG000233 | PLF_629_ | PGF_1035 | ZP_04620 | 636 | | Chaperone protein DnaK | |
| fig\|527002 | IgaA | Yersinia aldovae ATCC 35236 | 527002,3 | NZ_ACCB01000015 | yaldo0001_ | VBIYerAld | PATRIC.52 | PATRIC | CDS | 18716 | 20854 | 2139 | - | FIG000040 | PLF_629_ | PGF_0001 | ZP_04619 | 712 | | IgaA: a membrane protein that prevents overactivation of the Rcs regulatory system | |
| fig\|527002 | RcsF | Yersinia aldovae ATCC 35236 | 527002,3 | NZ_ACCB01000035 | | VBIYerAld | PATRIC.52 | PATRIC | CDS | 3797 | 4153 | 357 | + | FIG013040 | PLF_629_ | PGF_00037945 | | 118 | | Protein RcsF | |
| fig\|527002 | RcsC | Yersinia aldovae ATCC 35236 | 527002,3 | NZ_ACCB01000043 | yaldo0001_ | VBIYerAld | PATRIC.52 | PATRIC | CDS | 14376 | 17243 | 2868 | + | FIG000042 | PLF_629_ | PGF_0005 | ZP_04621 | 955 | | Sensor hist | GO:0004673\|protein histidine kinase activity |
| fig\|527002 | RcsB | Yersinia aldovae ATCC 35236 | 527002,3 | NZ_ACCB01000043 | yaldo0001_ | VBIYerAld | PATRIC.52 | PATRIC | CDS | 17474 | 18127 | 654 | - | FIG000041 | PLF_629_ | PGF_0042 | ZP_04621 | 217 | | DNA-binding capsular synthesis response regulator RcsB | |

Table-S2

| | | | | | | | | | | | | | | | | | | | | | |
|---|---|---|---|---|---|---|---|---|---|---|---|---|---|---|---|---|---|---|---|---|---|
| fig\|527002 | RcsD | Yersinia aldovae ATCC 35236 | 527002,3 | NZ_ACCB0100004 | yaldo0001_ | VBIYerAld | PATRIC.52 | PATRIC | CDS | 18130 | 20823 | 2694 | - | FIG000045 | PLF_629_ | PGF_0003 | ZP_04621 | 897 | | Phosphotransferase RcsD | |
| fig\|349968 | DnaK | Yersinia bercovieri ATCC 43970 | 349968.5 | NZ_AALC02000002 | yberc0001_ | VBIYerBer | PATRIC.34 | PATRIC | CDS | 46181 | 48094 | 1914 | - | FIG00023 | PLF_629_ | PGF_1035 | ZP_04626 | 637 | | Chaperone protein DnaK | |
| fig\|349968 | RcsF | Yersinia bercovieri ATCC 43970 | 349968,5 | NZ_AALC02000032 | | VBIYerBer | PATRIC.34 | PATRIC | CDS | 41969 | 42325 | 357 | - | FIG01304 | PLF_629_ | PGF_00037945 | | 118 | | Protein RcsF | |
| fig\|349968 | RcsD | Yersinia bercovieri ATCC 43970 | 349968,5 | NZ_AALC02000057 | yberc0001_ | VBIYerBer | PATRIC.34 | PATRIC | CDS | 4319 | 7012 | 2694 | + | FIG000045 | PLF_629_ | PGF_0003 | ZP_04629 | 897 | | Phosphotransferase RcsD | |
| fig\|349968 | RcsB | Yersinia bercovieri ATCC 43970 | 349968,5 | NZ_AALC02000057 | yberc0001_ | VBIYerBer | PATRIC.34 | PATRIC | CDS | 7015 | 7671 | 657 | + | FIG000041 | PLF_629_ | PGF_0042 | ZP_04629 | 218 | | DNA-binding capsular synthesis response regulator RcsB | |
| fig\|349968 | RcsC | Yersinia bercovieri ATCC 43970 | 349968,5 | NZ_AALC02000057 | yberc0001_ | VBIYerBer | PATRIC.34 | PATRIC | CDS | 7870 | 10740 | 2871 | - | FIG000042 | PLF_629_ | PGF_0005 | ZP_04629 | 956 | | Sensor hist | GO:0004673\|protein histidine kinase activity |
| fig\|349968 | IgaA | Yersinia bercovieri ATCC 43970 | 349968,5 | NZ_AALC02000006 | yberc0001_ | VBIYerBer | PATRIC.34 | PATRIC | CDS | 19230 | 21377 | 2148 | - | FIG00004 | PLF_629_ | PGF_0001 | ZP_04627 | 715 | | IgaA: a membrane protein that prevents overactivation of the Rcs regulatory system | |
| fig\|393305 | DnaK | Yersinia enterocolitica subsp. enterocolitica 8081 | 393305.7 | NC_008800 | YE0609 | VBIYerEnt | PATRIC.39 | PATRIC | CDS | 700187 | 702094 | 1908 | + | FIG00023 | PLF_629_ | PGF_1035 | YP_00100 | 635 | dnaK | Chaperone protein DnaK | |
| fig\|393305 | RcsC | Yersinia enterocolitica subsp. enterocolitica 8081 | 393305,7 | NC_008800 | YE1396 | VBIYerEnt | PATRIC.39 | PATRIC | CDS | 1562600 | 1565467 | 2868 | + | FIG000042 | PLF_629_ | PGF_0005 | YP_00100 | 955 | rcsC | Sensor hist | GO:0004673\|protein histidine kinase activity |
| fig\|393305 | RcsB | Yersinia enterocolitica subsp. enterocolitica 8081 | 393305,7 | NC_008800 | YE1397 | VBIYerEnt | PATRIC.39 | PATRIC | CDS | 1565655 | 1566302 | 648 | - | FIG000041 | PLF_629_ | PGF_0042 | YP_00100 | 215 | rcsB | DNA-binding capsular synthesis response regulator RcsB | |
| fig\|393305 | RcsD | Yersinia enterocolitica subsp. enterocolitica 8081 | 393305,7 | NC_008800 | YE1398 | VBIYerEnt | PATRIC.39 | PATRIC | CDS | 1566305 | 1568998 | 2694 | - | FIG000045 | PLF_629_ | PGF_0003 | YP_00100 | 897 | | Phosphotransferase RcsD | |
| fig\|393305 | RcsF | Yersinia enterocolitica subsp. enterocolitica 8081 | 393305,7 | NC_008800 | YE3256 | VBIYerEnt | PATRIC.39 | PATRIC | CDS | 3556570 | 3556701 | 132 | + | FIG01304 | PLF_629_ | PGF_0003 | YP_00100 | 43 | rcsF | Protein RcsF | |
| fig\|393305 | IgaA | Yersinia enterocolitica subsp. enterocolitica 8081 | 393305,7 | NC_008800 | YE3982 | VBIYerEnt | PATRIC.39 | PATRIC | CDS | 4328156 | 4330303 | 2148 | + | FIG00004 | PLF_629_ | PGF_0001 | YP_00100 | 715 | | IgaA: a membrane protein that prevents overactivation of the Rcs regulatory system | |
| fig\|349966 | DnaK | Yersinia frederiksenii ATCC 33641 | 349966.5 | NZ_AALE02000001 | yfred0001_ | VBIYerFre | PATRIC.34 | PATRIC | CDS | 49075 | 50985 | 1911 | + | FIG00023 | PLF_629_ | PGF_1035 | ZP_04630 | 636 | | Chaperone protein DnaK | |
| fig\|349966 | RcsD | Yersinia frederiksenii ATCC 33641 | 349966,5 | NZ_AALE02000010 | yfred0001_ | VBIYerFre | PATRIC.34 | PATRIC | CDS | 24559 | 27252 | 2694 | + | FIG000045 | PLF_629_ | PGF_0003 | ZP_04632 | 897 | | Phosphotransferase RcsD | |
| fig\|349966 | RcsB | Yersinia frederiksenii ATCC 33641 | 349966,5 | NZ_AALE02000010 | yfred0001_ | VBIYerFre | PATRIC.34 | PATRIC | CDS | 27255 | 27902 | 648 | + | FIG000041 | PLF_629_ | PGF_0042 | ZP_04632 | 215 | | DNA-binding capsular synthesis response regulator RcsB | |
| fig\|349966 | RcsC | Yersinia frederiksenii ATCC 33641 | 349966,5 | NZ_AALE02000010 | yfred0001_ | VBIYerFre | PATRIC.34 | PATRIC | CDS | 28047 | 30896 | 2850 | - | FIG000042 | PLF_629_ | PGF_0005 | ZP_04632 | 949 | | Sensor hist | GO:0004673\|protein histidine kinase activity |
| fig\|349966 | IgaA | Yersinia frederiksenii ATCC 33641 | 349966,5 | NZ_AALE02000012 | yfred0001_ | VBIYerFre | PATRIC.34 | PATRIC | CDS | 83965 | 86109 | 2145 | + | FIG00004 | PLF_629_ | PGF_0001 | ZP_04632 | 714 | | IgaA: a membrane protein that prevents overactivation of the Rcs regulatory system | |
| fig\|349966 | RcsF | Yersinia frederiksenii ATCC 33641 | 349966,5 | NZ_AALE02000003 | | VBIYerFre | PATRIC.34 | PATRIC | CDS | 4031 | 4162 | 132 | + | FIG01304 | PLF_629_ | PGF_00037945 | | 43 | | Protein RcsF | |
| fig\|349965 | DnaK | Yersinia intermedia ATCC 29909 | 349965.6 | NZ_AALF02000045 | yinte0001_ | VBIYerInt6 | PATRIC.34 | PATRIC | CDS | 18711 | 20621 | 1911 | + | FIG00023 | PLF_629_ | PGF_1035 | ZP_04638 | 636 | | Chaperone protein DnaK | |
| fig\|349965 | RcsF | Yersinia intermedia ATCC 29909 | 349965,6 | NZ_AALF02000009 | yinte0001_ | VBIYerInt6 | PATRIC.34 | PATRIC | CDS | 3806 | 4162 | 357 | + | FIG01304 | PLF_629_ | PGF_0003 | ZP_04636 | 118 | | Protein RcsF | |
| fig\|349965 | IgaA | Yersinia intermedia ATCC 29909 | 349965,6 | NZ_AALF02000010 | yinte0001_ | VBIYerInt6 | PATRIC.34 | PATRIC | CDS | 49999 | 52137 | 2139 | - | FIG00004 | PLF_629_ | PGF_0001 | ZP_04636 | 712 | | IgaA: a membrane protein that prevents overactivation of the Rcs regulatory system | |
| fig\|349965 | RcsD | Yersinia intermedia ATCC 29909 | 349965,6 | NZ_AALF02000044 | yinte0001_ | VBIYerInt6 | PATRIC.34 | PATRIC | CDS | 17375 | 20068 | 2694 | + | FIG000045 | PLF_629_ | PGF_0003 | ZP_04638 | 897 | | Phosphotransferase RcsD | |
| fig\|349965 | RcsB | Yersinia intermedia ATCC 29909 | 349965,6 | NZ_AALF02000044 | yinte0001_ | VBIYerInt6 | PATRIC.34 | PATRIC | CDS | 20071 | 20724 | 654 | + | FIG000041 | PLF_629_ | PGF_0042 | ZP_04638 | 217 | | DNA-binding capsular synthesis response regulator RcsB | |
| fig\|349965 | RcsC | Yersinia intermedia ATCC 29909 | 349965,6 | NZ_AALF02000044 | yinte0001_ | VBIYerInt6 | PATRIC.34 | PATRIC | CDS | 21007 | 23805 | 2799 | - | FIG000042 | PLF_629_ | PGF_0005 | ZP_04638 | 932 | | Sensor hist | GO:0004673\|protein histidine kinase activity |
| fig\|527012 | DnaK | Yersinia kristensenii ATCC 33638 | 527012.3 | NZ_ACCA0100003 | ykris0001_ | VBIYerKri1 | PATRIC.52 | PATRIC | CDS | 13017 | 14927 | 1911 | + | FIG00023 | PLF_629_ | PGF_1035 | ZP_04625 | 636 | | Chaperone protein DnaK | |
| fig\|527012 | RcsF | Yersinia kristensenii ATCC 33638 | 527012,3 | NZ_ACCA01000022 | | VBIYerKri1 | PATRIC.52 | PATRIC | CDS | 3947 | 4078 | 132 | + | FIG01304 | PLF_629_ | PGF_00037945 | | 43 | | Protein RcsF | |
| fig\|527012 | IgaA | Yersinia kristensenii ATCC 33638 | 527012,3 | NZ_ACCA01000047 | ykris0001_ | VBIYerKri1 | PATRIC.52 | PATRIC | CDS | 27216 | 29363 | 2148 | - | FIG00004 | PLF_629_ | PGF_0001 | ZP_04626 | 715 | | IgaA: a membrane protein that prevents overactivation of the Rcs regulatory system | |
| fig\|527012 | RcsC | Yersinia kristensenii ATCC 33638 | 527012,3 | NZ_ACCA0100005 | ykris0001_ | VBIYerKri1 | PATRIC.52 | PATRIC | CDS | 28959 | 31757 | 2799 | + | FIG000042 | PLF_629_ | PGF_0005 | ZP_04622 | 932 | | Sensor hist | GO:0004673\|protein histidine kinase activity |
| fig\|527012 | RcsB | Yersinia kristensenii ATCC 33638 | 527012,3 | NZ_ACCA0100005 | ykris0001_ | VBIYerKri1 | PATRIC.52 | PATRIC | CDS | 32060 | 32707 | 648 | - | FIG000041 | PLF_629_ | PGF_0042 | ZP_04622 | 215 | | DNA-binding capsular synthesis response regulator RcsB | |
| fig\|527012 | RcsD | Yersinia kristensenii ATCC 33638 | 527012,3 | NZ_ACCA0100005 | ykris0001_ | VBIYerKri1 | PATRIC.52 | PATRIC | CDS | 32710 | 35403 | 2694 | - | FIG000045 | PLF_629_ | PGF_0003 | ZP_04622 | 897 | | Phosphotransferase RcsD | |
| fig\|349967 | DnaK | Yersinia mollaretii ATCC 43969 | 349967.4 | NZ_AALD02000018 | ymoll0001_ | VBIYerMol | PATRIC.34 | PATRIC | CDS | 49293 | 51203 | 1911 | - | FIG00023 | PLF_629_ | PGF_1035 | ZP_04641 | 636 | | Chaperone protein DnaK | |
| fig\|349967 | RcsF | Yersinia mollaretii ATCC 43969 | 349967,4 | NZ_AALD02000009 | ymoll0001_ | VBIYerMol | PATRIC.34 | PATRIC | CDS | 123791 | 124147 | 357 | - | FIG01304 | PLF_629_ | PGF_0003 | ZP_04640 | 118 | | Protein RcsF | |
| fig\|349967 | IgaA | Yersinia mollaretii ATCC 43969 | 349967,4 | NZ_AALD02000015 | ymoll0001_ | VBIYerMol | PATRIC.34 | PATRIC | CDS | 84089 | 86236 | 2148 | + | FIG00004 | PLF_629_ | PGF_0001 | ZP_04640 | 715 | | IgaA: a membrane protein that prevents overactivation of the Rcs regulatory system | |
| fig\|349967 | RcsD | Yersinia mollaretii ATCC 43969 | 349967,4 | NZ_AALD02000017 | ymoll0001_ | VBIYerMol | PATRIC.34 | PATRIC | CDS | 62880 | 65573 | 2694 | + | FIG000045 | PLF_629_ | PGF_0003 | ZP_04640 | 897 | | Phosphotransferase RcsD | |
| fig\|349967 | RcsB | Yersinia mollaretii ATCC 43969 | 349967,4 | NZ_AALD02000017 | ymoll0001_ | VBIYerMol | PATRIC.34 | PATRIC | CDS | 65576 | 66232 | 657 | + | FIG000041 | PLF_629_ | PGF_0042 | ZP_04640 | 218 | | DNA-binding capsular synthesis response regulator RcsB | |
| fig\|349967 | RcsC | Yersinia mollaretii ATCC 43969 | 349967,4 | NZ_AALD02000017 | ymoll0001_ | VBIYerMol | PATRIC.34 | PATRIC | CDS | 66632 | 69649 | 3018 | - | FIG000042 | PLF_629_ | PGF_0005 | ZP_04640 | 1005 | | Sensor hist | GO:0004673\|protein histidine kinase activity |
| fig\|214092 | DnaK | Yersinia pestis CO92 | 214092.21 | NC_003143 | YPO0468 | VBIYerPes | PATRIC.21 | PATRIC | CDS | 497339 | 499249 | 1911 | + | FIG00023 | PLF_629_ | PGF_1035 | YP_00234 | 636 | dnaK | Chaperone protein DnaK | |
| fig\|214092 | RcsF | Yersinia pestis CO92 | 214092,21 | NC_003143 | YPO1070 | VBIYerPes | PATRIC.21 | PATRIC | CDS | 1213202 | 1213609 | 408 | - | FIG01304 | PLF_629_ | PGF_0003 | YP_00234 | 135 | rcsF | Protein RcsF | |
| fig\|214092 | RcsC | Yersinia pestis CO92 | 214092,21 | NC_003143 | YPO1217 | VBIYerPes | PATRIC.21 | PATRIC | CDS | 1373616 | 1376465 | 2850 | + | FIG000042 | PLF_629_ | PGF_0005 | YP_00234 | 949 | rcsC | Sensor hist | GO:0004673\|protein histidine kinase activity |
| fig\|214092 | RcsB | Yersinia pestis CO92 | 214092,21 | NC_003143 | YPO1218 | VBIYerPes | PATRIC.21 | PATRIC | CDS | 1376533 | 1377186 | 654 | - | FIG000041 | PLF_629_ | PGF_0042 | YP_00234 | 217 | rcsB | DNA-binding capsular synthesis response regulator RcsB | |
| fig\|214092 | RcsD | Yersinia pestis CO92 | 214092,21 | NC_003143 | | VBIYerPes | PATRIC.21 | PATRIC | CDS | 1377189 | 1377761 | 573 | - | FIG000045 | PLF_629_ | PGF_00034034 | | 190 | | Phosphotransferase RcsD | |
| fig\|214092 | RcsD | Yersinia pestis CO92 | 214092,21 | NC_003143 | | VBIYerPes | PATRIC.21 | PATRIC | CDS | 1377929 | 1379806 | 1878 | - | FIG000045 | PLF_629_ | PGF_00034034 | | 625 | | Phosphotransferase RcsD | |
| fig\|214092 | IgaA | Yersinia pestis CO92 | 214092,21 | NC_003143 | YPO0142 | VBIYerPes | PATRIC.21 | PATRIC | CDS | 155018 | 157165 | 2148 | - | FIG00004 | PLF_629_ | PGF_0001 | YP_00234 | 715 | | IgaA: a membrane protein that prevents overactivation of the Rcs regulatory system | |
| fig\|273123 | DnaK | Yersinia pseudotuberculosis IP 32953 | 273123.10 | NC_006155 | YPTB0611 | VBIYerPse | PATRIC.27 | PATRIC | CDS | 724649 | 726559 | 1911 | + | FIG00023 | PLF_629_ | PGF_1035 | YP_06915 | 636 | dnaK | Chaperone protein DnaK | |
| fig\|273123 | RcsC | Yersinia pseudotuberculosis IP 32953 | 273123,1 | NC_006155 | YPTB1257 | VBIYerPse | PATRIC.27 | PATRIC | CDS | 1500619 | 1503468 | 2850 | + | FIG000042 | PLF_629_ | PGF_0005 | YP_06979 | 949 | rcsC | Sensor hist | GO:0004673\|protein histidine kinase activity |
| fig\|273123 | RcsB | Yersinia pseudotuberculosis IP 32953 | 273123,1 | NC_006155 | YPTB1258 | VBIYerPse | PATRIC.27 | PATRIC | CDS | 1503536 | 1504189 | 654 | - | FIG000041 | PLF_629_ | PGF_0042 | YP_06979 | 217 | rcsB | DNA-binding capsular synthesis response regulator RcsB | |
| fig\|273123 | RcsD | Yersinia pseudotuberculosis IP 32953 | 273123,1 | NC_006155 | YPTB1259 | VBIYerPse | PATRIC.27 | PATRIC | CDS | 1504192 | 1506810 | 2619 | - | FIG000045 | PLF_629_ | PGF_0003 | YP_06979 | 872 | | Phosphotransferase RcsD | |
| fig\|273123 | RcsF | Yersinia pseudotuberculosis IP 32953 | 273123,1 | NC_006155 | YPTB2976 | VBIYerPse | PATRIC.27 | PATRIC | CDS | 3514557 | 3514964 | 408 | + | FIG01304 | PLF_629_ | PGF_0003 | YP_07148 | 135 | rcsF | Protein RcsF | |
| fig\|273123 | IgaA | Yersinia pseudotuberculosis IP 32953 | 273123,1 | NC_006155 | YPTB3758 | VBIYerPse | PATRIC.27 | PATRIC | CDS | 4459529 | 4461676 | 2148 | + | FIG00004 | PLF_629_ | PGF_0001 | YP_07223 | 715 | | IgaA: a membrane protein that prevents overactivation of the Rcs regulatory system | |
| fig\|527004 | DnaK | Yersinia rohdei ATCC 43380 | 527004.3 | NZ_ACCD0100002 | yrohd0001 | VBIYerRoh | PATRIC.52 | PATRIC | CDS | 28455 | 30365 | 1911 | + | FIG00023 | PLF_629_ | PGF_1035 | ZP_04613 | 636 | | Chaperone protein DnaK | |
| fig\|527004 | IgaA | Yersinia rohdei ATCC 43380 | 527004,3 | NZ_ACCD01000011 | yrohd0001 | VBIYerRoh | PATRIC.52 | PATRIC | CDS | 39613 | 41736 | 2124 | + | FIG00004 | PLF_629_ | PGF_0001 | ZP_04612 | 707 | | IgaA: a membrane protein that prevents overactivation of the Rcs regulatory system | |
| fig\|527004 | RcsD | Yersinia rohdei ATCC 43380 | 527004,3 | NZ_ACCD0100001 | yrohd0001 | VBIYerRoh | PATRIC.52 | PATRIC | CDS | 1978 | 4671 | 2694 | + | FIG000045 | PLF_629_ | PGF_0003 | ZP_04612 | 897 | | Phosphotransferase RcsD | |
| fig\|527004 | RcsB | Yersinia rohdei ATCC 43380 | 527004,3 | NZ_ACCD0100001 | yrohd0001 | VBIYerRoh | PATRIC.52 | PATRIC | CDS | 4674 | 5321 | 648 | + | FIG000041 | PLF_629_ | PGF_0042 | ZP_04612 | 215 | | DNA-binding capsular synthesis response regulator RcsB | |
| fig\|527004 | RcsC | Yersinia rohdei ATCC 43380 | 527004,3 | NZ_ACCD0100001 | yrohd0001 | VBIYerRoh | PATRIC.52 | PATRIC | CDS | 5524 | 8373 | 2850 | - | FIG000042 | PLF_629_ | PGF_0005 | ZP_04612 | 949 | | Sensor hist | GO:0004673\|protein histidine kinase activity |
| fig\|527004 | RcsF | Yersinia rohdei ATCC 43380 | 527004,3 | NZ_ACCD0100001 | yrohd0001 | VBIYerRoh | PATRIC.52 | PATRIC | CDS | 3680 | 4036 | 357 | + | FIG01304 | PLF_629_ | PGF_0003 | ZP_04612 | 118 | | Protein RcsF | |
| fig\|29486. | DnaK | Yersinia ruckeri strain Big Creek 74 | 29486.45 | CP011078 | UGYR_12890 | | PATRIC.29 | PATRIC | CDS | 2854584 | 2856491 | 1908 | + | FIG00023 | PLF_629_ | PGF_10357457 | | 635 | dnaK | Chaperone protein DnaK | |
| fig\|29486. | RcsF | Yersinia ruckeri strain Big Creek 74 | 29486,45 | CP011078 | UGYR_06560 | | PATRIC.29 | PATRIC | CDS | 1422241 | 1422651 | 411 | + | FIG01304 | PLF_629_ | PGF_00037945 | | 136 | rcsF | Protein RcsF | |
| fig\|29486. | IgaA | Yersinia ruckeri strain Big Creek 74 | 29486,45 | CP011078 | UGYR_11025 | | PATRIC.29 | PATRIC | CDS | 2459272 | 2461416 | 2145 | - | FIG00004 | PLF_629_ | PGF_00013790 | | 714 | | IgaA: a membrane protein that prevents overactivation of the Rcs regulatory system | |
| fig\|29486. | RcsD | Yersinia ruckeri strain Big Creek 74 | 29486,45 | CP011078 | UGYR_04615 | | PATRIC.29 | PATRIC | CDS | 982139 | 984862 | 2724 | + | FIG000045 | PLF_629_ | PGF_00034034 | | 907 | | Phosphotransferase RcsD | |
| fig\|29486. | RcsB | Yersinia ruckeri strain Big Creek 74 | 29486,45 | CP011078 | UGYR_04620 | | PATRIC.29 | PATRIC | CDS | 984864 | 985514 | 651 | + | FIG000041 | PLF_629_ | PGF_00421954 | | 216 | | DNA-binding capsular synthesis response regulator RcsB | |
| fig\|29486. | RcsC | Yersinia ruckeri strain Big Creek 74 | 29486,45 | CP011078 | UGYR_04625 | | PATRIC.29 | PATRIC | CDS | 985609 | 988455 | 2847 | - | FIG000042 | PLF_629_ | PGF_00050577 | | 948 | | Sensor hist | GO:0004673\|protein histidine kinase activity |
| fig\|100236 | DnaK | Yokenella regensburgei ATCC 43003 | 1002368.3 | AGCL01000008 | HMPREF0 | VBIYokReg | PATRIC.10 | PATRIC | CDS | 44410 | 46326 | 1917 | + | FIG00023369 | | PGF_1035 | EHM51166 | 638 | | Chaperone protein DnaK | |
| fig\|100236 | RcsD | Yokenella regensburgei ATCC 43003 | 1002368,3 | AGCL01000047 | HMPREF0 | VBIYokReg | PATRIC.10 | PATRIC | CDS | 1105 | 3759 | 2655 | + | FIG00004592 | | PGF_0003 | EHM44771 | 884 | | Phosphotransferase RcsD | |
| fig\|100236 | RcsB | Yokenella regensburgei ATCC 43003 | 1002368,3 | AGCL01000047 | HMPREF0 | VBIYokReg | PATRIC.10 | PATRIC | CDS | 3776 | 4426 | 651 | + | FIG00004153 | | PGF_0042 | EHM44772 | 216 | | DNA-binding capsular synthesis response regulator RcsB | |
| fig\|100236 | RcsC | Yokenella regensburgei ATCC 43003 | 1002368,3 | AGCL01000047 | HMPREF0 | VBIYokReg | PATRIC.10 | PATRIC | CDS | 4459 | 7377 | 2919 | - | FIG00004208 | | PGF_0005 | EHM44773 | 972 | | Sensor hist | GO:0004673\|protein histidine kinase activity |
| fig\|100236 | RcsF | Yokenella regensburgei ATCC 43003 | 1002368,3 | AGCL01000008 | HMPREF0 | VBIYokReg | PATRIC.10 | PATRIC | CDS | 234977 | 235243 | 267 | - | FIG01304060 | | PGF_0003 | EHM51340 | 88 | | Protein RcsF | |
| fig\|100236 | IgaA | Yokenella regensburgei ATCC 43003 | 1002368,3 | AGCL01000019 | HMPREF0 | VBIYokReg | PATRIC.10 | PATRIC | CDS | 42101 | 44230 | 2130 | + | FIG00004077 | | PGF_0001 | EHM50407 | 709 | | IgaA: a membrane protein that prevents overactivation of the Rcs regulatory system | |

**Table S3.** Genera and species of the order *Enterobacterales* selected for the functional analysis of IgaA orthologs

| name | taxa |
|---|---|
| *Dickeya_dadantii_3937* | *Dickeya* |
| *Dickeya_paradisiaca_NCPPB_2511* | *Dickeya* |
| *Dickeya_zeae_Ech1591* | *Dickeya* |
| *Escherichia_albertii_TW07627* | *Escherichia* |
| *Escherichia_coli_O104_H4_str__2011C_3493* | *Escherichia* |
| *Escherichia_coli_O157_H7_str__Sakai* | *Escherichia* |
| *Escherichia_coli_O83_H1_str__NRG_857C* | *Escherichia* |
| *Escherichia_coli_str__K_12_substr__MG1655* | *Escherichia* |
| *Escherichia_coli_UMN026* | *Escherichia* |
| *Escherichia_fergusonii_ATCC_35469* | *Escherichia* |
| *Escherichia_hermannii_NBRC_105704* | *Escherichia* |
| *Escherichia_vulneris_NBRC_102420* | *Escherichia* |
| *Photorhabdus_asymbiotica_strain_ATCC_43949* | *Photorhabdus* |
| *Photorhabdus_luminescens_subsp__laumondii_TTO1* | *Photorhabdus* |
| *Photorhabdus_temperata_subsp__thracensis_strain_DSM_15199* | *Photorhabdus* |
| *Salmonella_bongori_NCTC_12419* | *Salmonella* |
| *Salmonella_enterica_subsp__arizonae_serovar_62_z4_z23___strain_RSK* | *Salmonella* |
| *Salmonella_enterica_subsp__enterica_serovar_Paratyphi_A_str__ATCC_9* | *Salmonella* |
| *Salmonella_enterica_subsp__enterica_serovar_Typhi_str__CT18* | *Salmonella* |
| *Salmonella_enterica_subsp__enterica_serovar_Typhi_str__Ty2* | *Salmonella* |
| *Salmonella_enterica_subsp__enterica_serovar_Typhimurium_str__14028S* | *Salmonella* |
| *Salmonella_enterica_subsp__enterica_serovar_Typhimurium_str__SL1344* | *Salmonella* |
| *Shigella_boydii_Sb227* | *Shigella* |
| *Shigella_dysenteriae_Sd197* | *Shigella* |
| *Shigella_flexneri_5a_str_M90T* | *Shigella* |
| *Shigella_sonnei_Ss046* | *Shigella* |
| *Sodalis_glossinidius_str___morsitans_* | *Sodalis* |
| *Yersinia_aldovae_ATCC_35236* | *Yersinia* |
| *Yersinia_bercovieri_ATCC_43970* | *Yersinia* |
| *Yersinia_enterocolitica_subsp__enterocolitica_8081* | *Yersinia* |
| *Yersinia_frederiksenii_ATCC_33641* | *Yersinia* |
| *Yersinia_intermedia_ATCC_29909* | *Yersinia* |
| *Yersinia_kristensenii_ATCC_33638* | *Yersinia* |
| *Yersinia_mollaretii_ATCC_43969* | *Yersinia* |
| *Yersinia_pestis_CO92* | *Yersinia* |
| *Yersinia_pseudotuberculosis_IP_32953* | *Yersinia* |
| *Yersinia_rohdei_ATCC_43380* | *Yersinia* |
| *Yersinia_ruckeri_strain_Big_Creek_74* | *Yersinia* |

Strains selected for cloning of *igaA* orthologs

**Table S4.** *S.* Typhimurium/*E. coli* strains and plasmids used in this study

| Bacterial strain/ plasmid | Relevant genotype | Source/ reference |
|---|---|---|
| *S.* Typhimurium | | |
| SL1344 | *hisG64,* virulent strain | (Hoiseth and Stocker, 1981) |
| MD0835 | SL1344 *igaA2*::KXX Δ(*apbE'-rcsC'*) | (Mariscotti and Garcia-Del Portillo, 2008) |
| MD1736 | SL1344 / pBAD24 | This study |
| MD1750 | SL1344 / pBAD24 *igaA*-Myc (*Shigella flexneri)* | This study |
| MD1751 | SL1344 / pBAD24 *igaA*-Myc (*Dickeya dadantii)* | This study |
| MD1752 | SL1344 / pBAD24 *igaA*-Myc (*Sodalis glossinidius)* | This study |
| MD1753 | SL1344 / pBAD24 *igaA*-Myc (*Photorhabdus luminescens)* | This study |
| MD1754 | SL1344 / pBAD24 *igaA*-Myc (*Yersinia enterocolitica)* | This study |
| MD1755 | SL1344 / pBAD24 *igaA*-Myc (*S.* Typhimurium) | This study |
| *E. coli* | | |
| DH10B | $F^-$ *mcrA* Δ(*mrr-hsdRMS-mcrBC*) ϕ80dlacZΔM15 Δ*lacX*74 *deoR recA*1 *endA*1 *araD*139 Δ(*ara*, *leu*)7697 *galU galK* $\lambda^-$ *rpsL nupG* | (Grant et al., 1990) |
| MD1739 | DH10B / pBAD24 | This study |
| MD1742 | DH10B / pBAD24 *igaA*-Myc (*Shigella flexneri*) | This study |
| MD1743 | DH10B / pBAD24 *igaA*-Myc (*Dickeya dadantii*) | This study |
| MD1745 | DH10B / pBAD24 *igaA*-Myc (*Sodalis glossinidius*) | This study |
| MD1747 | DH10B / pBAD24 *igaA*-Myc (*Photorhabdus luminescens*) | This study |
| MD1749 | DH10B / pBAD24 *igaA*-Myc (*Yersinia enterocolitica*) | This study |
| Plasmids | | |
| pGEM-T | $Amp^R$, cloning vector | Lab stock |

| | | |
|---|---|---|
| pBAD24 | Amp$^R$, expression vector (L-arabinose inducible) | National Institute of Genetics, Japan |
| pLR1741 | pGEM-T::*igaA*-Myc (*Shigella flexneri*) | This study |
| pLR1742 | pBAD24::*igaA*-Myc (*Shigella flexneri*) | This study |
| pLR1743 | pBAD24::*igaA*-Myc (*Dickeya dadantii*) | This study |
| pLR1744 | pGEM-T::*igaA*-Myc (*Sodalis glossinidius*) | This study |
| pLR1745 | pBAD24::*igaA*-Myc (*Sodalis glossinidius*) | This study |
| pLR1746 | pGEM-T::*igaA*-Myc (*Photorhabdus luminescens*) | This study |
| pLR1747 | pBAD24::*igaA*-Myc (*Photorhabdus luminescens*) | This study |
| pLR1748 | pGEM-T::*igaA*-Myc (*Yersinia enterocolitica*) | This study |
| pLR1749 | pBAD24::*igaA*-Myc (*Yersinia enterocolitica*) | This study |
| pLR1755 | pBAD24::*igaA*-Myc (*S.* Typhimurium) | This study |

**Table S5.** Primer oligonucleotides used in this study*

| Name | Sequence (5' – 3') | Purpose |
|---|---|---|
| PR-1 | CTA **GCT AGC** AGG AGG AAT TCA CCA TGA GCG TAA CAG TCG TTA TAT TG | cloning *igaA P. luminescens* |
| PR-2 | CCC **AAG CTT** TCA CAG ATC CTC TTC TGA GAT GAG TTT TTG TTG TCG CCA ACG CGT AGG | |
| PR-3 | GTT ACA AAA TAT TCG TAA AGA G | sequence verification *igaA P. luminescens* |
| PR-4 | GAT GGA AGG GAT GGA ATT GCG | |
| SOD-1 | CTA **GCT AGC** AGG AGG AAT TCA CCA TGA GCG CAA TCG TCT ATG TAC TGG C | cloning *igaA S. glossinidius* |
| SOD-2 | CCC **AAG CTT** TCA CAG ATC CTC TTC TGA GAT GAG TTT TTG TTG GTC GCG CCA AAC GCG GTT | |
| SO-3 | GCA ATG AAC AGG TTG AGC TGC | sequence verification *igaA S. glossinidius* |
| SO-4 | CGT CGC TAC CTT GAA CCA GGC | |
| YER-1 | CTA **GCT AGC** AGG AGG AAT TCA CCA TGA GCA CAA TAG TCT TAA TAT TGG CC | cloning *igaA Y. enterolitica* |
| YER-2 | CCC **AAG CTT** TCA CAG ATC CTC TTC TGA GAT GAG TTT TTG TTG ACG TAG GAA AGG CGC TGG | |
| YER-3 | CAA ACG ATC CCT TTG GTT ATT TCC | sequence verification *igaA Y. enterocolitica* |
| YER-4 | CCT GAC ATT AAT GAC CGG TTC AC | |
| ECH-1 | CTA **GCT AGC** AGG AGG AAT TCA CCA TGC GCA CAA TAC TCA TCA TA | cloning *igaA D. dadantii* |
| ECH-2 | CCC **AAG CTT** TCA CAG ATC CTC TTC TGA GAT GAG TTT TTG TTG TCC GTA CAT GGC GGT G | |
| ERW-3 | CGG GAA ACG CTG GAA GAA C | sequence verification *igaA D. dadantii* |
| ERW-4 | GTT GAG CCT GAT TCT GCT GTT G | |
| COLI-1 | CTA **GCT AGC** AGG AGG AAT TCA CCA TGA GCA CCA TTG TGA TT | cloning *igaA S. flexneri* |
| COLI-2 | CCC **AAG CTT** TCA CAG ATC CTC TTC TGA GAT GAG TTT TTG TTG TTC GAT AAG GCT TTC TGA A | |
| SHI-3 | GGC TAT GCC TTG CAA CCT GTT | sequence verification *igaA S. flexneri* |
| SHI-4 | GAT ATG CCG CTG AAA TTC ACT | |

| | | |
|---|---|---|
| IgaApB24-1 | CCC **GCT AGC** AGG AGG AAT TCA CCA TGA GCA CCA TTC TGA TTT TTA | cloning *igaA S.* Typhimurium |
| IgaApB24-3 | CCC **AAG CTT** TCA CAG ATC CTC TTC TGA GAT GAG TTT TTG TTG GAT GAG ATT TTC CGG AGA GAG | |
| pBAD-1 | ATC GCA ACT CTC TAC TGT TT | verification insert in pBAD24 |
| pBAD-2 | GAT TTA ATC TGT ATC AGG CTG | |

* In bold italics, sites of restriction enzymes designed for cloning of the PCR product. In blue, sequence encoding the Myc epitope.
